# Discovery of Niclosamide Analogs with Potent Mitochondrial Uncoupling Activity with reduced toxicity

**DOI:** 10.1101/2025.07.04.663252

**Authors:** Haowen Jiang, Alessio Macorano, Enming Xing, Mohamed Jedoui, Shabber Mohammed, Vanessa Lee, Jeffrey Cheng, Lain McDonough, Xiaolin Cheng, Jiangbin Ye, Pui Kai Li

## Abstract

Mitochondrial uncouplers have shown clinical potential across various diseases, including cancer. Niclosamide, an FDA-approved anthelmintic drug, acts as a mild mitochondrial uncoupler and has demonstrated anticancer activity in multiple preclinical cancer models. However, its clinical application remains limited, with some attributing this to poor bioavailability, while the underlying mechanisms are still unclear. Here, we demonstrate that niclosamide exhibits a dose-dependent biphasic effect, promoting uncoupling at low concentration while acting as a mitochondrial inhibitor at high concentration, which could restrict its therapeutic window and limit efficacy. To overcome this challenge, we aimed to develop next-generation mitochondrial uncouplers (MUs) by synthesizing and evaluating novel Niclosamide derivatives with enhanced therapeutic potential. Through structural modifications, we optimized uncoupling activity while reducing inhibitory toxicity, thereby expanding the pharmacological window. Our findings suggest that fine-tuning the molecular structure of mitochondrial uncouplers could provide a safer and more effective metabolic reprogramming strategy for cancer treatment.

---

Mitochondrial uncoupling occurs naturally in various physiological contexts, such as thermogenesis, and is mediated by endogenous uncoupling proteins (UCPs), fatty acids, and hormones^1^. Mitochondrial uncouplers (MUs) are compounds that induce mitochondrial uncoupling by dissipating the proton gradient generated by the electron transport chain (ETC), which normally drives ATP synthesis via ATP synthase ^2–5^. By dissipating the proton gradient, uncouplers eliminate the thermodynamic constraint on electron transport, enabling the ETC to operate at maximal rates as mitochondria strive to restore the gradient. This elevated electron flux drives increased oxygen consumption, as oxygen serves as the terminal electron acceptor.

Synthetic small-molecule mitochondrial uncouplers have emerged as promising therapeutics for obesity, metabolic syndrome, and aging-related disorders, with growing interest in their potential for cancer treatment ^2, 6–10^. Among these, Niclosamide (Nic) - an antiparasitic drug and mild mitochondrial uncoupler—has garnered significant attention for its potential when repurposed for cancer therapy. ^10–18^. While Nic was initially studied for its inhibition of oncogenic pathways (e.g., Wnt/β-catenin, mTOR, STAT3)^11, 12, 19^, emerging evidence suggests that mitochondrial uncoupling could represent its primary anticancer effector mechanism, potentially driving metabolic and epigenetic remodeling in tumors. For instance, NEN (Nic’s ethanolamine salt) stimulates ETC activity, elevating NAD^+^/NADH to increase pyruvate/lactate (reversing Warburg) and α-KG/2-HG ratios^20^ while suppressing reductive carboxylation^21^. These metabolic reprograming drive epigenetic remodeling via CpG-island demethylation and TET-mediated DNA hydroxymethylation. In neuroblastoma model, these changes promote neuronal differentiation, silence oncogenes (MYCN, β-catenin), and activate tumor suppressors (e.g., p53) while inhibiting HIF signaling^20^.

While suboptimal systemic bioavailability of Nic (plasma concentrations: 0.1–0.72 μM)^12, 22, 23^ has been widely implicated as its primary limitation, however, this rationale fails to explain its limited clinical adoption in colorectal cancer (NCT02687009 and NCT02519582), where the drug directly interacts with intestinal tumors at high local concentrations unconstrained by systemic absorption. This paradox highlights unresolved resistance mechanisms independent of pharmacokinetics, necessitating the development of novel Nic analogs that retain its mitochondrial uncoupling activity while overcoming this unknown limitation.

The uncoupling activity of Nic and its structural analogs involves shuttling protons across the mitochondrial membrane (**Figure 1A**). Upon deprotonation of the phenolic OH, an intramolecular six-membered hydrogen-bonded ring forms between that oxygen and the aniline nitrogen^24^. This H-bonded ring delocalizes the negative charge, therefore stabilizing the anionic form of niclosamide and increase hydrophobicity of the salicylanilide scaffold ∼40-fold as indicated by Storey et al^25^. Such weak acid with a hydrophobic scaffold has the intrinsic ability to carry protons across membrane as both anion A^-^ and neutral form HA can be absorbed in the membrane solution interface. Such capability is closely related to modifications on the salicylanilide core scaffold, which influences the dissociation ability of the phenolic hydroxyl group. Side chain substituents on the scaffold need to be electron-withdrawing to maintain activity^17, 19^, whereas the absence of electron-withdrawing groups or the introduction of electron-donating groups eliminates activity. At concentrations above its uncoupling threshold, niclosamide may inhibit mitochondrial respiration through non-uncoupling mechanisms, similar to FCCP. This effect likely reflects impaired substrate-supported respiration, potentially due to interference with mitochondrial substrate transport across the inner membrane, as direct inhibition of the electron transport chain has been ruled out^26^. This high-dose inhibition critically depends on the strong electron-withdrawing 4-nitro substituent on B-ring^13, 26^ (**Figure 1A**), whose removal abolishes oxygen consumption blockade^26^. Moreover, excessive proton shuttling can trigger opening of the mitochondrial permeability transition pore (mPTP), leading to membrane potential collapse and matrix swelling^7^. We hypothesize that niclosamide exerts a dose-dependent biphasic effect on mitochondrial function, where stimulation at low concentrations and inhibition at higher doses together result in a narrow therapeutic window that limits its clinical applicability. Therefore, optimizing the therapeutic window of niclosamide for anticancer applications requires balancing its mitochondrial uncoupling efficacy with the onset of direct respiratory inhibition at higher doses.

**Figure 1.**
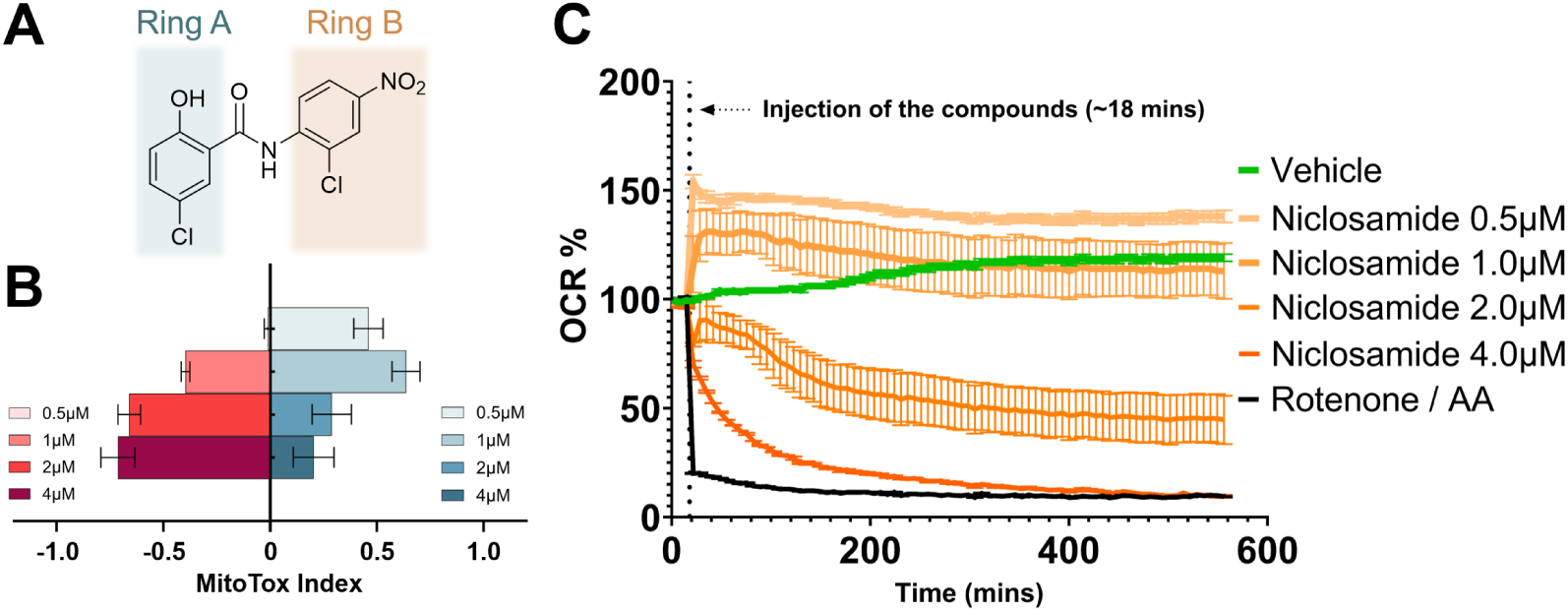
The MitoTox assay and extended kinetic analysis illustrate the uncoupling effects and oxygen consumption ratio (OCR) of Niclosamide. **A.** the structure of Niclosamide, with labeled rings for easy reference. **B.** MitoTox profile of niclosamide. Blue bars represent the uncoupling activity, while the red bars indicate toxicity, as measured by a 45-minute MitoTox assay. **C.** The OCR is assessed using a 9-hour Seahorse assay, where values exceeding 100% suggest sustained uncoupling activity. Conversely, if the OCR falls below 100%, it signals inhibition of respiration.

## Dose-dependent biphasic effect of niclosamide on mitochondrial respiration

To validate our hypothesis, we systematically investigated the dose-dependent effects of Nic on mitochondrial status determined by MitoTox assay. There is an increase in the uncoupling activity as the concentration of Nic increased from 0.5 to 1 μM. However, the uncoupling activity decreases as the concentration increases 2 µM and 4 µM. In addition, there is a corresponding increase in mitochondria inhibition from 2 µM to 4 µM (**Figure 1B**). The longitudinal OCR analysis revealed distinct concentration-dependent trajectories: sustained protonophoric uncoupling persisted throughout the 9-hour assay at 0.5 μM (evidenced by maintained OCR elevation ∼130% baseline), whereas 1 μM elicited transient uncoupling limited to the initial 5-hour phase (OCR peak at 200 min followed by progressive decline to ∼95% baseline). Notably, higher concentrations (2-4 μM) induced rapid mitochondrial suppression, with 4 μM mirroring the complete respiratory inhibition observed in Rotenone/Antimycin A controls (OCR collapse to ∼5% baseline by 400 min), indicative of electron transport chain blockade (**Figure 1C**). These data collectively establish a narrow therapeutic window (0.5-1 μM) for Nic to exert uncoupling activity without triggering mitochondrial respiration failure, beyond which mitochondrial inhibition becomes the dominant phenotype.

## Synthesis and evaluation of the next generation of Niclosamide analogs

To overcome the narrow therapeutic window of Nic and mechanistically dissect how A/B-ring substitutions modulate the uncoupling-inhibition dichotomy, we rationally designed 30 analogs through fine-tuning phenolic hydroxyl (A-ring) and chlorophenyl (B-ring) modifications, followed by systematic therapeutic index quantification via MitoTox profiling (**Figure 2**). Salicylanilide scaffold—the structural core of Nic—functions as a weakly acidic, lipophilic protonophore, shuttling protons across the inner mitochondrial membrane to uncouple oxidative phosphorylation and stimulate ETC activity while dissipating the membrane potential. The ortho-hydroxy group can form an intramolecular hydrogen bond in both its protonated and deprotonated states; hence, fine-tuning the electronic properties of the A- and B-rings offers a route to modulate the hydroxy pKa, proton-binding affinity, and membrane permeability. Accordingly, we synthesized three analogs that preserve the salicylanilide core but vary substitution patterns on both phenyl rings to systematically investigate how these modifications reshape the electronic landscape and biological activity. All niclosamide analogs were synthesized via a one-step amide coupling reaction between the corresponding anilides and the substituted benzoic acids (detailed synthesis procedures can be found in the **supplementary information**). The analogs demonstrated a striking divergence in protonophoric efficacy and mitochondrial safety profiles, establishing a critical structure-dependent dissociation between uncoupling activity and cytotoxic liability.

**Figure 2.**
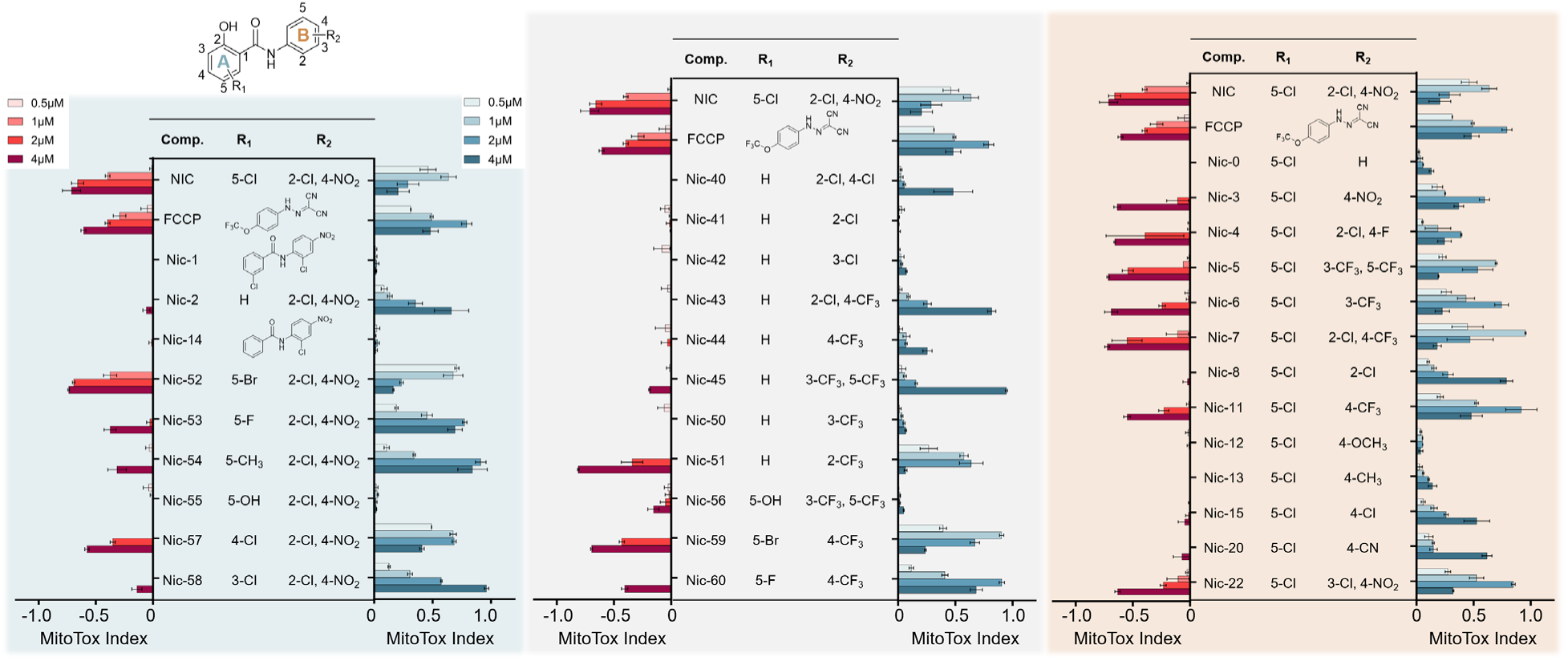
MitoTox profile of synthesized niclosamide analogs. Blue bars indicate uncoupling activity, while red bars represent inhibitory toxicity. Left panel: variations only in the A-ring substituents; right panel: variations only in the B-ring substituents; middle panel: variations in substituents of both rings.

Initial observations established a fundamental prerequisite: analogs devoid of mitochondrial uncoupling activity (e.g., Nic-1, Nic-12, Nic-14, Nic-41, Nic-42, Nic-50, Nic-55, Nic-56) consistently showed no mitochondrial toxicity, confirming the functional linkage between uncoupling activity and inhibitory potential. Significantly, this inhibition dependence manifested clear structural specificity, as evidenced by the divergent safety profiles of Nic-52, Nic-4-6 (high inhibition) versus Nic-2, Nic-8, Nic-40, Nic-43 (low inhibition) despite comparable uncoupling capacities.

Delving into structural determinants, we first dissected A-ring pharmacophores through systematic deconstruction. Removal of the phenolic hydroxyl group (Nic-1, Nic-14) completely abolished uncoupling activity, whereas chlorine substitution (Nic-2 vs parent Nic) proved non-essential. This established the hydroxyl group as the critical hydrogen-bonding motif for proton shuttle formation. Parallel B-ring investigations (Nic-53-58 series) revealed an electronic requirement: only electron-withdrawing substituents sustained uncoupling activity. Halogen positioning and electronic character modulated efficacy gradients, with electron-donating groups (Nic-12/13 methyl/methoxy) completely suppressing mitochondrial uncoupling function.

Therapeutic optimization emerged through combinatorial ring engineering. Maintaining the essential A-ring hydroxyl while tuning B-ring electronics allowed precise control over uncoupling intensity — a parametric relationship validated across hybrid analogs (**Figure 2 middle panel**). Notably, inhibition divergence within active analogs suggested secondary structural determinants beyond basic electronic requirements, possibly involving steric interactions with mitochondrial membranes or off-target binding.

## Dynamic Oxygen Consumption Profiling of Novel Nic Analogs Across Extended Time-Course and Concentration Gradients

To evaluate whether structural modifications to Niclosamide improved mitochondrial selectivity and uncoupling durability, we performed extended oxygen consumption profiling on four lead analogs— Nic-2, Nic-8, Nic-40, and Nic-43—across a 9-hour Seahorse assay (**Figure 3A**). All four compounds demonstrated sustained OCR elevation throughout the entire time course, in contrast to the biphasic trajectory of Niclosamide, which exhibits transient uncoupling followed by delayed mitochondrial inhibition. Notably, Nic-43 and Nic-8 maintained OCR levels exceeding 120% of baseline during the final 100 minutes, reflecting durable protonophoric activity with minimal late-phase toxicity. Quantitative analysis of average OCR between 400–500 minutes further confirmed that these analogs preserved respiratory activity significantly better than Niclosamide (**Figure 3B**). These enhancements are linked to strategic A- and B-ring modifications that maintain the critical A-ring hydroxyl while fine-tuning B-ring electronics to optimize uncoupling efficacy and reduce inhibition (**Figure 3C**). Collectively, these findings highlight the capacity of rational design to decouple uncoupling activity from respiratory suppression and expand the therapeutic window of mitochondrial uncouplers.

**Figure 3.**
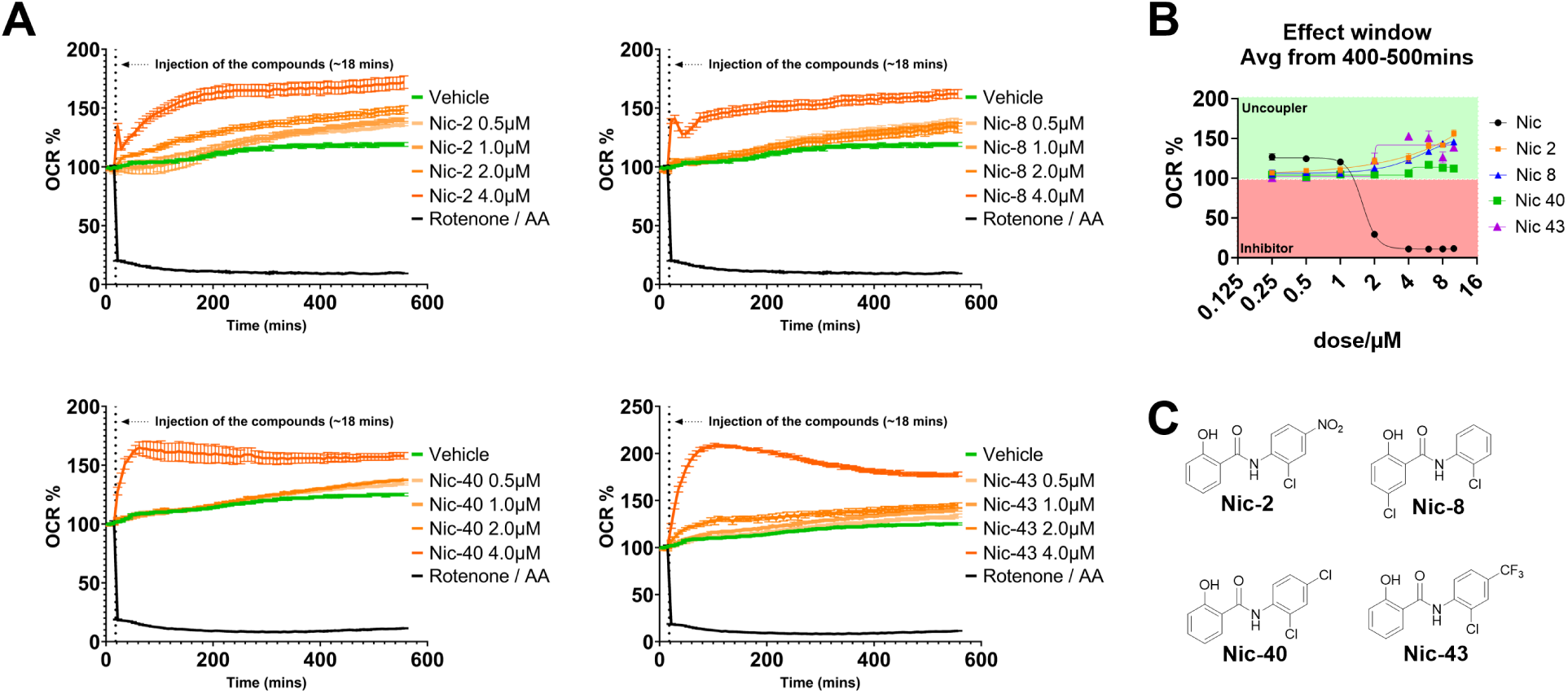
Dynamic Oxygen Consumption Profiling of Novel Nic Analogs. **A.** Nine-hour oxygen consumption kinetics of promising compounds with optimal therapeutic windows: Nic-2, Nic-8, Nic-40, and Nic-43; **B.** The average oxygen consumption over a 400–500 minute window indicates that these compounds demonstrate significantly enhanced uncoupling effects compared to Niclosamide, while exhibiting minimal toxicity; **C.** The chemical structures of Nic-2, Nic-8, Nic-40, and Nic-43.

## QSAR Modeling of Inhibitory and Uncoupling Activities of Niclosamide Analogs

The feature space governing the multidimensional structure-activity landscape of niclosamide analogs is complex and cannot be easily discerned through empirical observation alone, necessitating quantitative modeling. To address this, we developed Quantitative Structure-Activity Relationship (QSAR) models to elucidate the relationships between the inhibitory and uncoupler activities of Nic analogs and their steric and electronic properties. This approach provides a systematic framework to understand how aromatic substituents on the salicylanilide core influence mitochondrial uncoupling versus respiratory inhibition, enabling structure-based predictions to optimize therapeutic efficacy and safety.

We computed a variety of electronic and steric descriptors for the Nic analogs and developed QSAR models using Support Vector Regression (SVR) to correlate these computational chemistry features with MitoTox assay results. Fourteen descriptors were initially considered, including phenolic hydroxyl pKa (pKa), stabilization energy of the anionic form upon deprotonation (De_TOTAL), dipole moment (Dipole), COSMO solvent-excluded surface area (COSMO_AREA), HOMO energy (EHOMO), ring-specific hydrophobicity constants (∑π_A and ∑π_B), and Hammett Sigma Constants (σₘ for *meta* position at R1-4, R2-3, and R2-5; and σₚ for *para* position at R1-5, and R2-4, relative to the anilide nitrogen and phenolic hydroxy as annotated in **Figure 4**)^27, 28^. Due to the influence of steric effect at the *ortho* position, the substituent constant σ₀ cannot be defined as straightforwardly as the *meta* and *para* positions. Previous comprehensive reviews by Tribble et al.^29^ and Charton^30^ investigated σ₀ values for various substituents. In these works, the hydroxyl chemical shifts of 2-substituted phenols and the amine chemical shifts of 2-substituted anilines were employed as surrogate measures for the *ortho* substituent constants σ₀ for position R1-3 relative to phenolic hydroxy and R2-2 relative to anilide nitrogen. To maintain an appropriate feature-to-sample ratio for the dataset (30 compounds), model fitting was constrained to 6 features, resulting in an exhaustive evaluation of 3,003 unique feature combinations (combinatorial selection of 6 out of 14 features, **Figure 5A**).

**Figure. 4.**
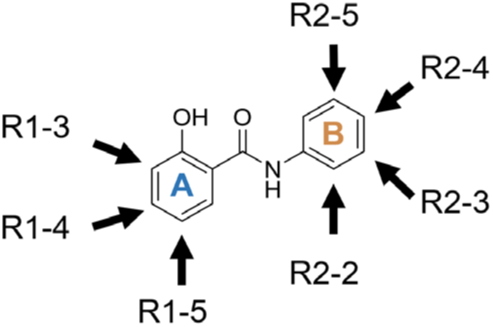
The annotation of positions for assigning substituents’ Hammett inductive parameter.

**Figure 5.**
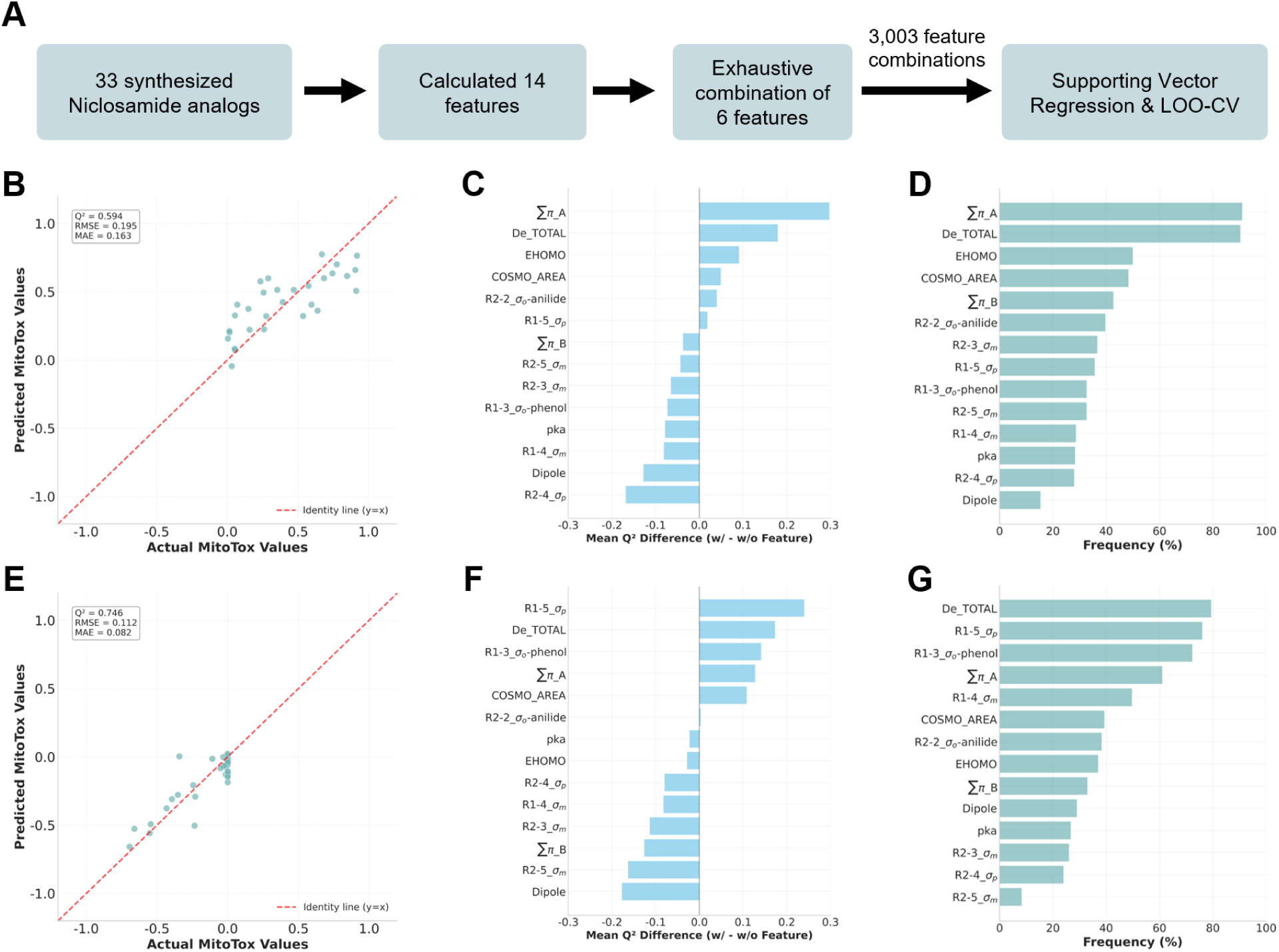
QSAR modeling analysis of niclosamide analogs. **A.** QSAR modeling workflow for all evaluated analogs. **B-D.** Uncoupling activity analysis at 2µM: **B.** predicted versus actual MitoTox profiling values, **C.** feature impact ranking based on mean LOOCV Q² difference, and **D.** feature occurrence frequency among top 10% performance models. **E-G.** Inhibitory activity analysis at 2µM: **E.** predicted versus actual MitoTox profiling values, **F.** feature impact ranking, and **G.** feature occurrence frequency among top 10% performance models.

Model performance exhibited concentration-dependent limitations. At 0.5 µM for both inhibitory and uncoupler activities, and at 1 µM for inhibitory activity, response distributions were largely uniform, rendering them uninformative for QSAR modeling due to minimal activity variation (**Supplement Figure 1**). Similarly, at 4 µM, model performance remained poor across all feature combinations (Q² = 0.407 for uncoupling and 0.587 for inhibition, **Supplement Figure 1)**, with applicability domain analyses provided in **Supplement Figure 2**. These findings suggest that at higher concentrations, non-structure-specific toxicity may dominate assay outcomes, potentially masking structure-activity relationships. Consequently, we focused our modeling efforts on the 2 µM dataset, where moderate structure–activity correlations were observed.

The best-performing models at 2 μM (**Figure 5B, 5E**) yielded Q² values of 0.594 for uncoupling activity and 0.746 for respiratory inhibition under leave-one-out cross-validation (LOOCV). The features used to fit uncoupling activity included pKa, COSMO_AREA, EHOMO, De_TOTAL, ∑π_A, ∑π_B; and those for respiratory inhibition included EHOMO, De_TOTAL, R1-3_σₒ-phenol, R1-5_σₚ, ∑π_A, ∑π_B. These results suggest that the stabilization energy of the deprotonated anionic form, ring-specific hydrophobicity constants, and HOMO energy are key contributors to both respiratory inhibitory and uncoupling activity profiles in the MitoTox assay.

To further assess the importance of each individual feature, we performed a bootstrapping analysis across models of all six-feature combinations. For each feature, we compared the mean Q² values of models with versus without the feature (**Figure 5C, 5F**). The A-ring hydrophobic constant (∑π_A) had a greater influence on uncoupling activity. In contrast, the Hammett Sigma Constants at the *para* and *ortho* positions on the A-ring (R1-5_σₚ and R1-3_σₒ-phenol) played more dominant roles in respiratory inhibition. Interestingly, hydrophobic constant on B-ring (∑π_B) showed much less impact to the model fitting on both activities. The stabilization energy of the deprotonated anionic form (De_TOTAL) showed a comparable impact on both activities. COSMO_AREA slightly favored respiratory inhibition over uncoupling activity, and the HOMO energy (EHOMO) contributed modestly to the fitting of uncoupling activity but had little effect on respiratory inhibition.

We also selected the top 10% of model based on Q² rankings under LOOCV (**Figure 5D, 5G**), and investigated feature occurrence frequency, which corroborated the results from the bootstrapping analysis. Applicability domain analyses (**Supplement Figure 2**) revealed that only Nic-57 and Nic-58 exceeded the S_value threshold of 3. Excluding these descriptors placed all compounds within the applicability domain, confirming that structural divergence arises specifically from local inductive effects.

Collectively, our analyses indicate that the mechanisms of action for niclosamide and its analogs are multifaceted. The distinct sets of features governing uncoupling and inhibitory activities yielded inconsistent predictive performance (**Supplement Figure 1**), likely reflecting the complex, multi-target nature of the niclosamide scaffold. Model performance at higher concentrations (4 μM) was notably inferior compared to that at lower concentrations (i.e., 2 µM), indicating that the compounds may involve more than one mechanism at elevated concentrations. Anion stabilization energy and ring-specific hydrophobicity have been thought to be key factors in predicting MitoTox assay outcomes for Nic analogs. However, our QSAR results revealed minimal differentiation in the descriptor space contributing to uncoupling versus inhibitory activity, suggesting that the chemical space defined by these features may be inherently limited for optimizing therapeutic selectivity and safety of Nic analogs.

## Conclusion

Niclosamide (Nic) is widely recognized as a well-established mitochondrial uncoupler, we demonstrated that mitochondrial uncoupling is a promising approach to inhibit the Warburg effect in cancer cells^31–34^, and while Nic could be an effective tool for inducing uncoupling, its narrow therapeutic window presents a significant challenge. In this work, we successfully showed that an improved therapeutic window can be established to balance pure uncoupling activity and minimize toxicity induced by Nic. By systematically fine-tuning and modifying different substituents on the Nic core scaffold, we identified Nic-2, Nic-8, Nic-40 and Nic-43 as promising candidates with sustained uncoupling activity and minimal toxicity. In the future, these compounds will be further evaluated in both *in vitro* and *in vivo* cancer models to assess their anti-cancer activity, pharmacokinetics, and pharmacodynamics. To further elucidate the structure-activity relationship, we also developed a quantitative structure-activity relationship (QSAR) model to decipher the molecular features that contribute to uncoupling activity and inhibition-related toxicity. Our work lays a solid foundation for utilizing mitochondrial uncouplers to reverse abnormal metabolic states in cancer cells. This approach offers the advantage of reprogramming the metabolic network rather than targeting a specific molecular entity, thereby potentially avoiding the rapid development of drug resistance.

## Supplemental Materials

**Supplement Figure 1.**
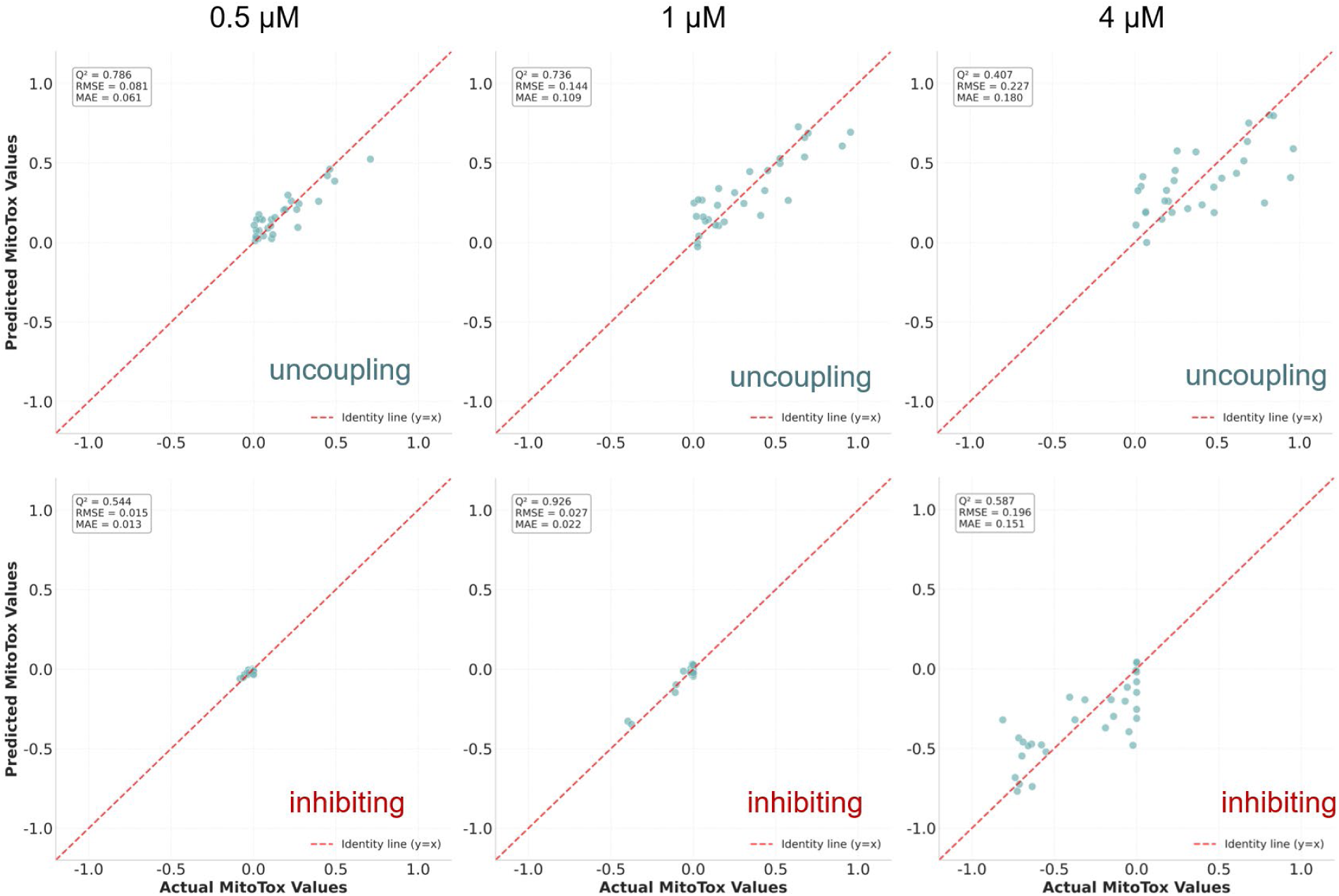
Actual vs. predicted plots for 0.5 µM, 1 µM, and 4 µM. An exhaustive combination of 6 out of 14 features for 0.5 µM, 1 µM, and 4 µM with uncoupling and inhibitory activities were performed, and the predicted versus actual MitoTox profiling values from the best model ranked by Q² scores were plotted. Leave-one-out cross-validation (LOOCV) were performed for each concentration and activity type.

**Supplementary Figure 2.**
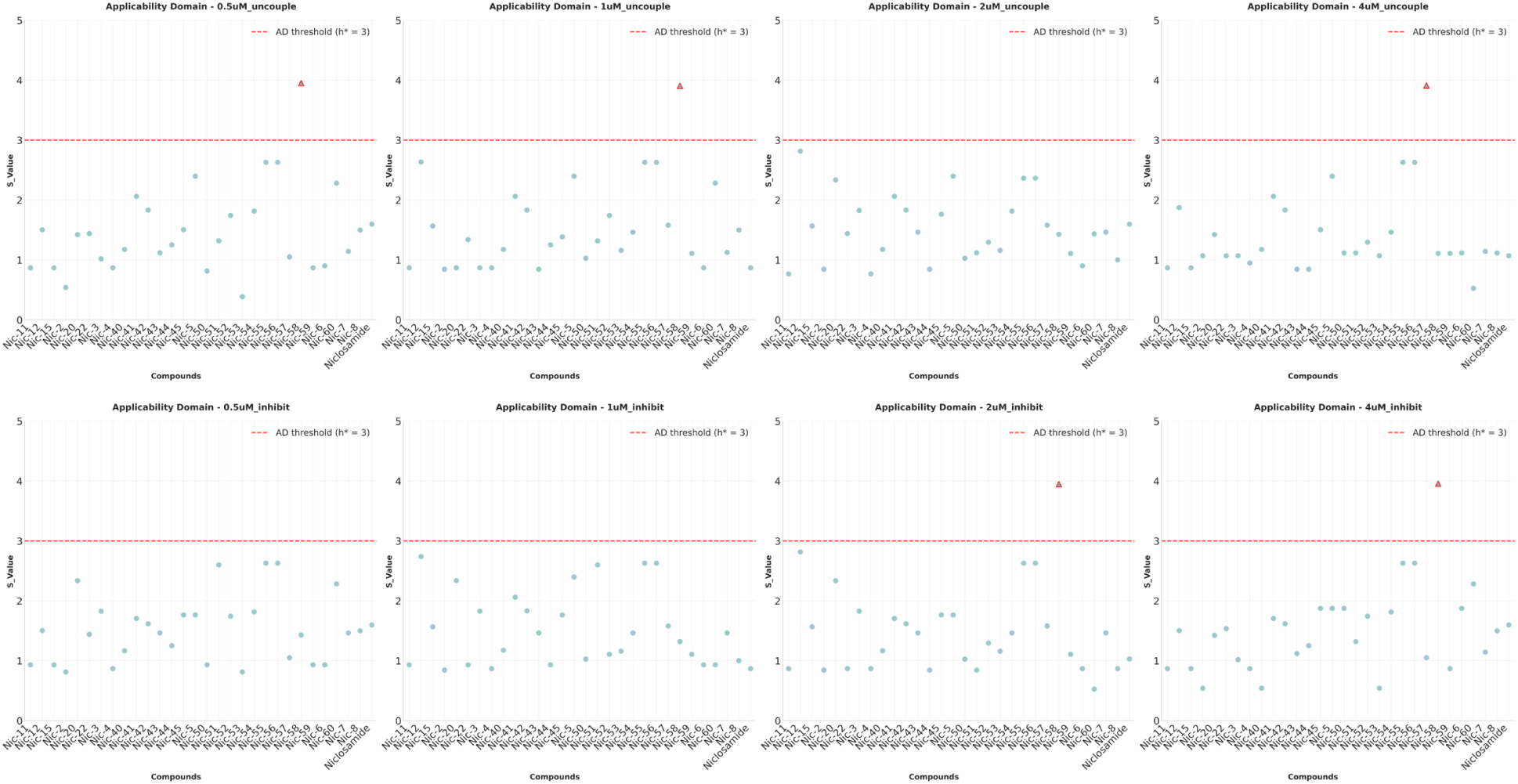
Applicability domain S_value vs. Compound IDs. The applicability domain of the best models obtained at four concentrations, for both uncoupling activity and inhibitory toxicity, using leave-one-out cross-validation (LOOCV) is shown. For each concentration and activity type, scatter plots of the computed S_value versus compound IDs were presented. AD threshold: Applicability Domain threshold.

## Materials and Methods

### Molecular Features Analysis

The following computational tools were used to develop the molecular features used in QSAR model development—Gaussian 16^1^, Alogps^2^, VegaZZ^3^, and Mopac 2016^4^—to derive molecular descriptors for both the protonated (AH) and deprotonated (A^−^) states of each compound. The workflow began with ligand preparation, where we used LigPrep^5^ and Epik^6^ from the Schrödinger Molecular Modelling Suite to generate protonation states within a pH range of 6.7 to 8, as well as relevant tautomers and stereoisomers, optimized under the OPLS3e force field^7^. Conformational searches were conducted using Macromodel^8–10^, applying an OPLS3e force field in a water solvent model with mixed torsional/low-mode sampling. Only conformers with an energy difference of ≤ 5.02 kcal/mol from the lowest-energy conformer were retained. The top 10 conformers for each molecule underwent to structure optimization and frequency calculations using Density Functional Theory (DFT) with the B3LYP/6-31G(d,p) level of theory and an implicit SMD water model. Further refinement included high-level optimization at B3LYP/6-311++G(d,p), followed by single-point energy calculation with MP2/aug-cc-PVDZ, providing a highly accurate electronic energy estimate^11, 12^. The Gibbs free energy was then averaged across conformers using the Boltzmann distribution, providing a weighted calculation of key molecular properties. Additional descriptors were calculated using VegaZZ in combination with the PM7 Hamiltonian (Mopac 2016), providing descriptors relevant to medicinal chemistry.

Specifically, we calculated:

1. pKa: Calculated for the phenolic hydroxyl group, relevant to bioavailability.
2. De_TOTAL: Stabilization energy of the anionic form upon protonation.
3. Dipole: The dipole moment, reflecting charge distribution, is impacted by electron-withdrawing/donating groups.
4. COSMO_AREA: Solvent-excluded surface area to highlight non-polar regions.
5. EHOMO: Energy of the highest occupied molecular orbital, indicating molecular reactivity.

We also assign the Hammett Sigma Constants for substituents R1-3 through R2-5, and the sum of ring-specific hydrophobic constants (∑π_A and ∑π_B)^13–16^.

### QSAR Pipeline

In this study, we used the Scikit-learn package^17^ for QSAR modeling. Specifically, we implemented an exhaustive feature combination workflow to systematically evaluate all possible descriptor combinations. This comprehensive strategy allows the identification of optimal feature subsets without relying on stepwise feature selection methods, which can be prone to local optima. Molecular features were first normalized to a mean of 0 and variance of 1 to ensure equal weighting of descriptors during model development. For model training, we employed Support Vector Regression (SVR) with an RBF kernel (C=10.0), focusing on combinations containing a fixed number of features (typically 5-6) to maintain an appropriate sample-to-feature ratio mitigate the risk of overfitting given our limited dataset size. For model validation, we implemented leave-one-out cross-validation (LOOCV) as it maximizes the size of the training set while still providing an unbiased performance estimate. Model quality was primarily assessed using the cross-validated determination coefficient (Q²), with root-mean-square error (RMSE) and mean absolute error (MAE) as supporting metrics. For each feature combination, we systematically evaluated model performance through cross-validation, stored all evaluation metrics, and identified the combinations yielding the highest predictive accuracy. The exhaustive approach ensures that all possible descriptor combinations are evaluated, avoiding the limitations of greedy optimization algorithms that may miss optimal solutions. For the best-performing feature combinations, comprehensive visualization tools were employed to interpret the models, including actual versus predicted plots, feature importance analyses, and chemical space representations. This rigorous evaluation framework ensures both statistical validity and mechanistic interpretability of the resulting QSAR models. The applicability domain was evaluated using the standardization approach proposed here^18^, in which the S_value were calculated using the same algorithm described in the paper and the corresponding values were plotted against the compound IDs.

### Chemistry

#### Organic Synthesis

Commercial reagents were used as received, without further purification. Anhydrous grade solvents were purchased and utilized directly without additional treatment. Flash column chromatography was performed using a CombiFlash instrument fitted with Flash Pure Buchi columns. Proton nuclear magnetic resonance (¹H NMR) spectra were obtained on a 400 MHz Bruker spectrometer. High-resolution electrospray ionization mass spectrometry (HR-ESI-MS) spectra were recorded on a Thermo Fisher Orbitrap Velos with an autosampler. Low-resolution mass spectrometry was conducted via liquid chromatography–mass spectrometry (LC-MS) on a Waters Acquity UPLC system, using atmospheric pressure chemical ionization (APCI) or electrospray ionization (ESI) as required.

The general synthetic scheme of all the Niclosamide analogs is shown below (**scheme 1**).

#### Scheme 1

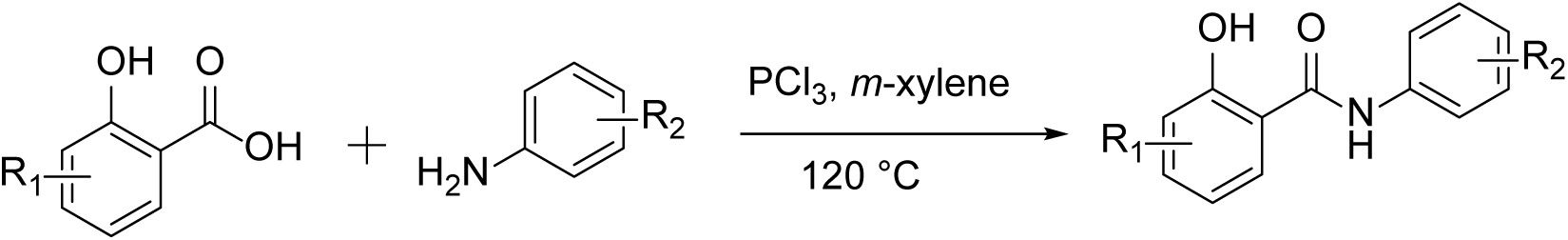

To the solution of corresponding carboxylic acids (1.0 eq) in *m*-xylene was added corresponding anilines (1.1 eq) and heated to 110°C. After 20 min, PCl_3_ (0.4 mmol) was added to reaction mixture and raised the temperature to 120 °C and stirred for another 3 hours. The reaction mixture was brought to 80°C and diluted with water. Major of the compounds are precipitated and filtered off, washed with Hexane and pet-ether and dried. The compounds which are not precipitated, followed the Combi flash purification by using EtOAc in Hexane system.

### 5-chloro-2-hydroxy-N-phenylbenzamide (Nic-0)

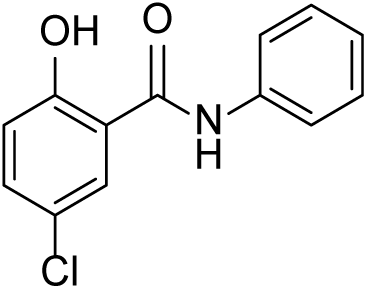

HNMR (300 MHz, DMSO-d6) δ 11.85 (s, 1H), 10.41 (s, 1H), 7.96 (d, *J* = 2.6 Hz, 1H), 7.70 (d, *J* = 7.6 Hz, 2H), 7.47 (dd, *J* = 8.8 Hz, 2.7 Hz, 1H), 7.38 (t, *J* = 7.5 Hz, 2H), 7.15 (t, *J* = 7.3 Hz, 1H), 7.01 (d, *J* = 8.7 Hz, 1H), LC-MS (ESI); [M-H]: 246.0

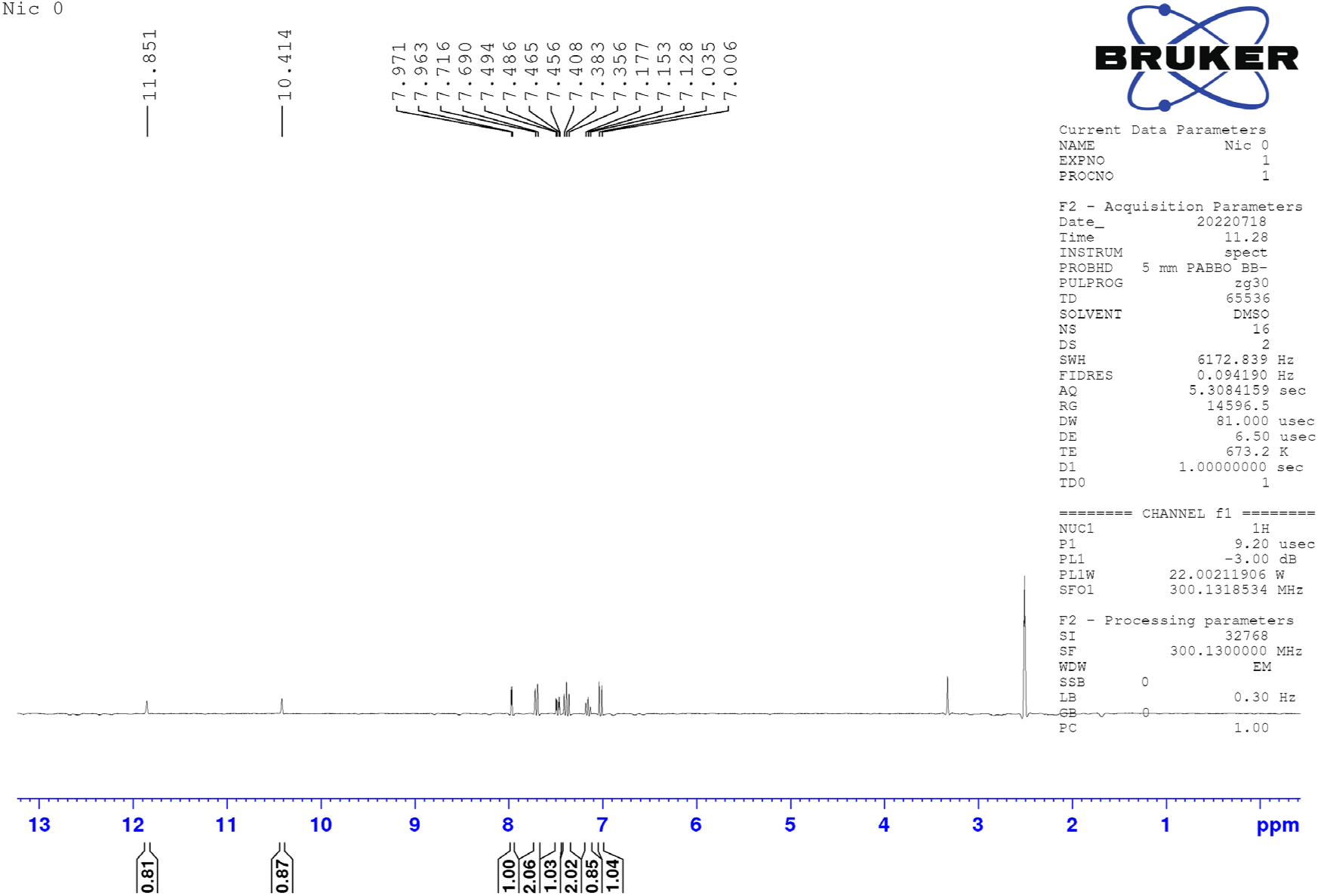

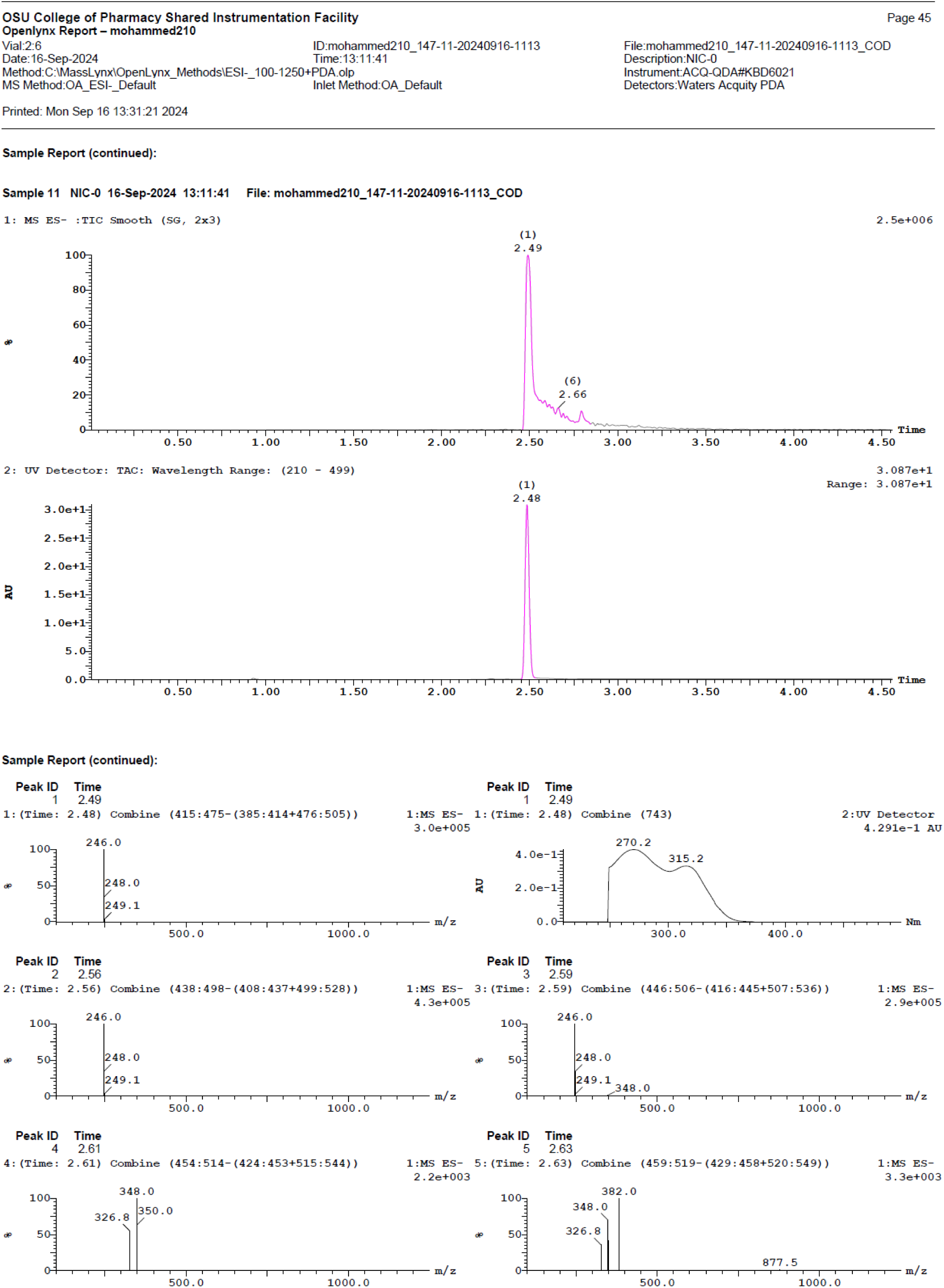

### 3-chloro-N-(2-chloro-4-nitrophenyl)benzamide (Nic-1)

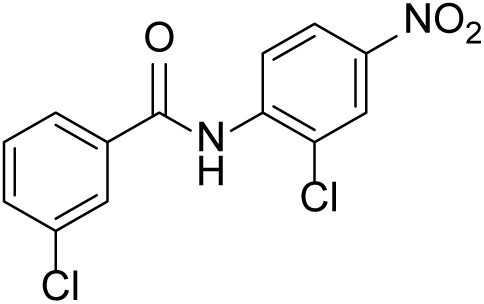

^1^HNMR (400 MHz, DMSO-d_6_) δ 10.46 (s, 1H), 8.41 (d, *J* = 2.5 Hz, 1H), 8.26 (dd, *J* = 8.9 Hz, 2.6 Hz, 1H), 8.04 - 8.0 (m, 2H), 7.96 - 7.94 (m, 1H), 7.74 - 7.71 (m, 1H), 7.61 (t, *J* = 7.8 Hz, 1H); LC-MS (ESI); [M-H]: 308.9

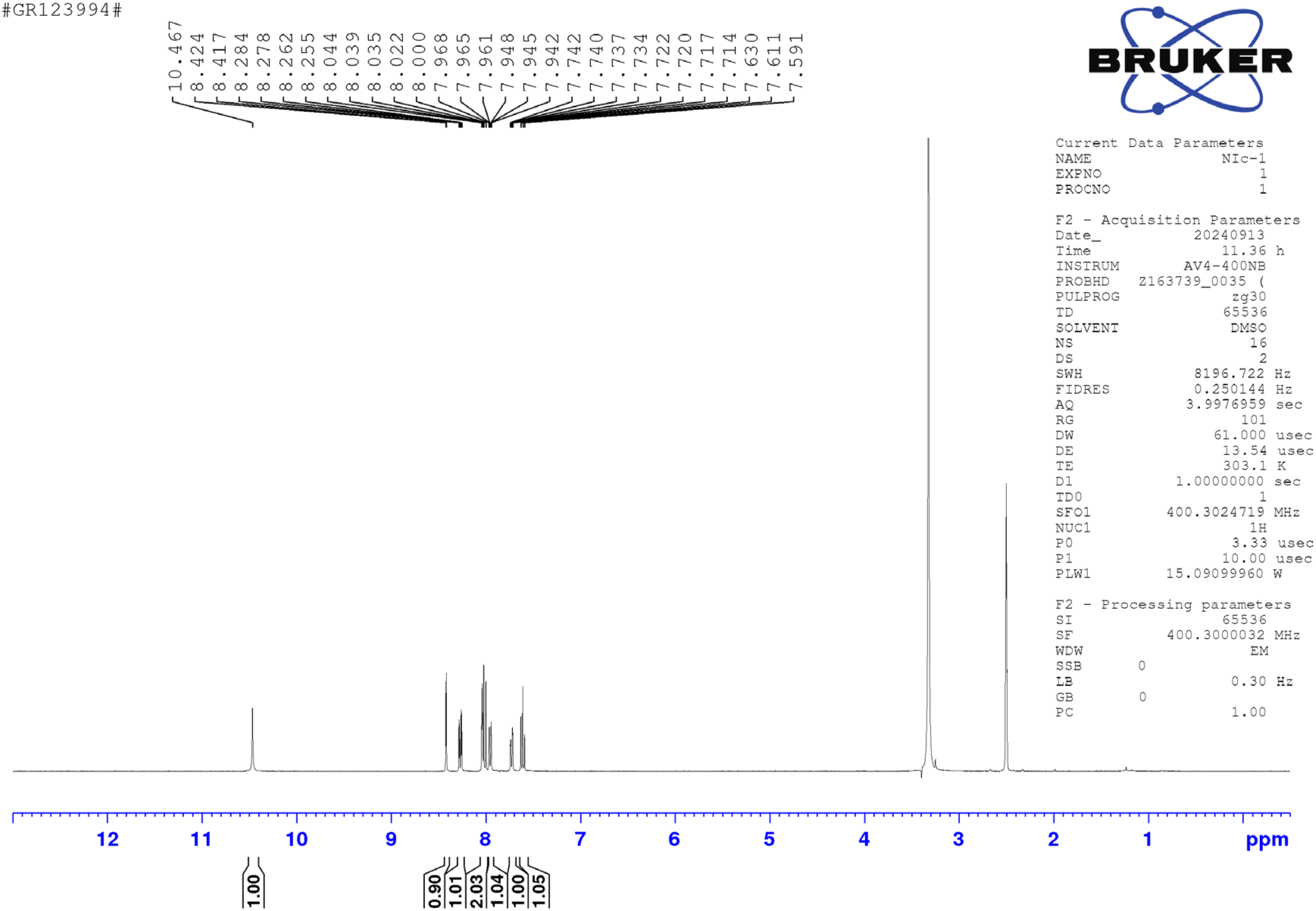

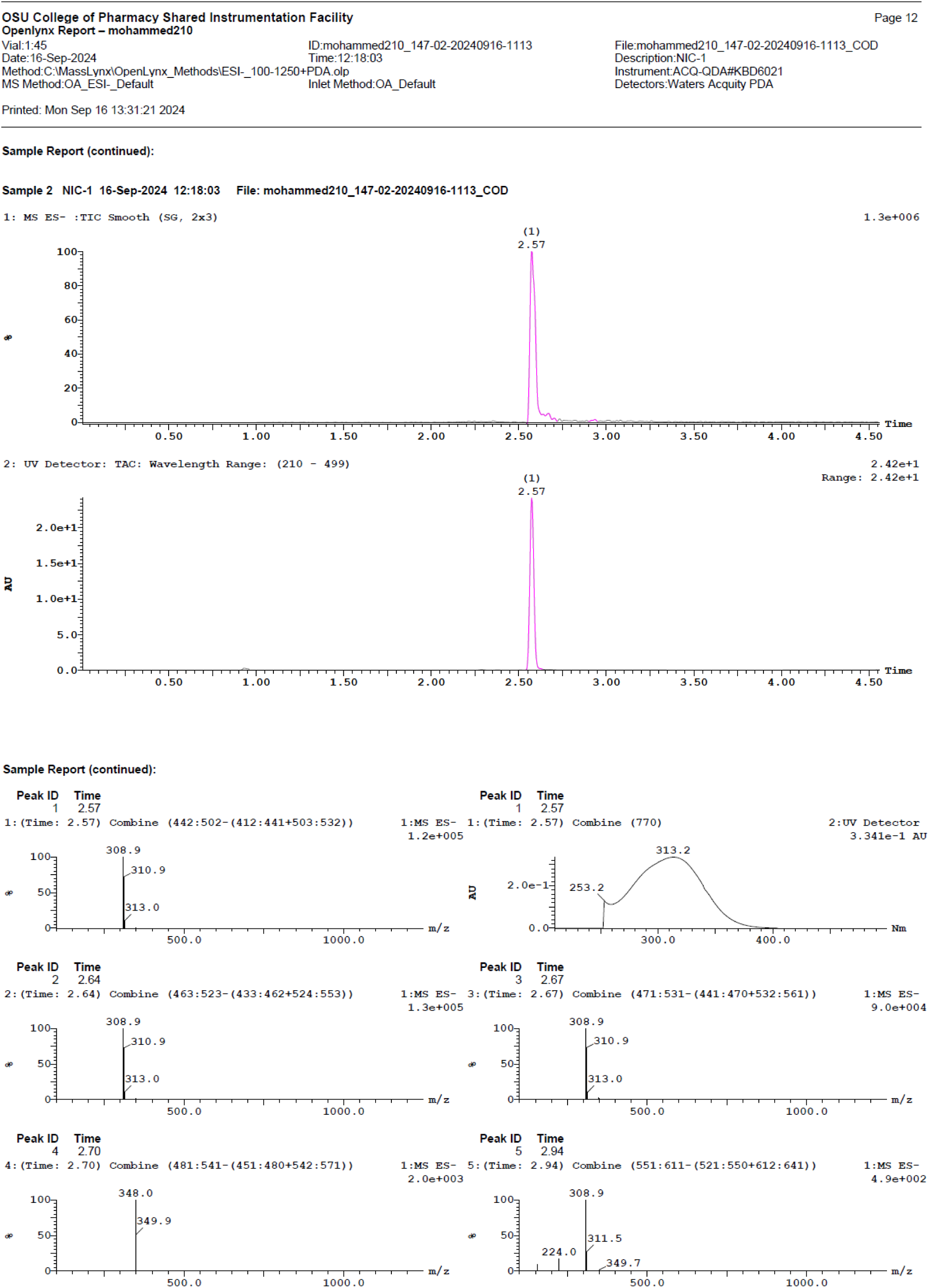

### N-(2-chloro-4-nitrophenyl)-2-hydroxybenzamide (Nic-2)

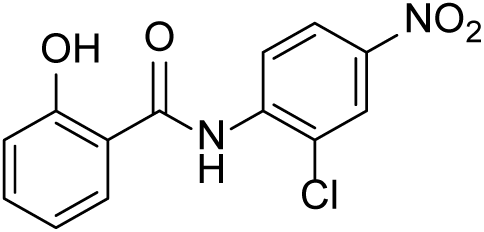

^1^HNMR (400 MHz, DMSO-d_6_) δ 12.12 (s, 1H), 11.37 (s, 1H), 8.83 (d, *J* = 9.2 Hz, 1H), 8.40 (d, *J* = 2.5 Hz, 1H), 8.26 (dd, *J* = 9.2 Hz, 2.5 Hz, 1H), 8.03 (dd, *J* = 7.8 Hz, 1.4 Hz, 1H), 7.48 (td, *J* = 8.4 Hz, 1.5 Hz, 1H), 7.07 - 7.00 (m, 2H); LC-MS (ESI); [M-H]: 290.9

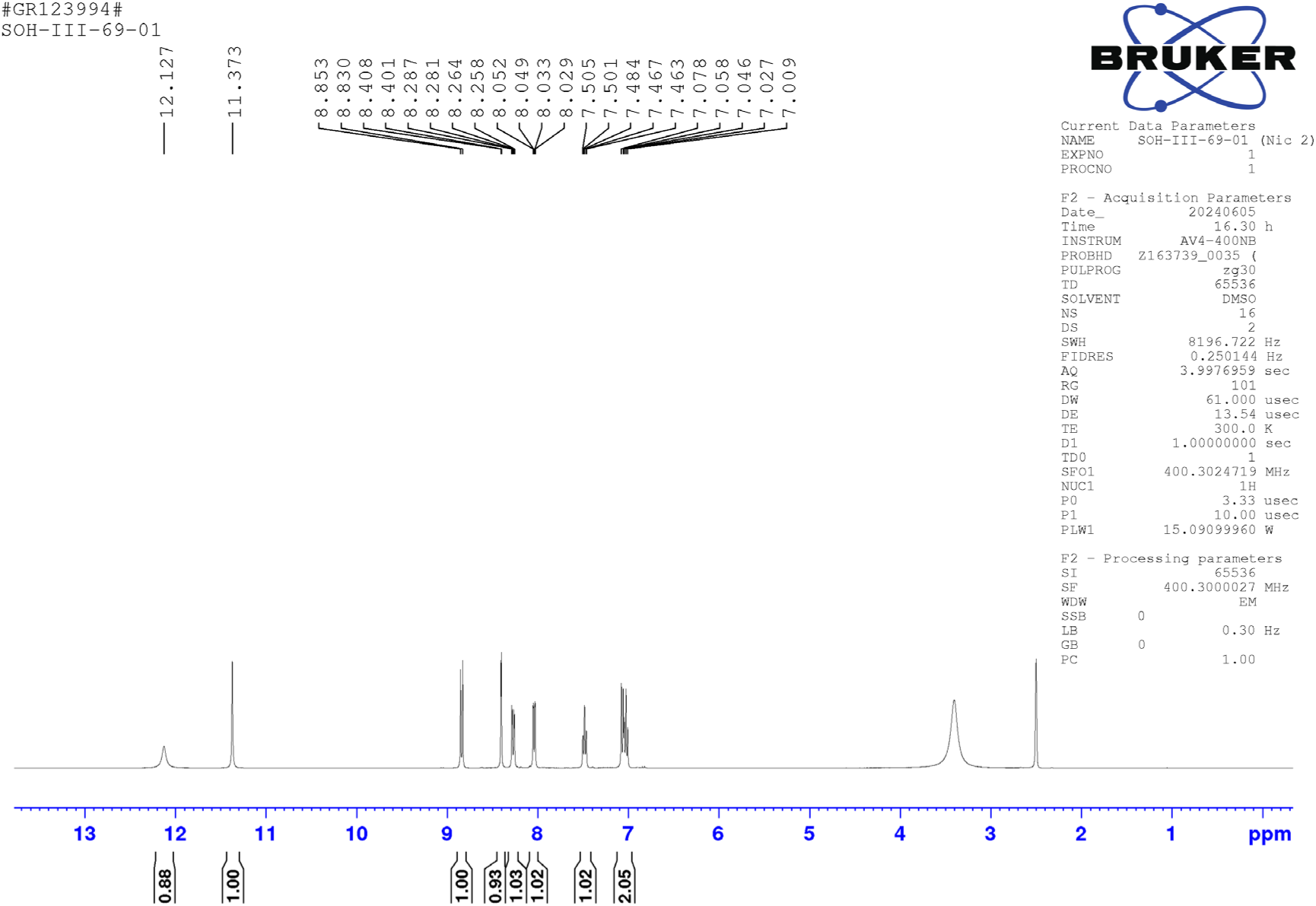

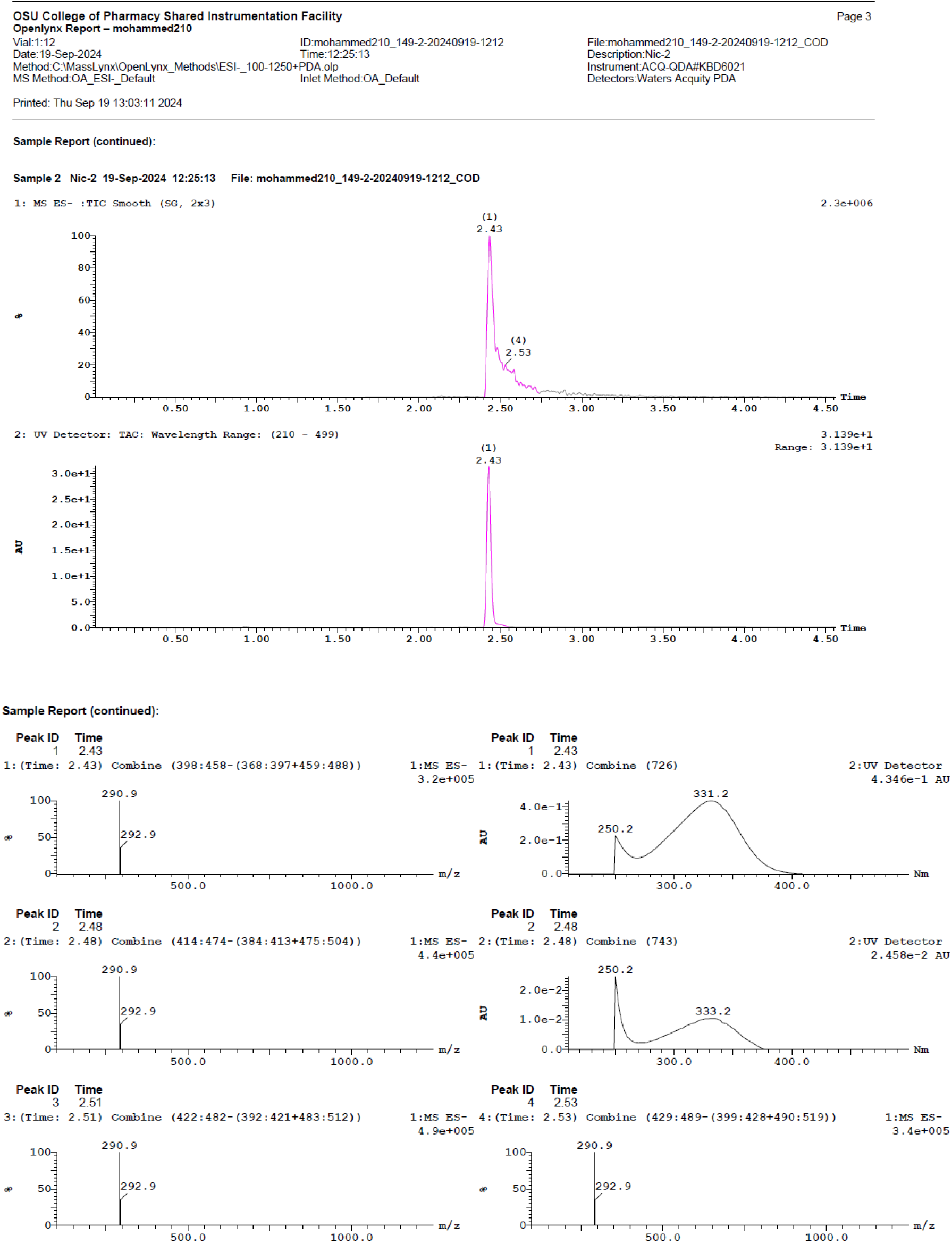

### 5-chloro-2-hydroxy-N-(4-nitrophenyl)benzamide (Nic 3)

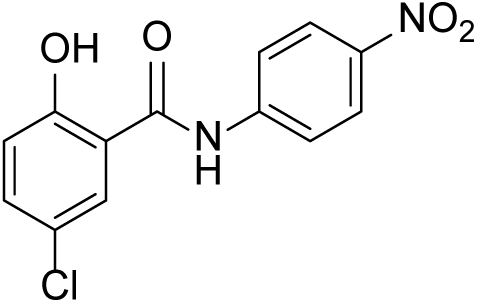

^1^HNMR (400 MHz, CD_3_OD) δ 8.28 (d, *J* = 9.2 Hz, 2H), 8.02 - 7.97 (m, 3H), 7.44 (dd, *J* = 8.8 Hz, 2.6 Hz, 1H), 6.99 (d, *J* = 8.8 Hz, 1H); LC-MS (ESI); [M+H]: 315.6

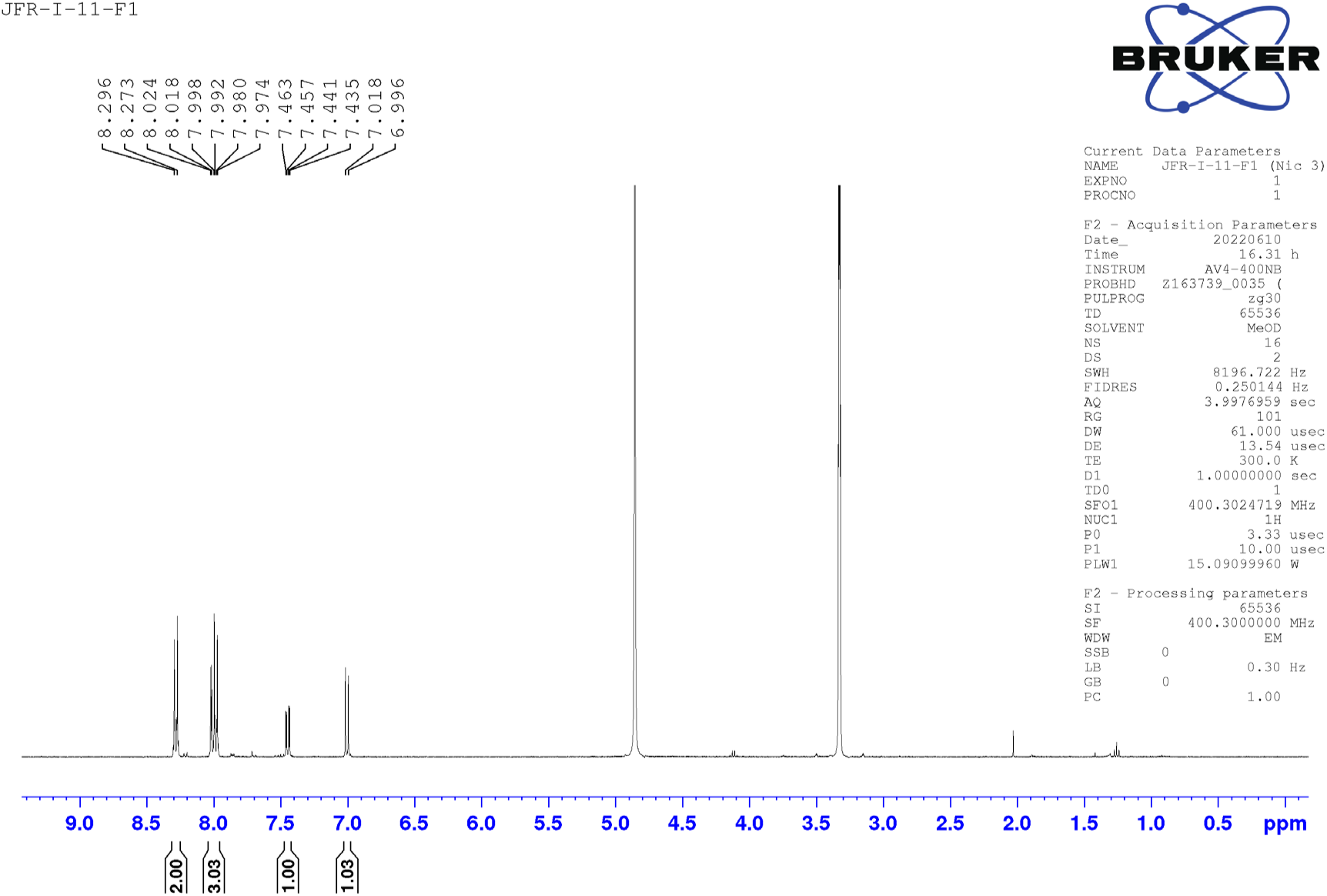

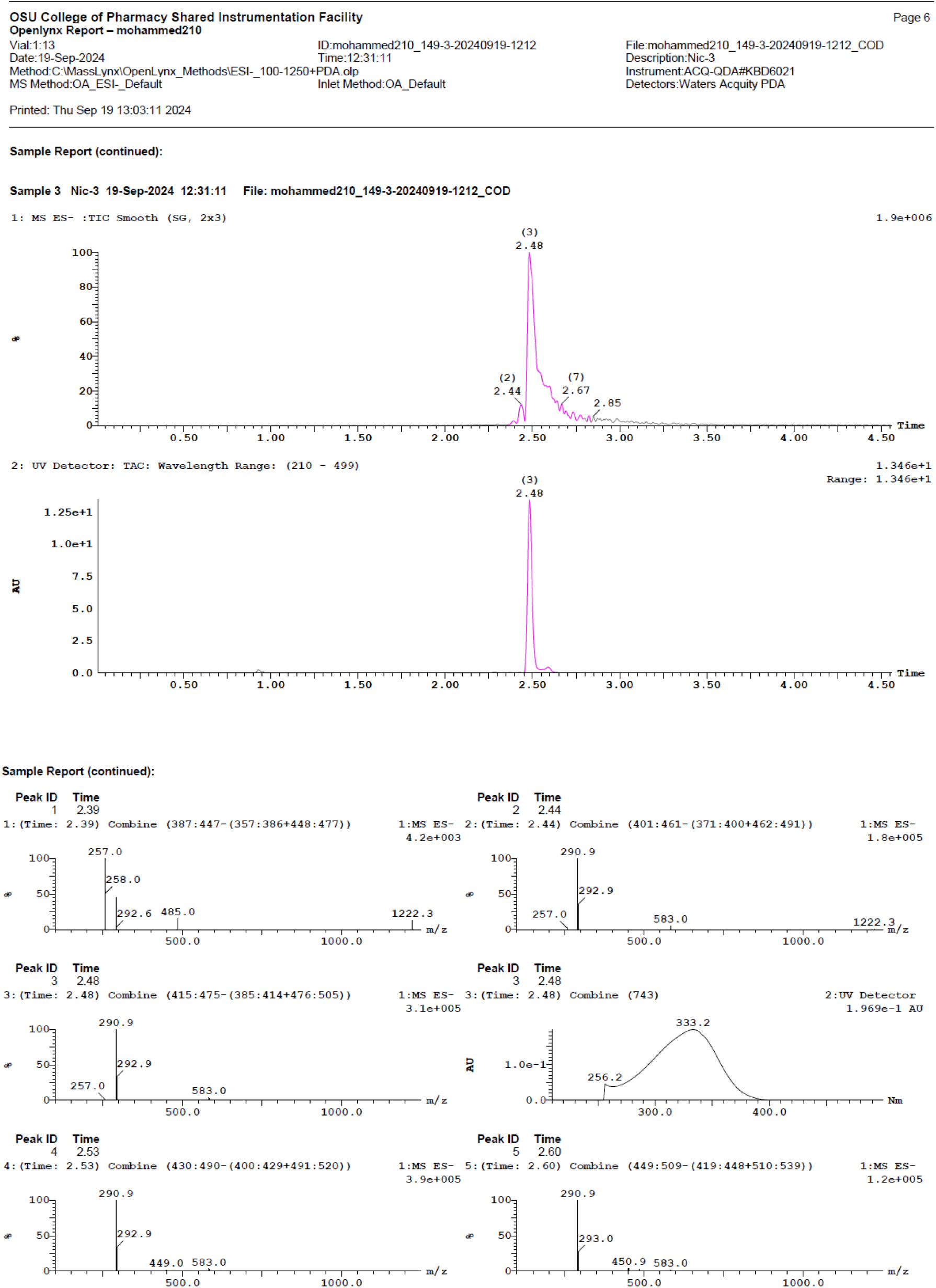

### 5-chloro-N-(2-chloro-4-fluorophenyl)-2-hydroxybenzamide (Nic-4)

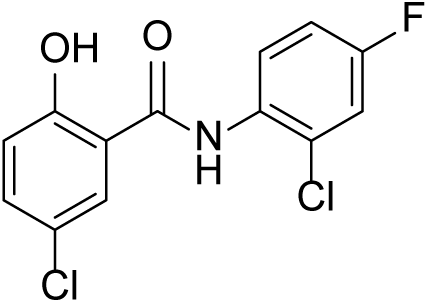

^1^HNMR (400 MHz, DMSO-d_6_) δ 12.22 (s, 1H), 10.78 (s, 1H), 8.32 - 8.28 (m, 1H), 7.98 (d, *J* = 2.8 Hz, 1H), 7.59 -7.49 (m, 2H), 7.32 -7.27(m, 1H), 7.06 (d, *J* = 8.7 Hz, 1H); LC-MS (ESI); [M- H]: 297.9

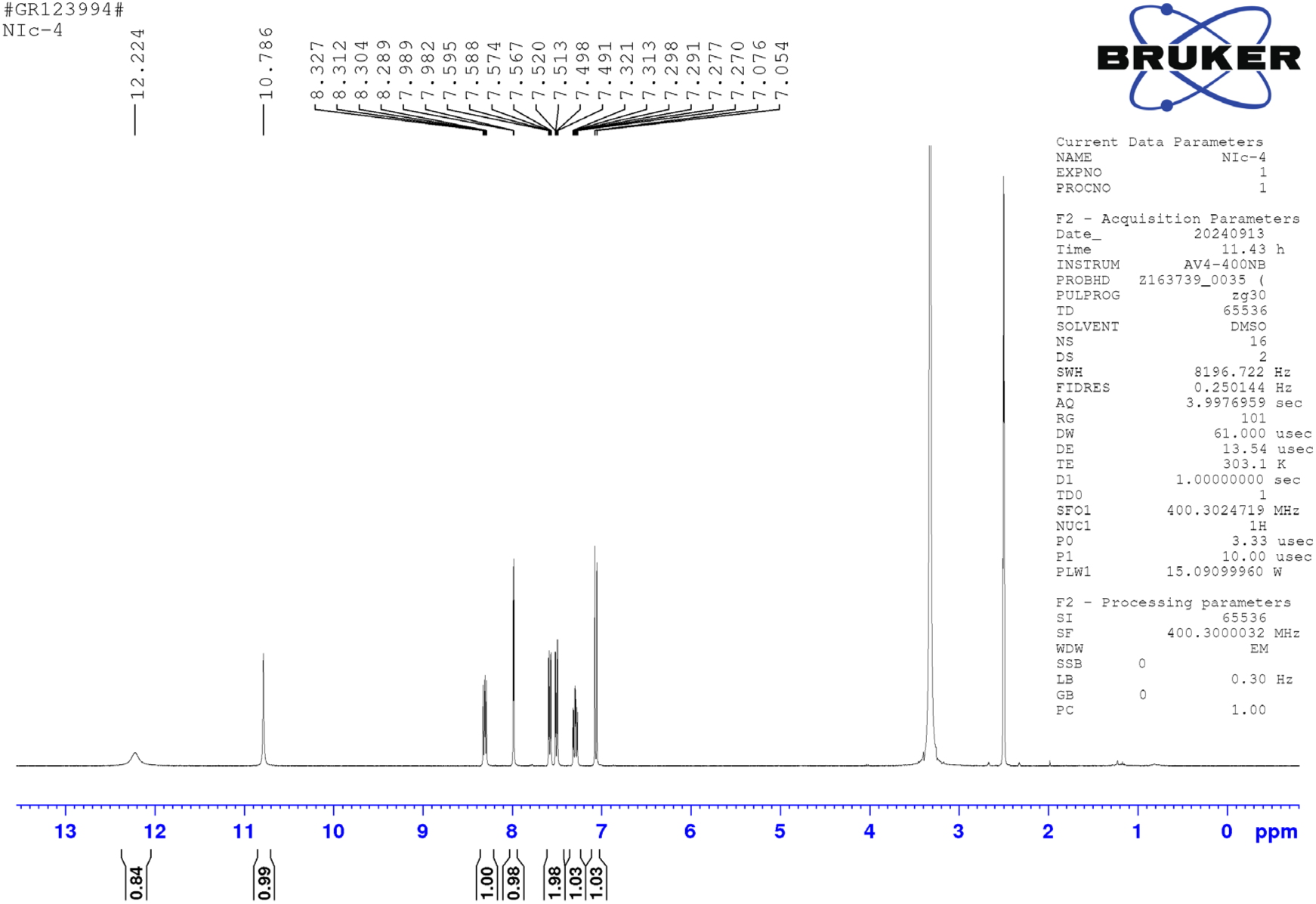

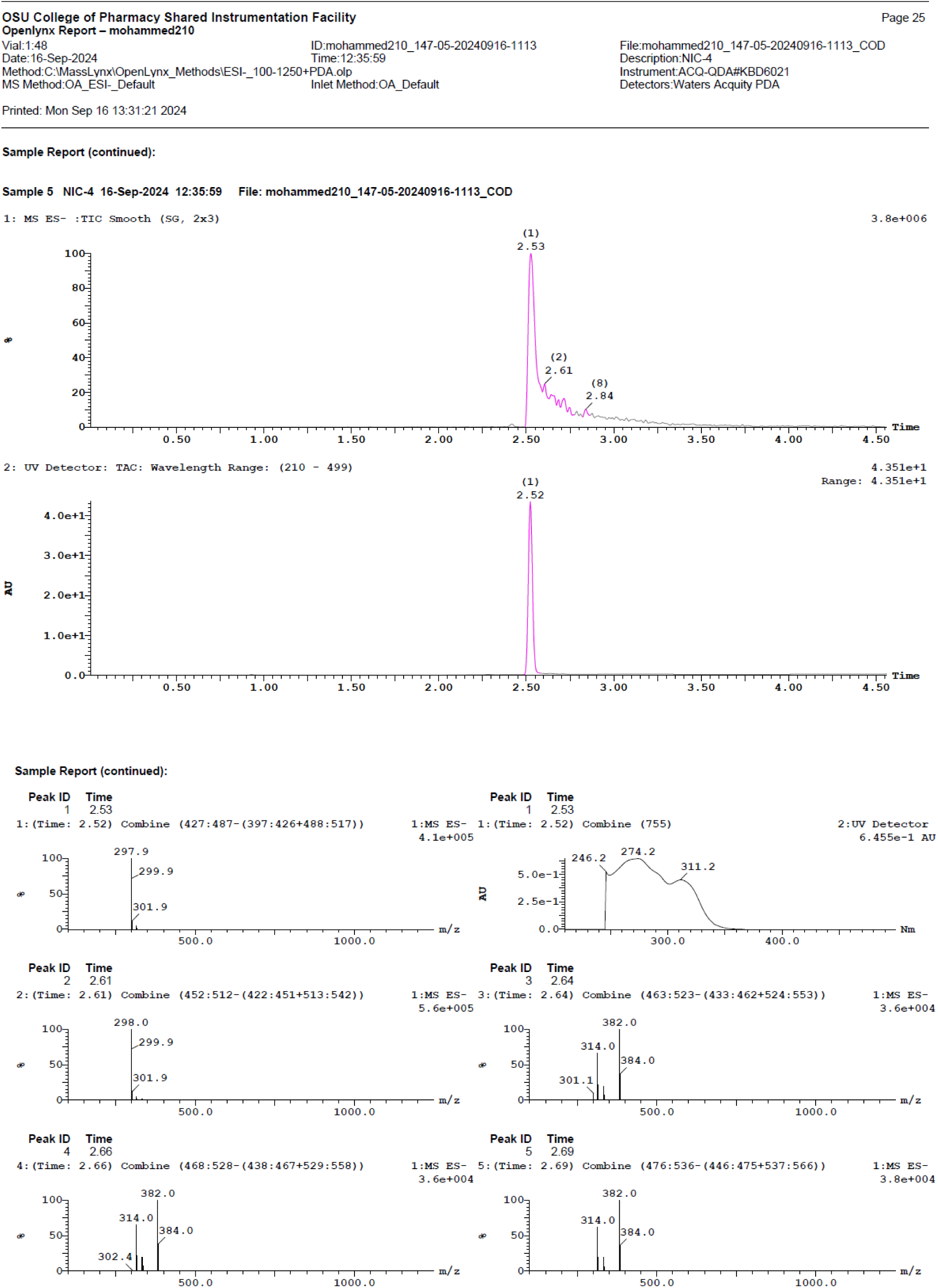

### N-(3,5-bis(trifluoromethyl)phenyl)-5-chloro-2-hydroxybenzamide (Nic 5)

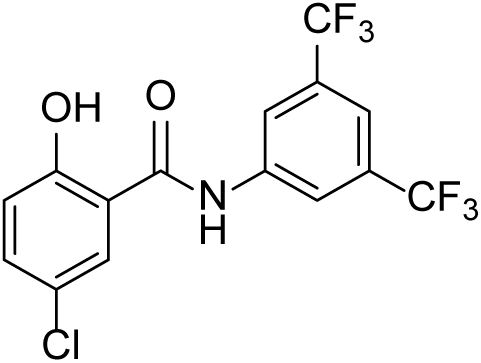

^1^HNMR (400 MHz, CDCl3) δ 11.35 (s, 1H), 8.16 - 8.11 (m, 3H), 7.74 (s, 1H), 7.55 (d, *J* = 2.4 Hz, 1H), 7.47 (dd, *J* = 8.8 Hz, 2.4 Hz, 1H), 7.05 (d, *J* = 8.8 Hz, 1H); LC-MS (ESI); [M-H]: 382.0

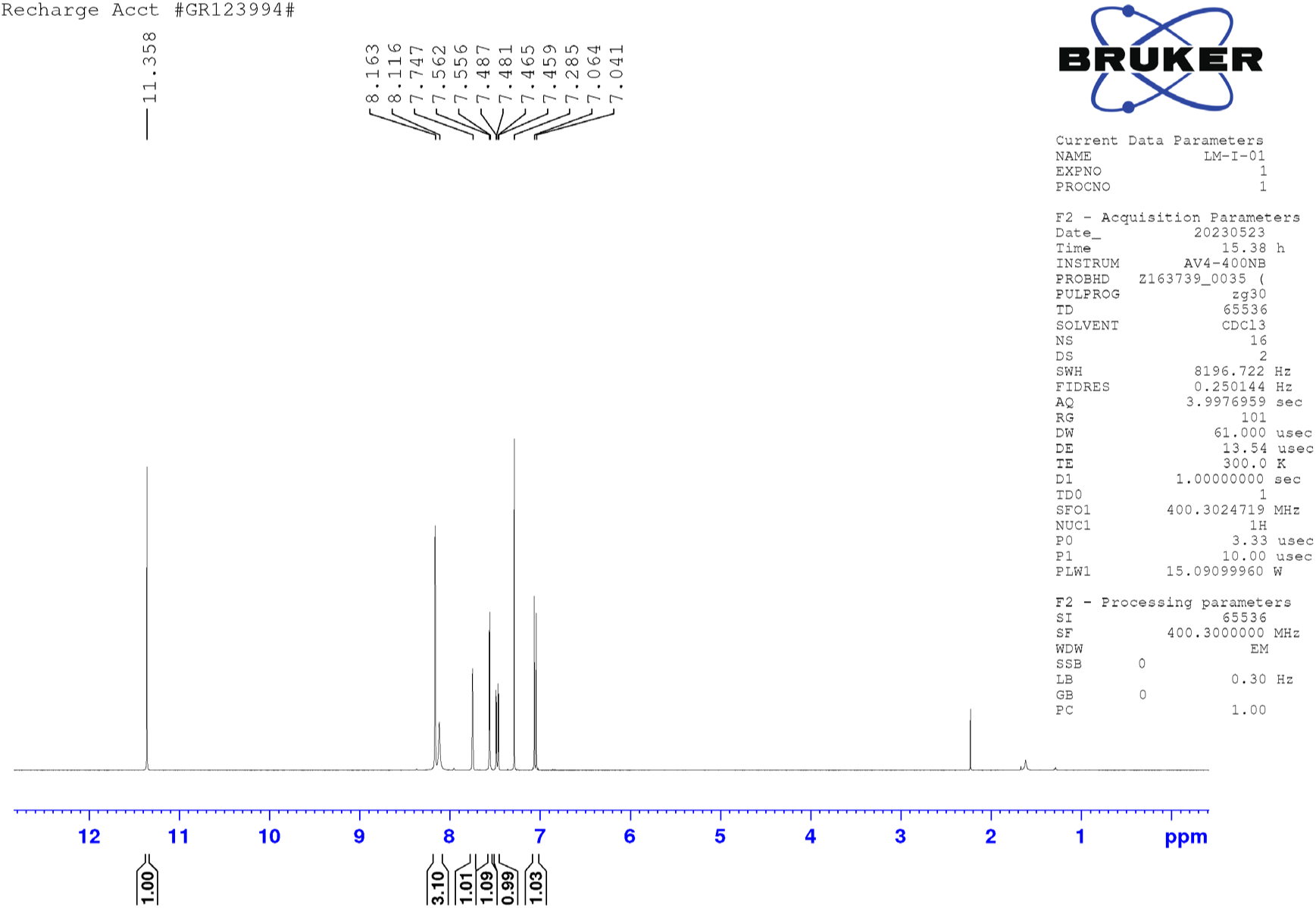

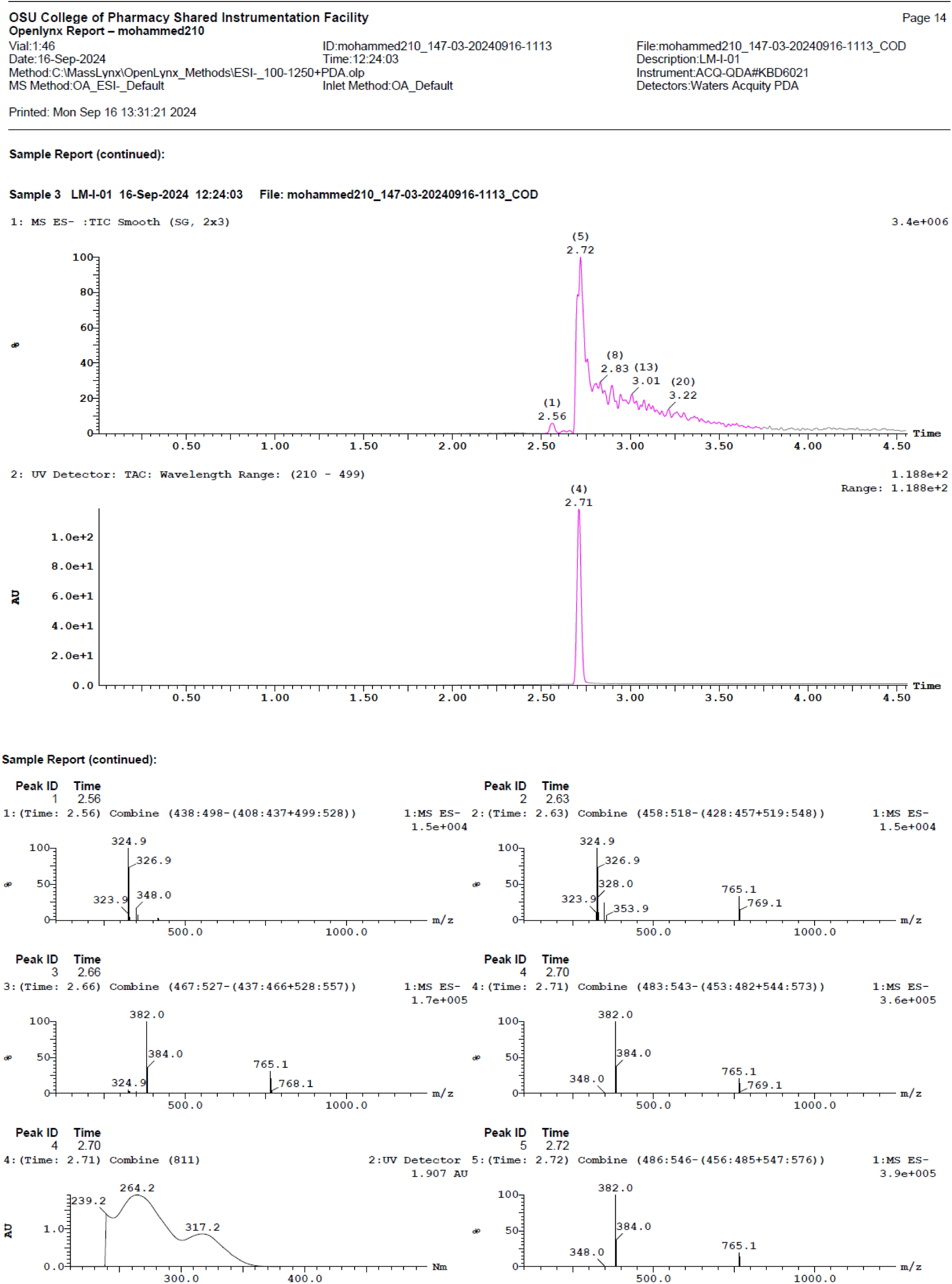

### 5-chloro-2-hydroxy-N-(3-(trifluoromethyl)phenyl)benzamide (Nic-6)

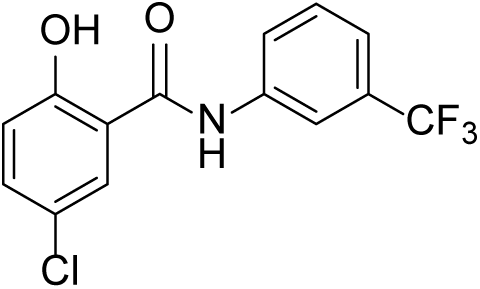

^1^HNMR (400 MHz, DMSO-d_6_) δ 11.62 (s, 1H), 7.91 (bs, 2H), 7.56 - 7.41 (m, 4H), 7.01 (dd, *J* = 8.8 Hz, 1H); LC-MS (ESI); [M-H]: 314.0

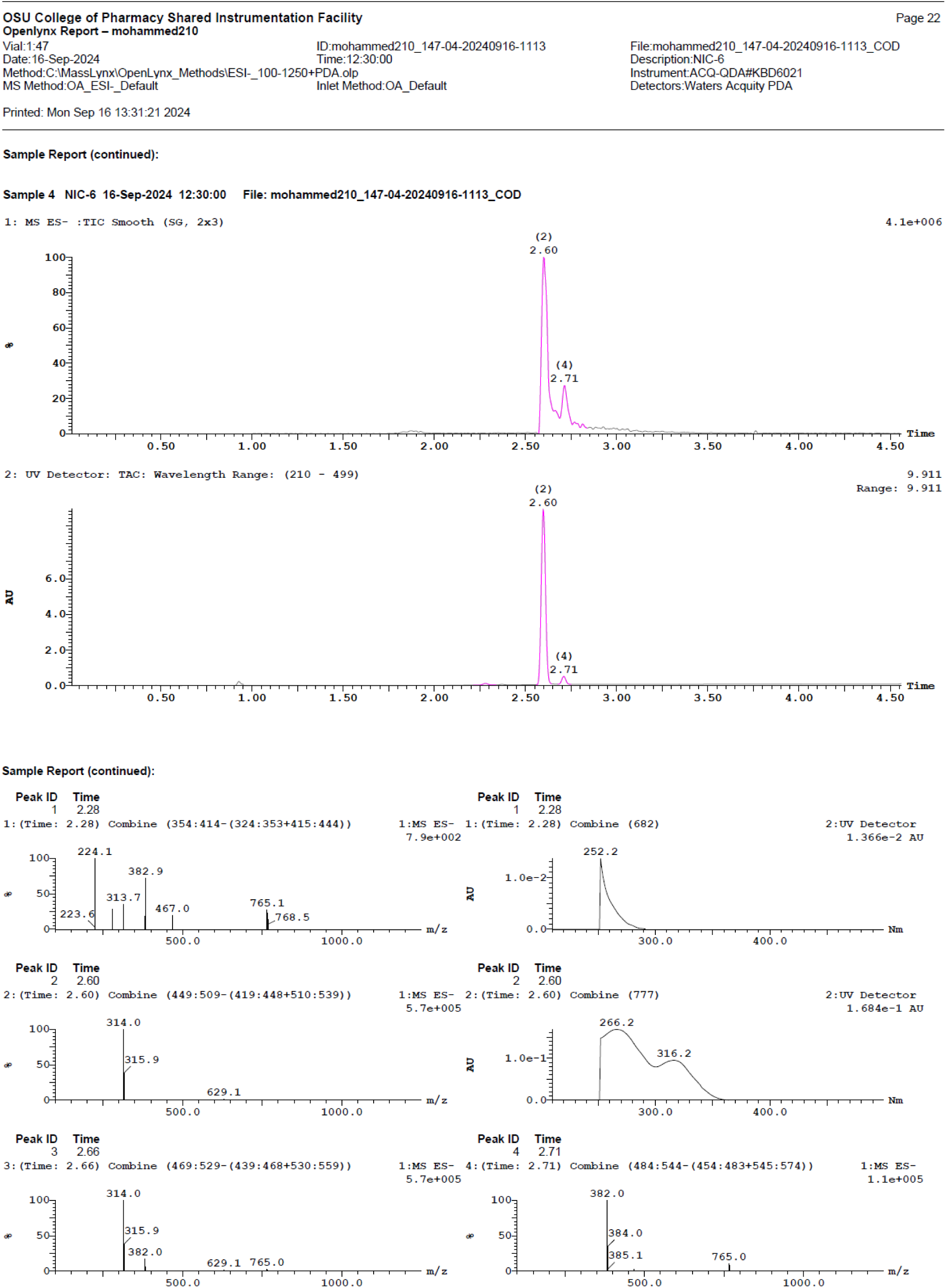

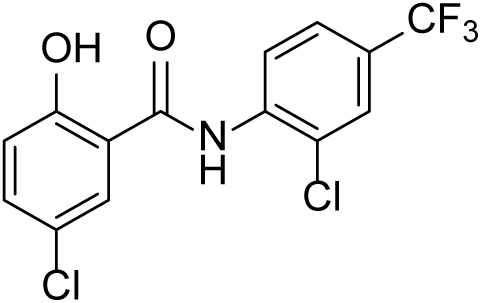

### 5-chloro-N-(2-chloro-4-(trifluoromethyl)phenyl)-2-hydroxybenzamide (Nic-7)

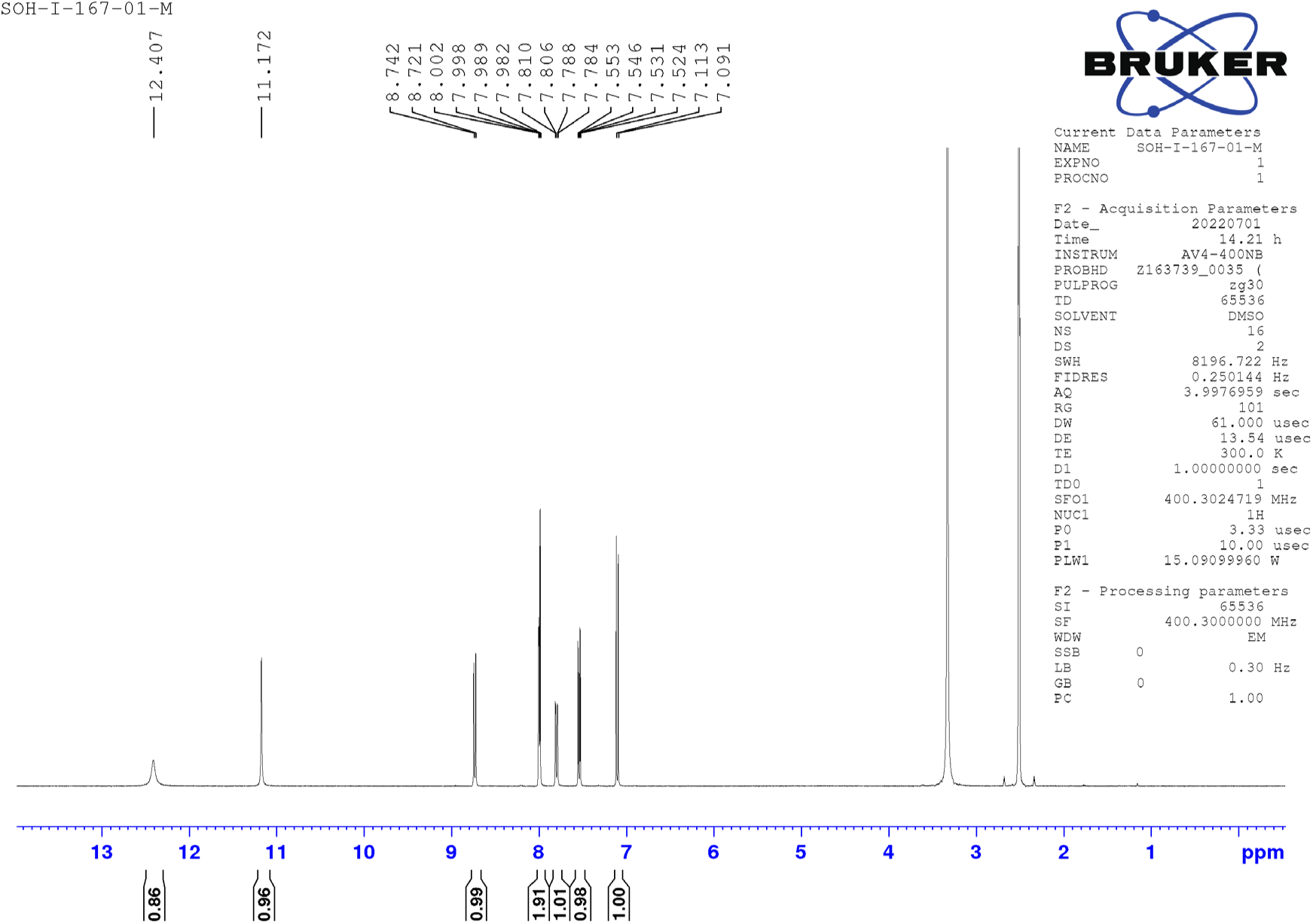

^1^HNMR (400 MHz, DMSO-d_6_) δ 12.4 (s, 1H), 11.17 (s, 1H), 8.73 (d, *J* = 8.4 Hz, 1H), 8.0 - 7.98 (m, 2H), 7.79 (dd, *J* = 8.7 Hz, 1.6 Hz, 1H), 7.53 (dd, *J* = 8.7 Hz, 2.8 Hz, 1H), 7.10 (d, *J* = 4.3 Hz, 1H); LC-MS (ESI); [M-H]: 348.0

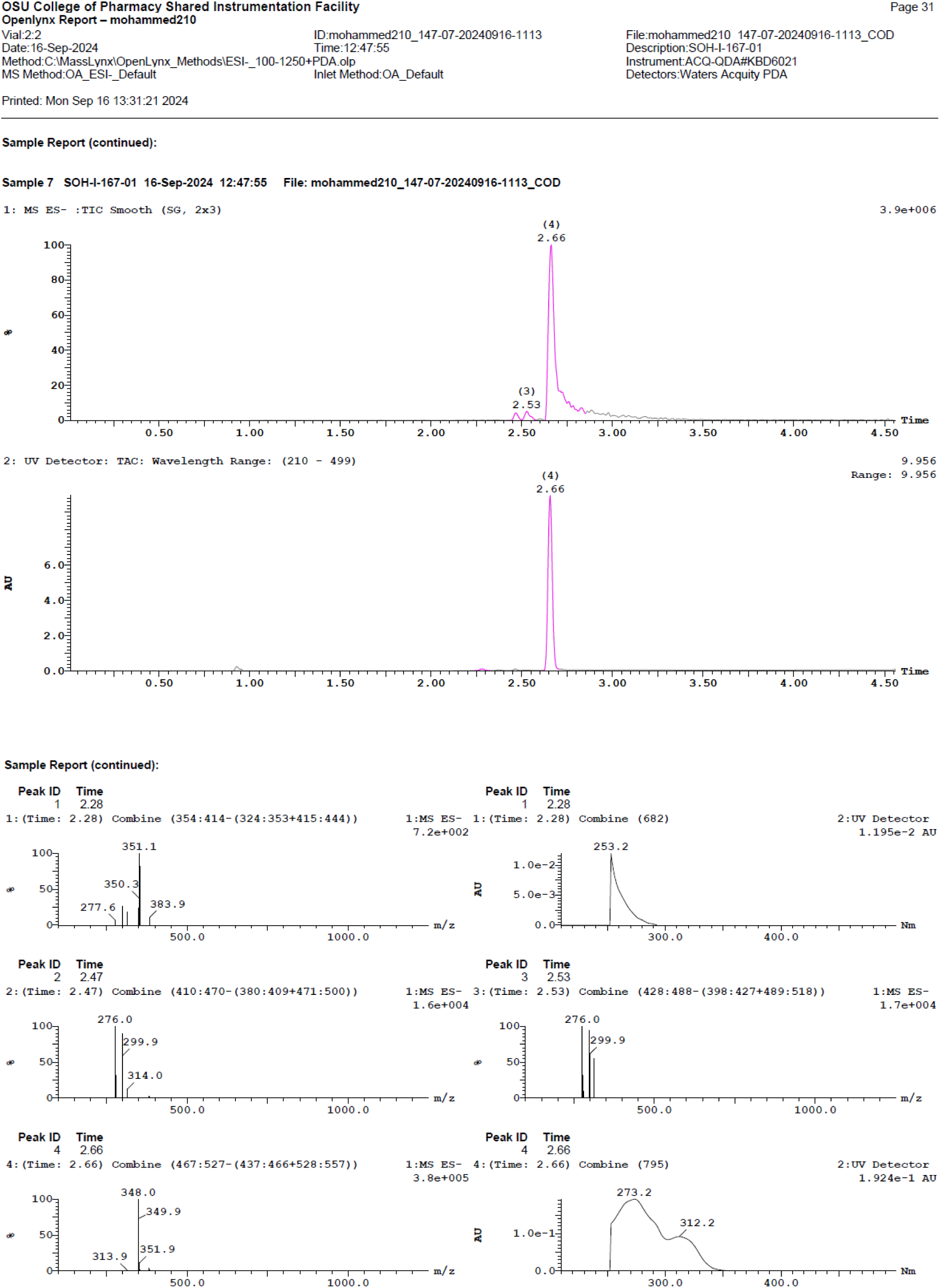

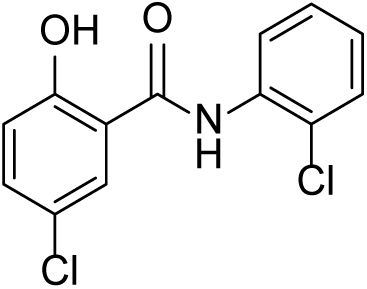

### 5-chloro-N-(2-chlorophenyl)-2-hydroxybenzamide (Nic-8)

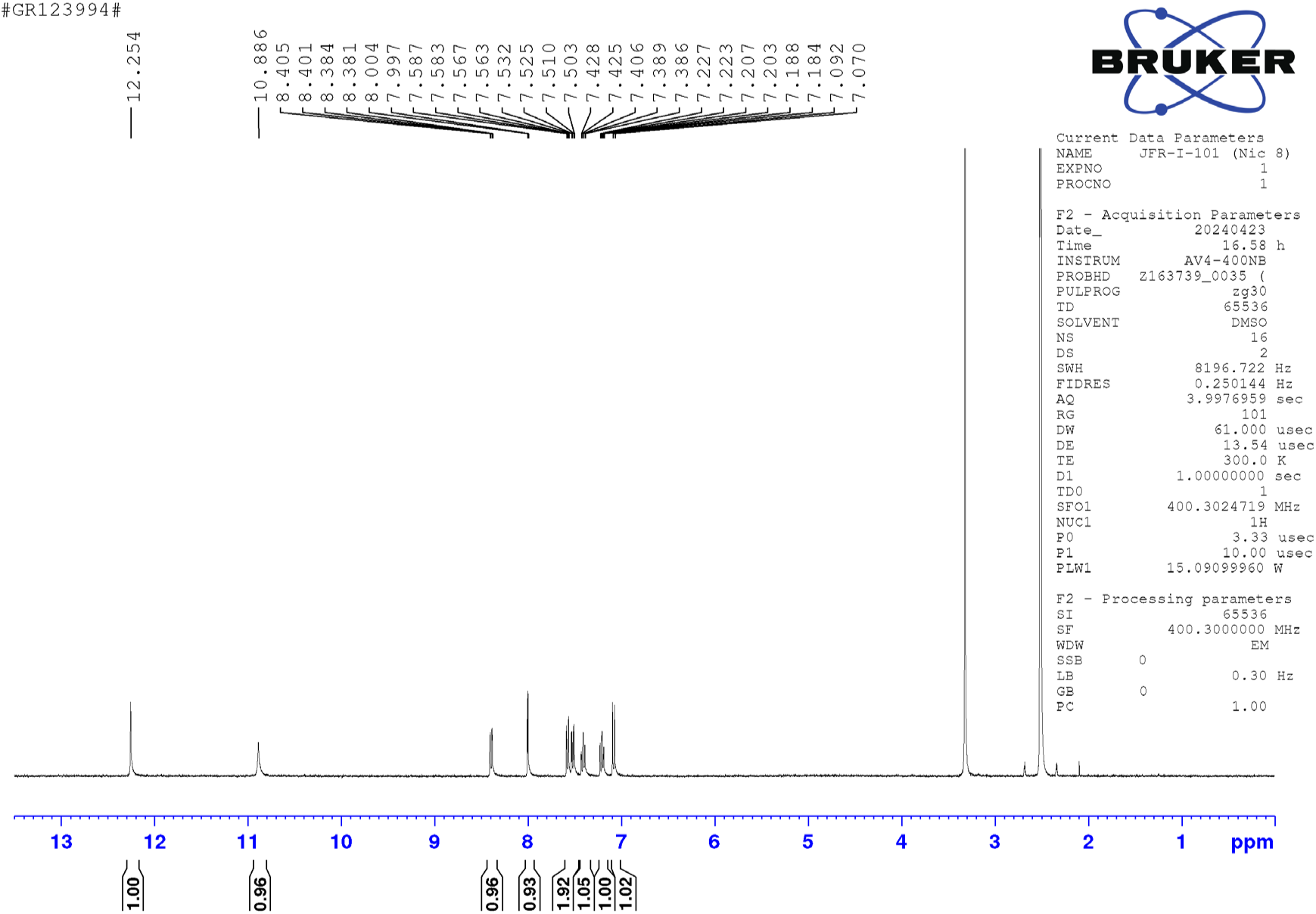

^1^HNMR (400 MHz, DMSO-d_6_) δ 12.25 (s, 1H), 10.88 (s, 1H), 8.39 (dd, *J* = 8.2 Hz, 1.48 Hz, 1H), 7.99 (d, *J* = 2.7 Hz, 1H), 7.58 - 7.50 (m, 2H), 7.40 (td, *J* = 8.7 Hz, 1.4 Hz, 1H), 7.20 (td, *J* = 7.9 Hz, 1.5 Hz, 1H), 7.06 (d, *J* = 8.7 Hz, 1H); LC-MS (ESI); [M+H]: 282.0; [M+2H]: 284.0; [M+4H]: 286.0

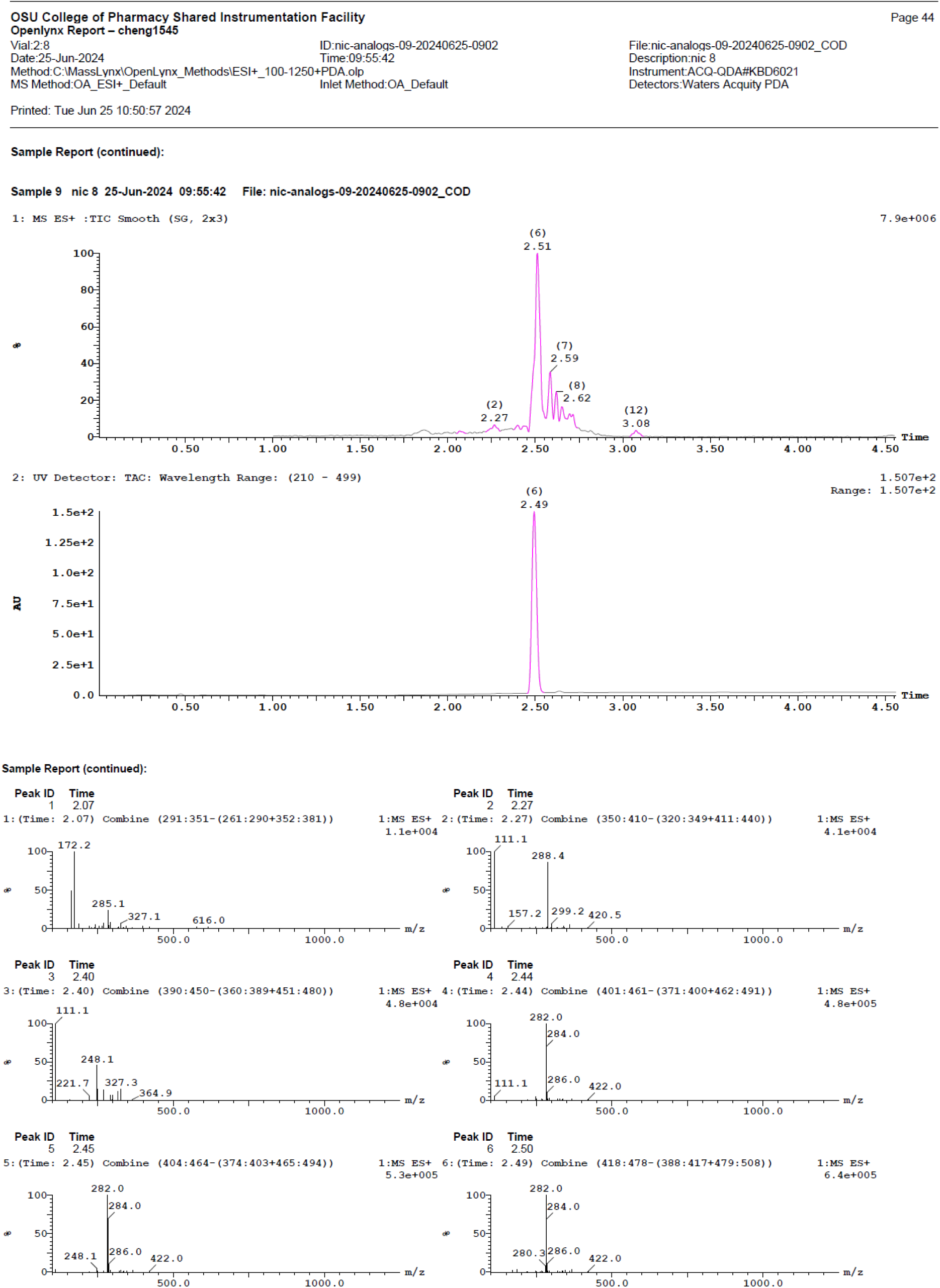

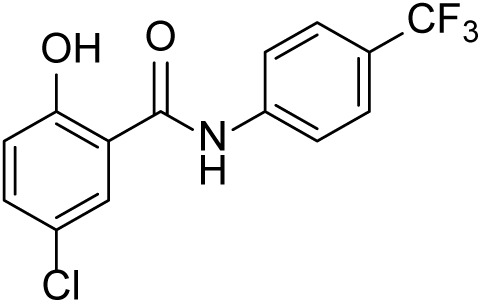

### 5-chloro-2-hydroxy-N-(4-(trifluoromethyl)phenyl)benzamide (Nic-11)

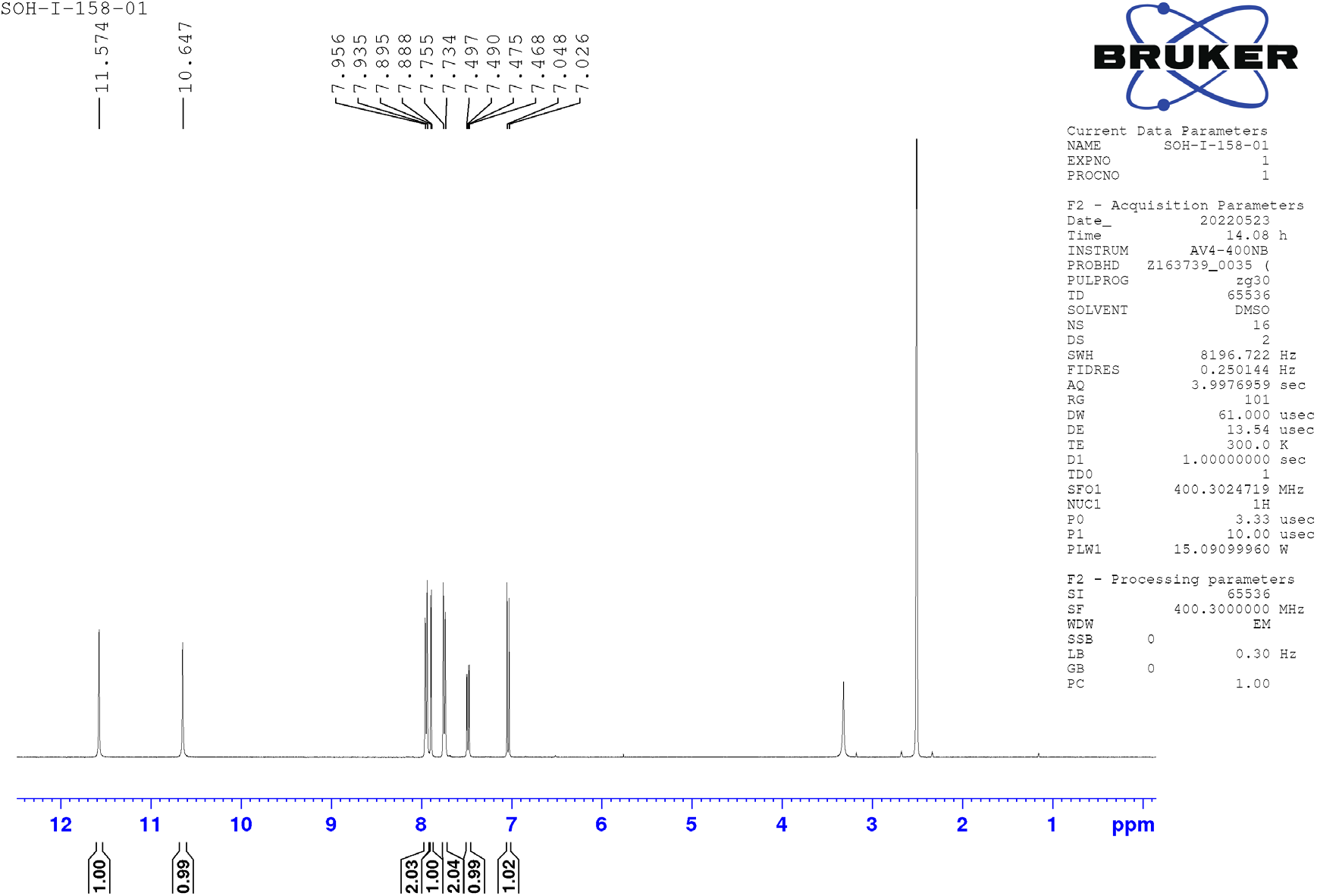

^1^HNMR (400 MHz, DMSO-d_6_) δ 11.5 (s, 1H), 10.6 (s, 1H), 7.94 (d, *J* = 8.4 Hz, 2H), 7.88 (d, *J* = 2.6 Hz, 1H), 7.74 (d, *J* = 8.6 Hz, 2H), 7.47 (dd, *J* = 8.8 Hz, 2.7 Hz, 1H), 7.03 (d, *J* = 8.7 Hz, 1H); LC-MS (ESI); [M-H]: 313.9

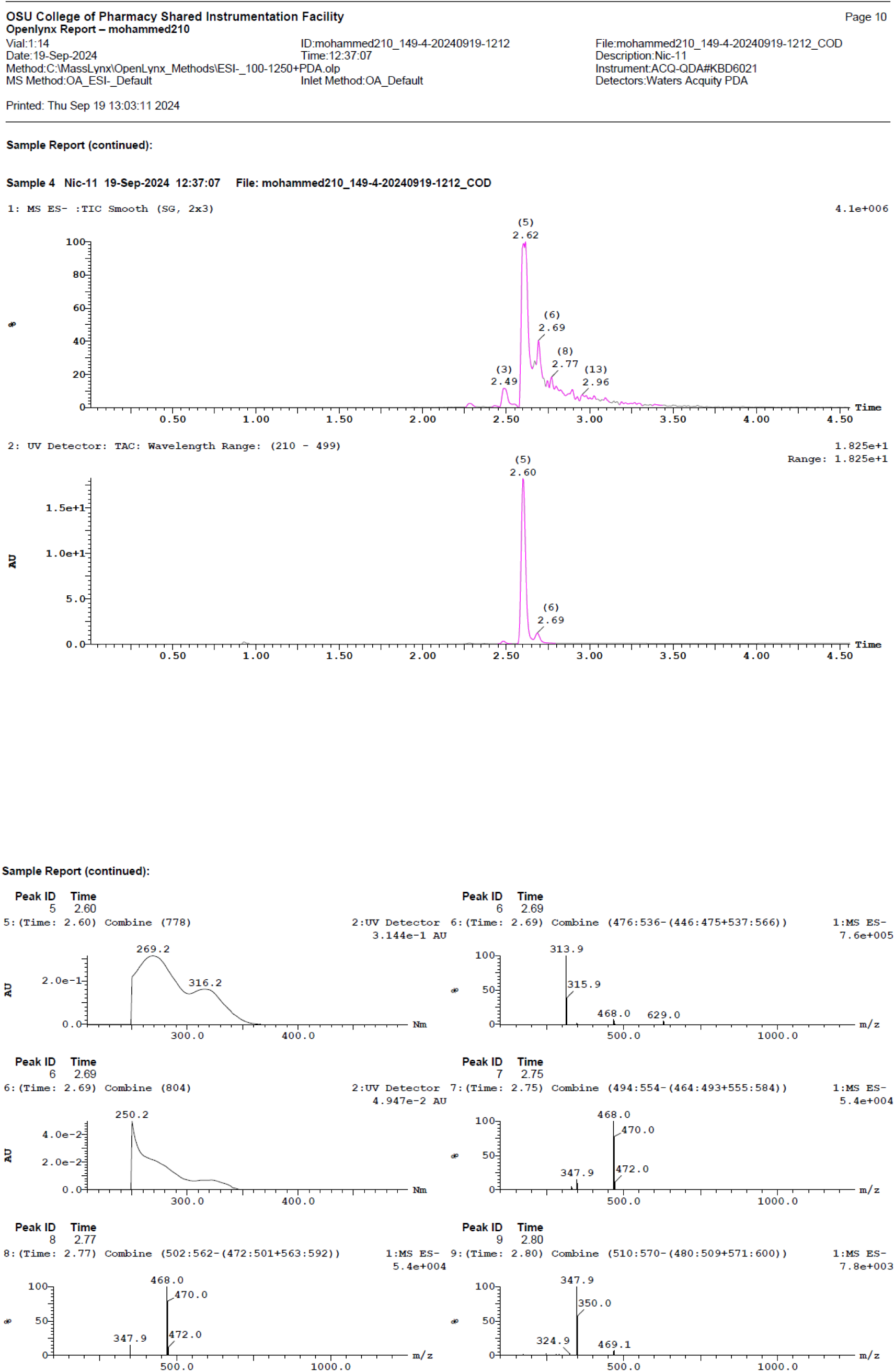

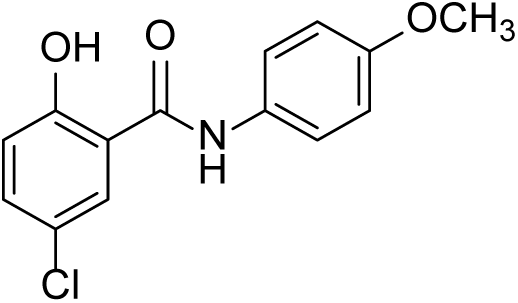

### 5-chloro-2-hydroxy-N-(4-methoxyphenyl)benzamide (Nic-12)

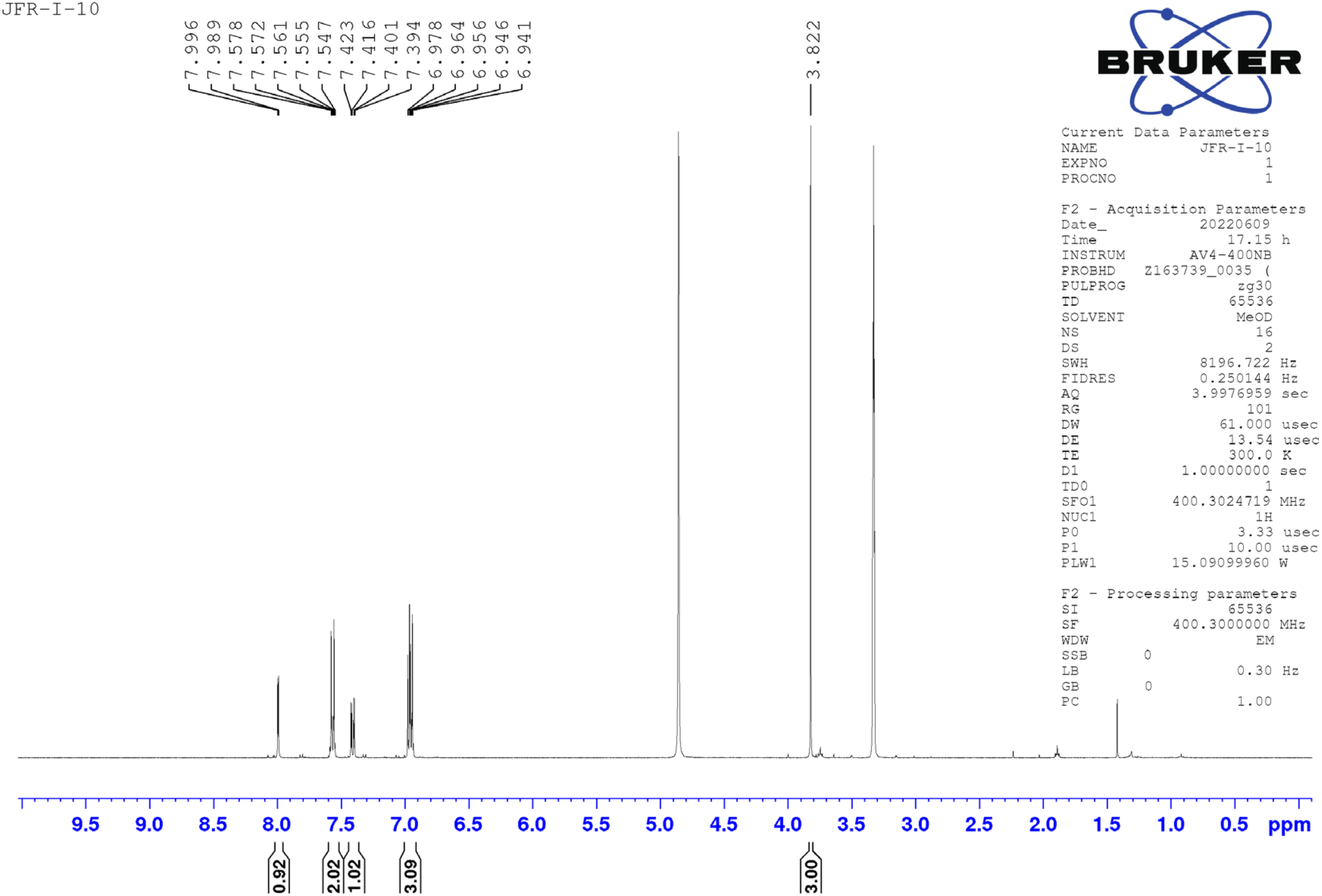

^1^HNMR (400 MHz, CD_3_OD) δ 7.98 (d, *J* = 2.6 Hz, 1H), 7.57 - 7.54 (m, 2H), 7.40 (dd, *J* = 8.8 Hz, 2.6 Hz, 1H), 6.97 - 6.94 (m, 3H), 3.82 (s, 3H); LC-MS (ESI); [M-H]: 276.0

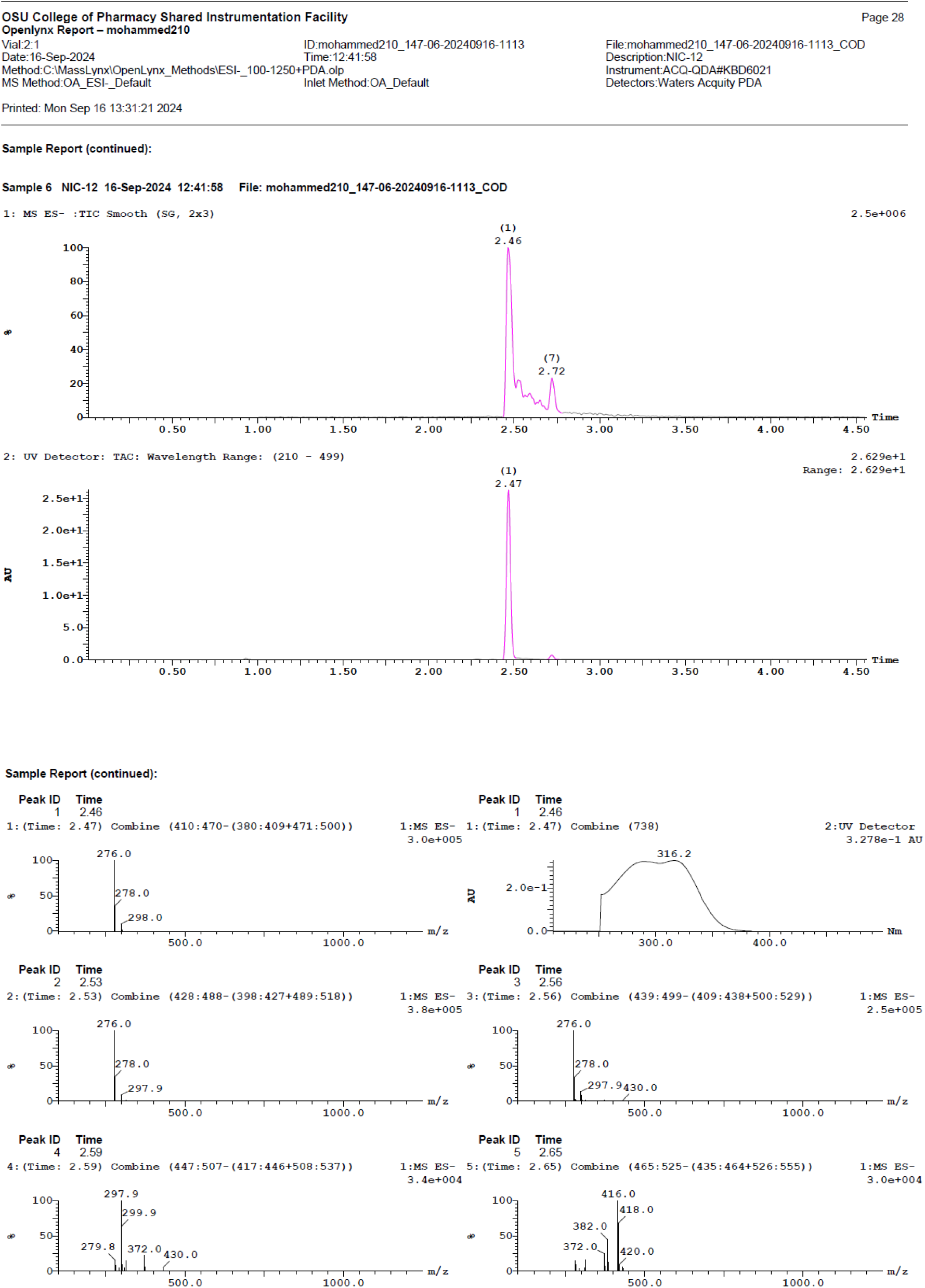

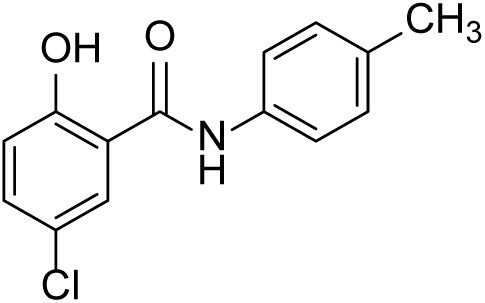

### 5-chloro-2-hydroxy-N-(p-tolyl)benzamide (Nic 13)1

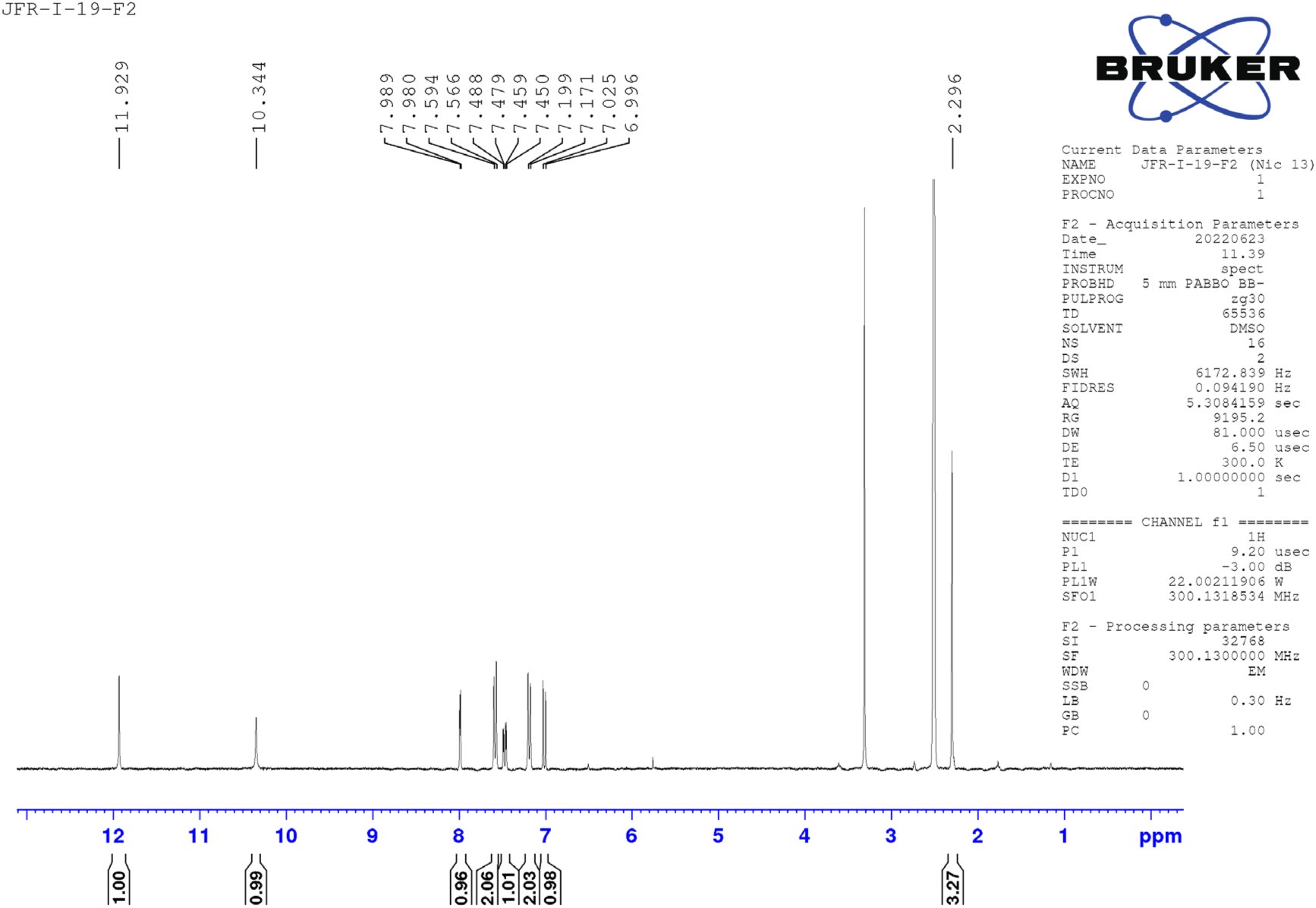

^1^HNMR (300 MHz, DMSO-d_6_) δ 11.92 (s, 1H), 11.34 (s, 1H), 7.97 (d, *J* = 2.6 Hz, 1H), 7.57 (d, *J* = 8.3 Hz, 2H), 7.46 (dd, *J* = 8.7 Hz, 2.6 Hz, 1H), 8.18 (d, *J* = 8.3 Hz, 2H), 7.0 (d, *J* = 7.2 Hz, 1H), 2.29 (s, 3H); LC-MS (ESI); [M-H]: 262.1

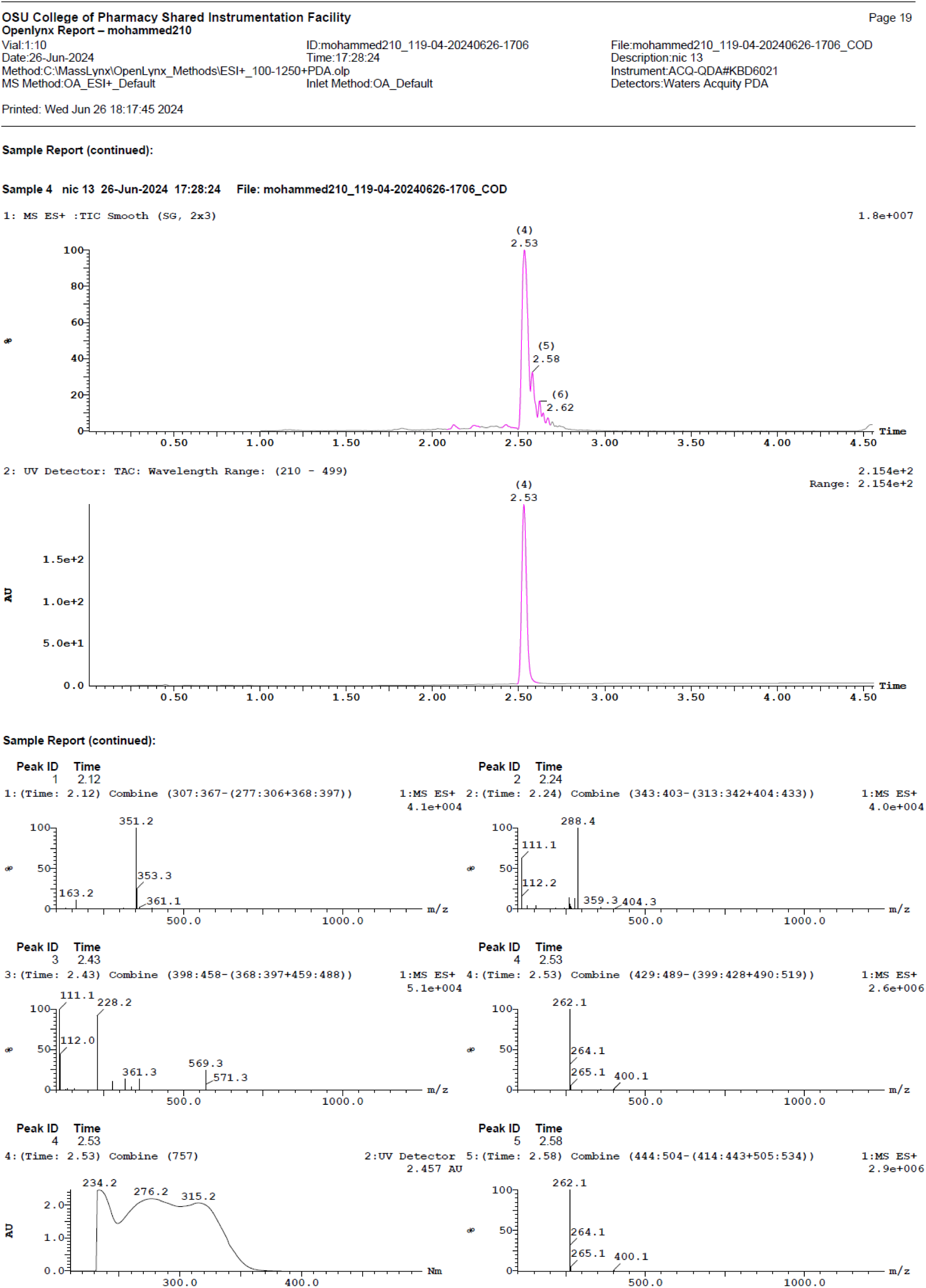

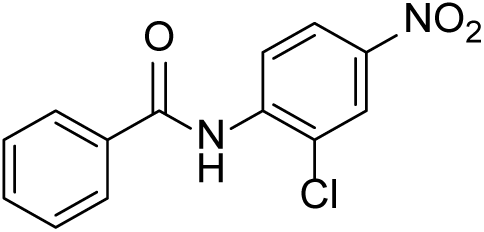

### N-(2-chloro-4-nitrophenyl)benzamide (Nic-14)^2^

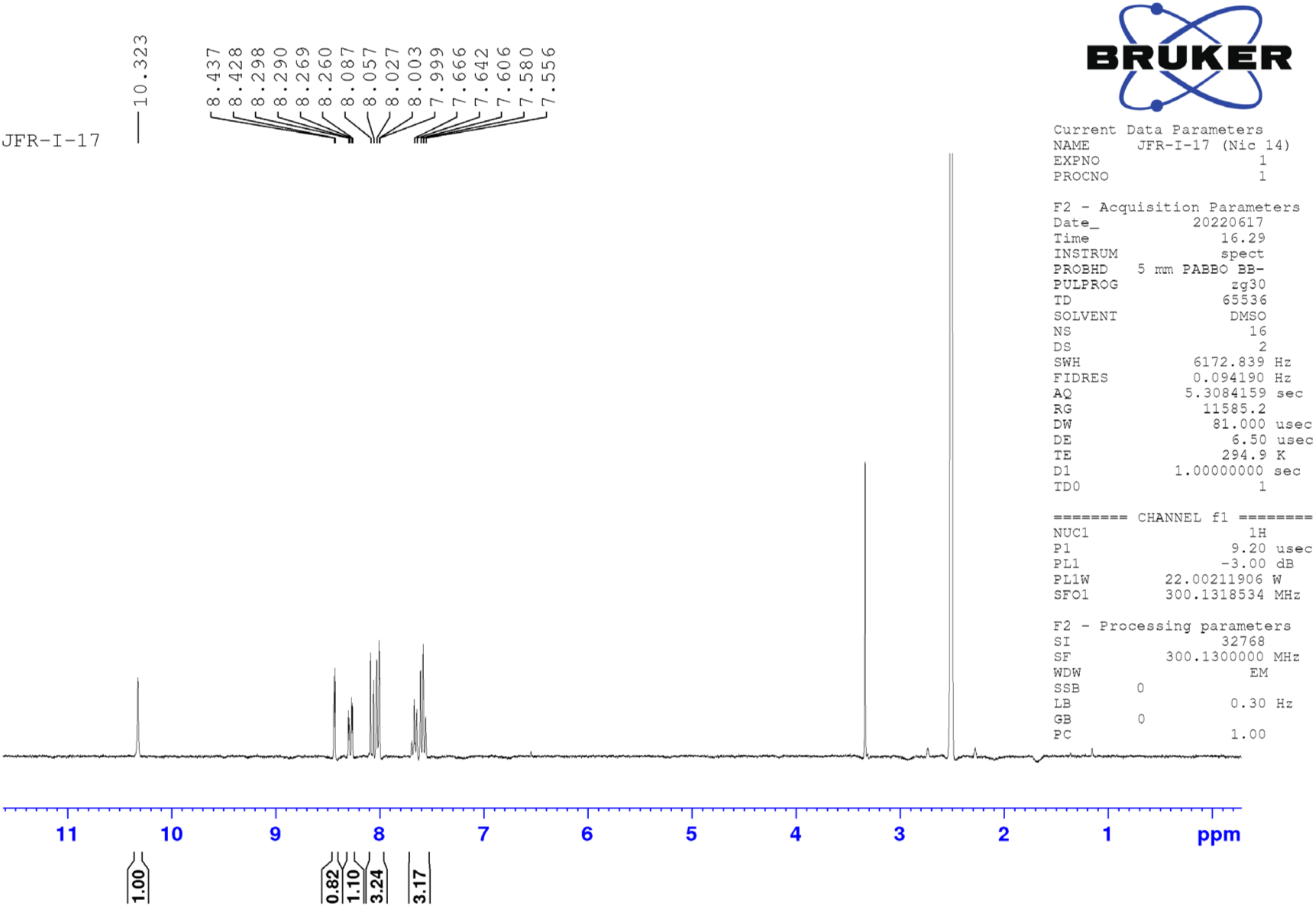

^1^HNMR (300 MHz, DMSO-d_6_) δ 10.32 (s, 1H), 8.42 (d, *J* = 2.5 Hz, 1H), 8.27 (dd, *J* = 8.9 Hz, 2.5 Hz, 1H), 8.08 -7.99 (m, 3H), 7.66 -7.55 (m, 3H); LC-MS (ESI); [M+H]: 277.1

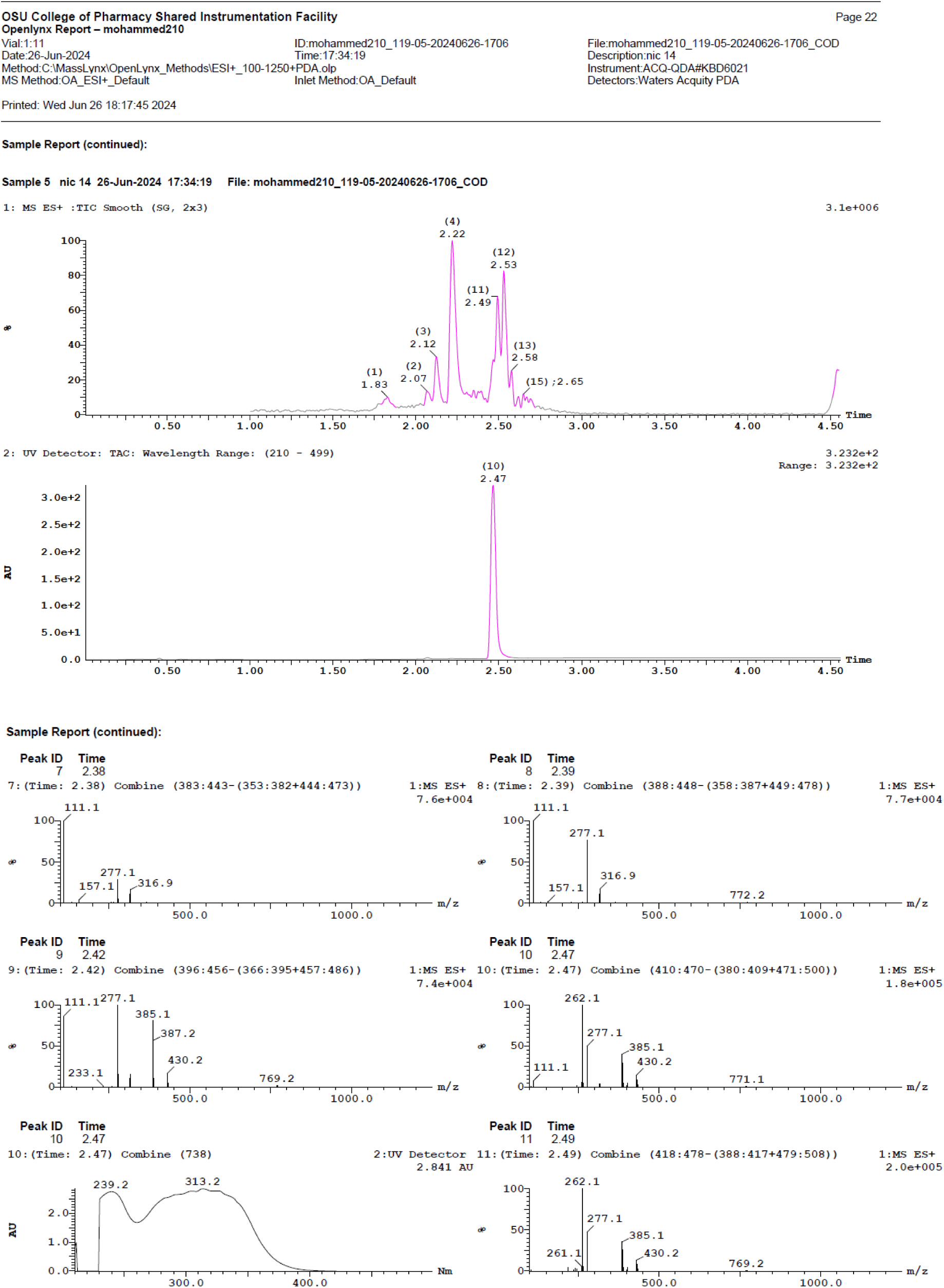

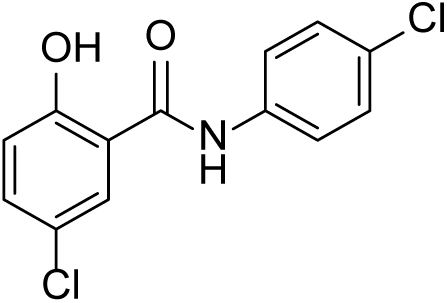

### 5-chloro-N-(4-chlorophenyl)-2-hydroxybenzamide (Nic-15)

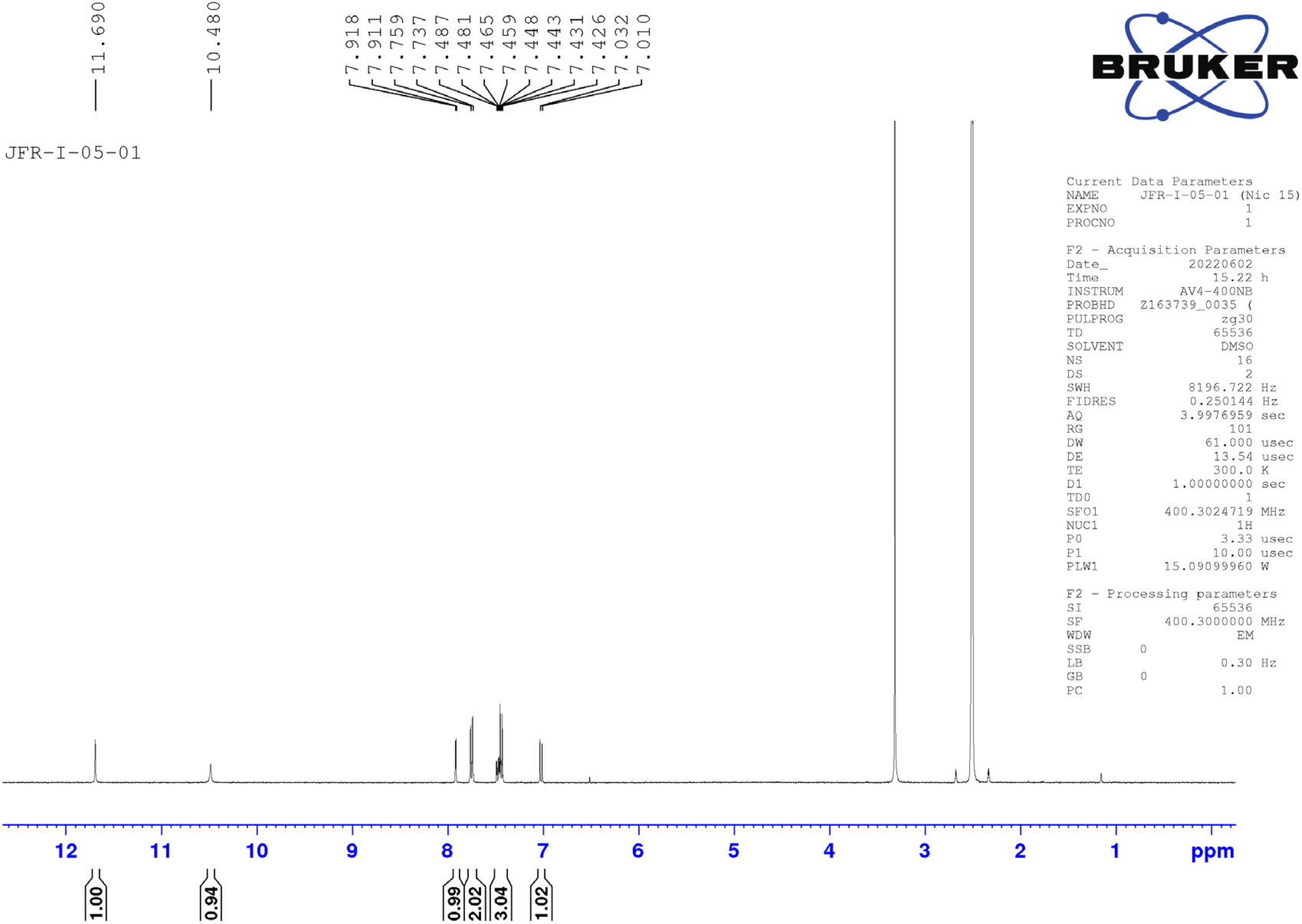

^1^HNMR (400 MHz, DMSO-d_6_) δ 11.69 (s, 1H), 10.48 (s, 1H), 7.91 (d, *J* = 2.6 Hz, 1H), 7.74 (d, *J* = 8.8 Hz, 2H), 7.48 - 7.42 (m, 3H), 7.01 (d, *J* = 8.8 Hz, 1H), ; LC-MS (ESI); [M+]: 282.1, [M+2H]: 284.0, [M+4H]: 286.0

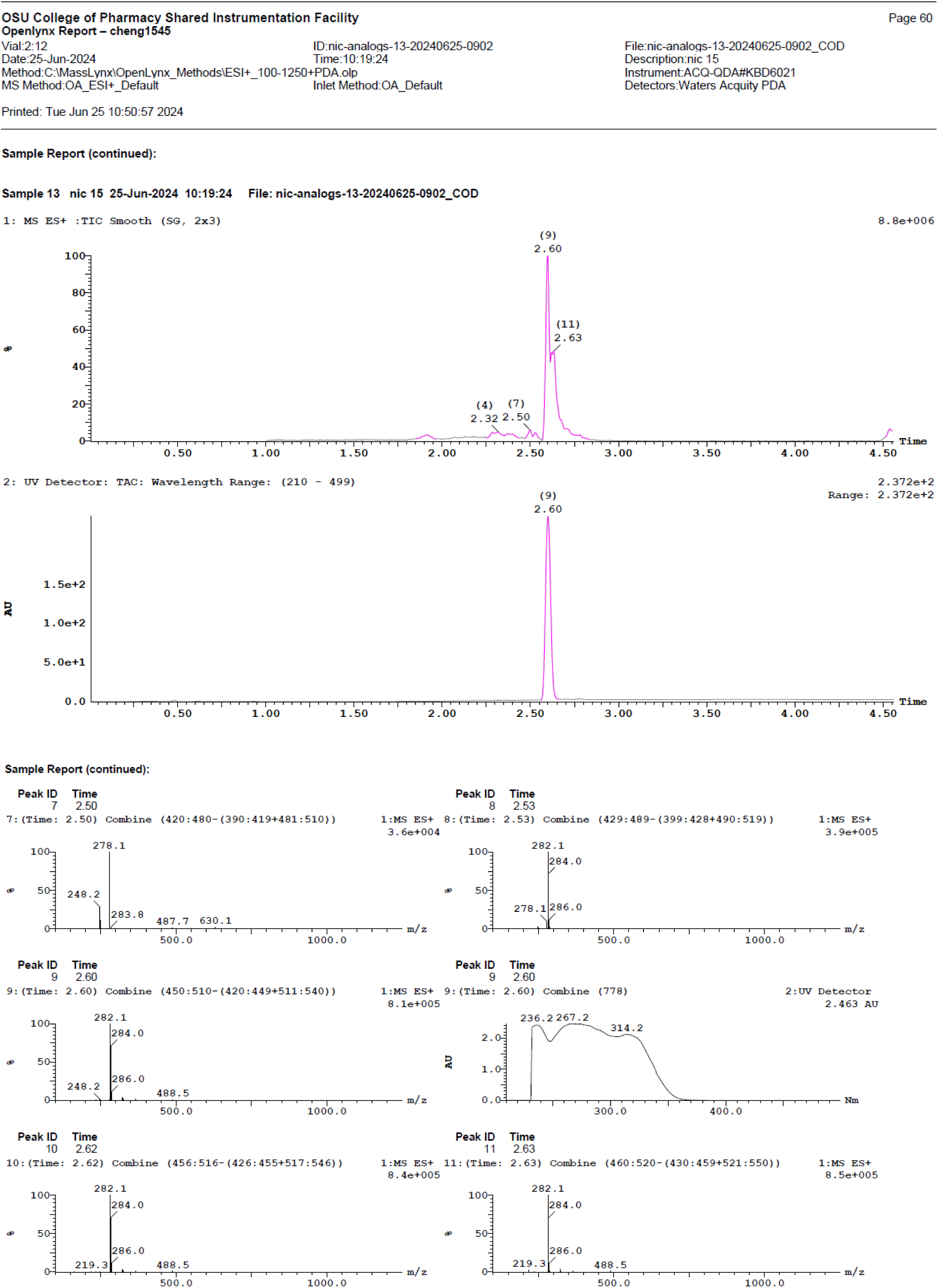

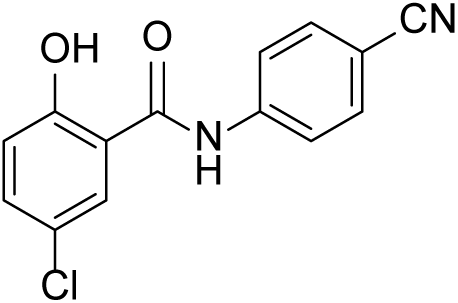

### 5-chloro-N-(4-cyanophenyl)-2-hydroxybenzamide (Nic-20)

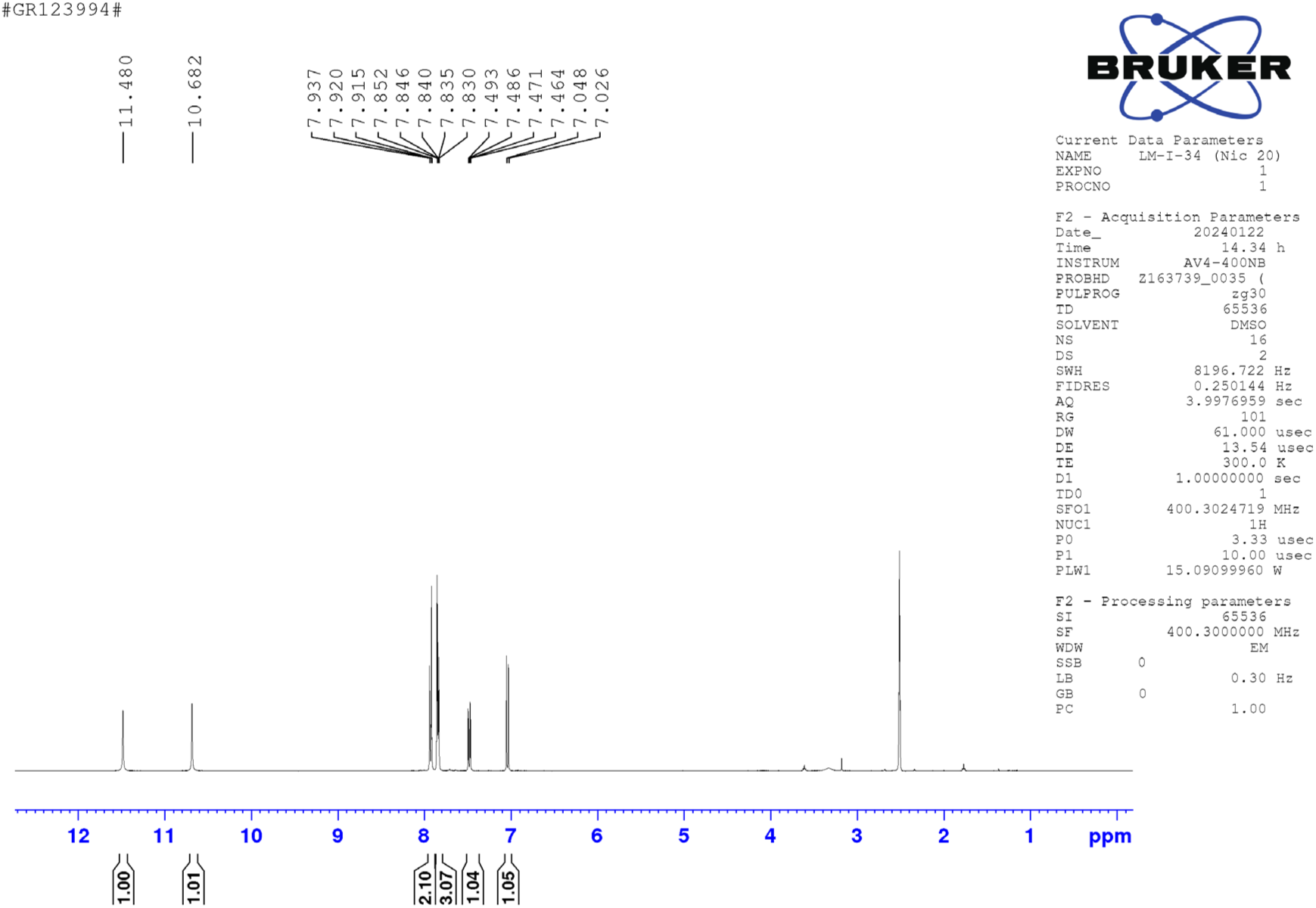

^1^HNMR (400 MHz, DMSO-d_6_) δ 11.48 (s, 1H), 10.68 (s, 1H), 7.93 - 7.91 (m, 2H), 7.85 - 7.82 (m, 3H), 7.47 (dd, *J* = 8.8 Hz, 2.7 Hz, 1H), 7.03 (d, *J* = 8.8 Hz, 1H); LC-MS (ESI); [M-H]: 271.0

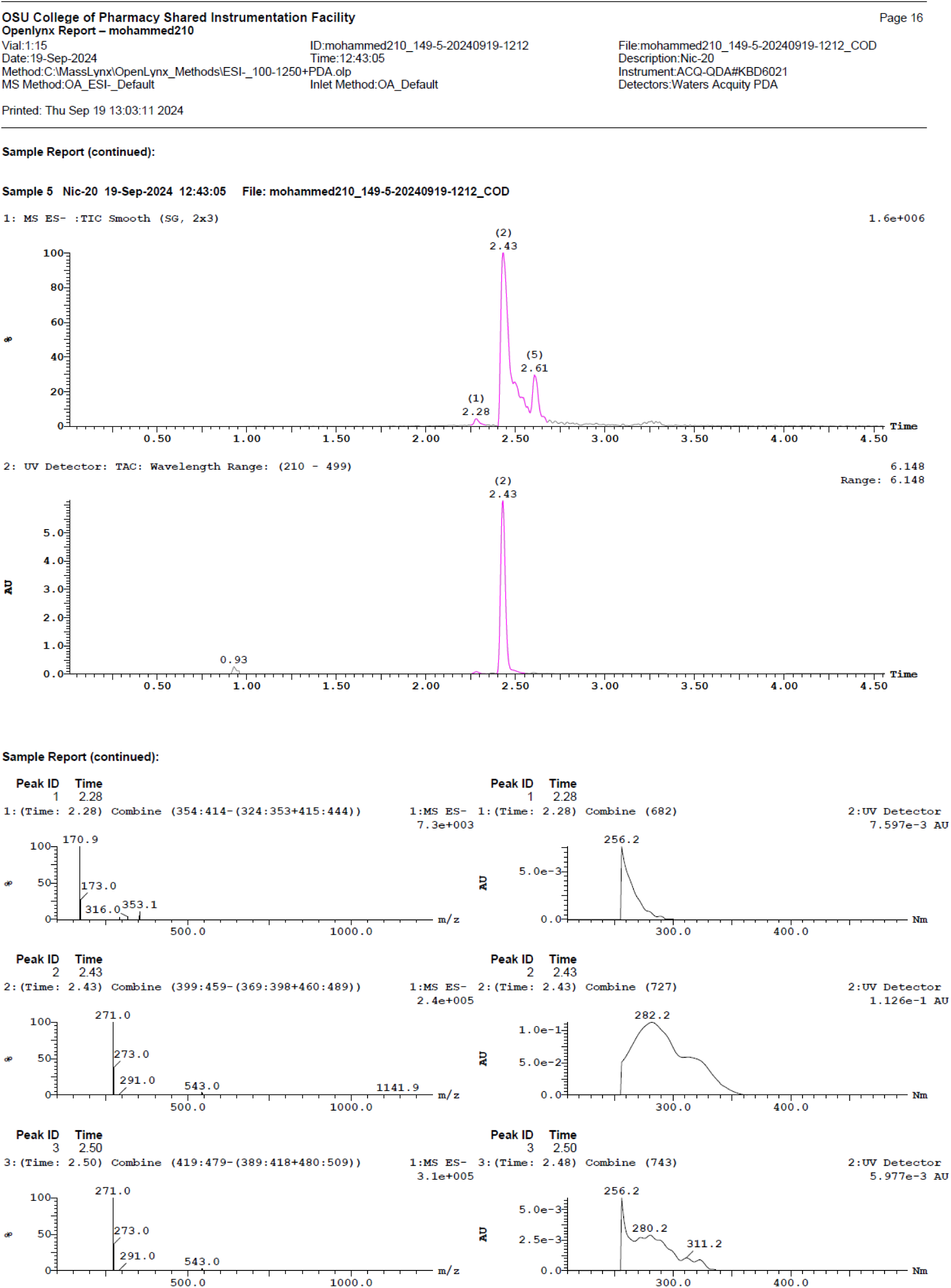

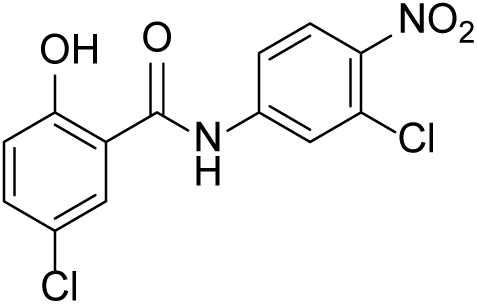

### 5-chloro-N-(3-chloro-4-nitrophenyl)-2-hydroxybenzamide (Nic-22)

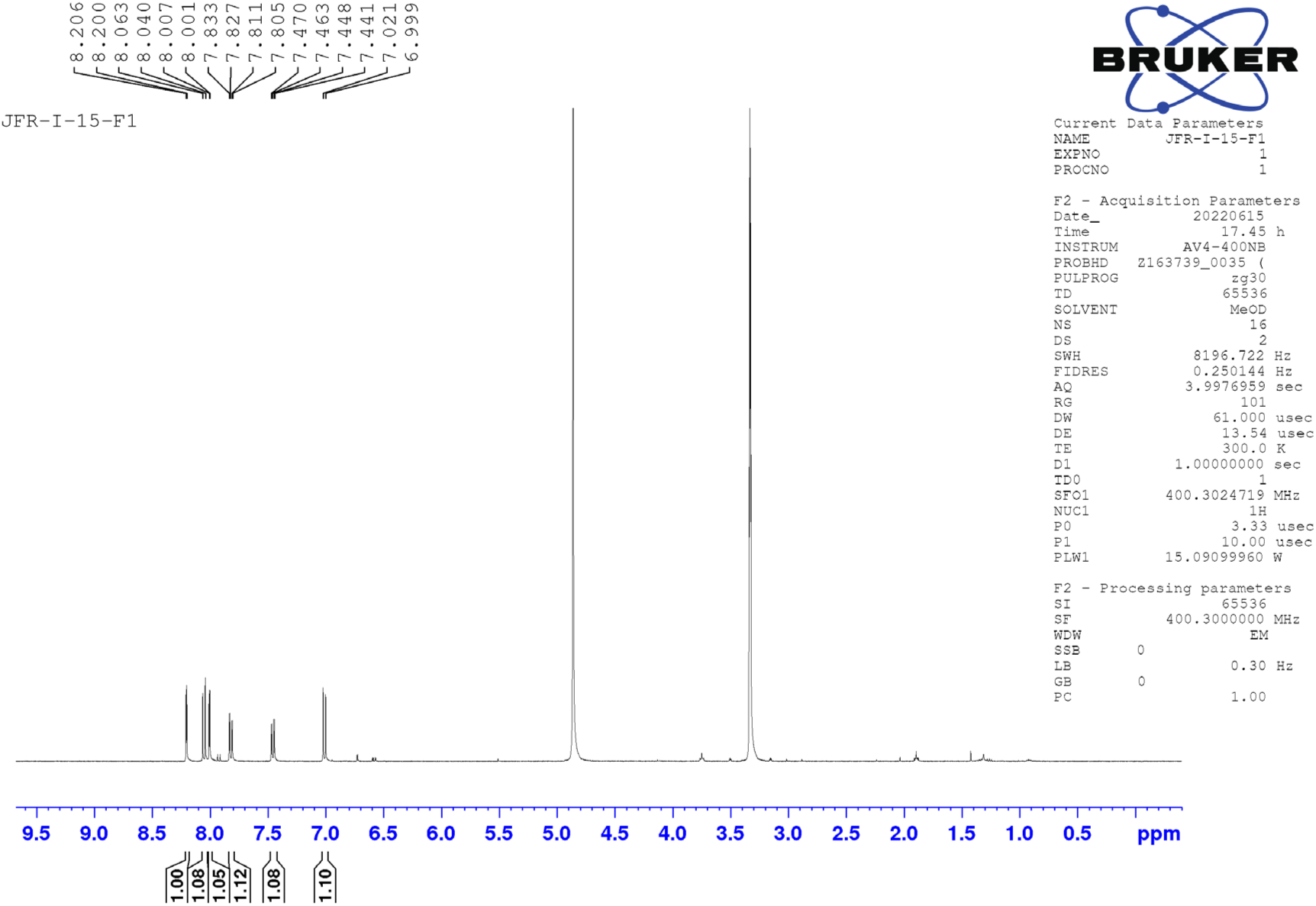

^1^HNMR (400 MHz, CD_3_OD) δ 8.20 (d, *J* = 2.2 Hz, 1H), 8.05 (d, *J* = 8.9Hz, 1H), 8.0 (d, *J* = 2.6 Hz, 1H), 7.81(dd, *J* = 8.9 Hz, 2.2 Hz, 1H), 7.81(dd, *J* = 8.9 Hz, 2.2 Hz, 1H), 6.9 (d, *J* = 8.8 Hz, 1H); LC-MS (ESI); [M-H]: 324.9

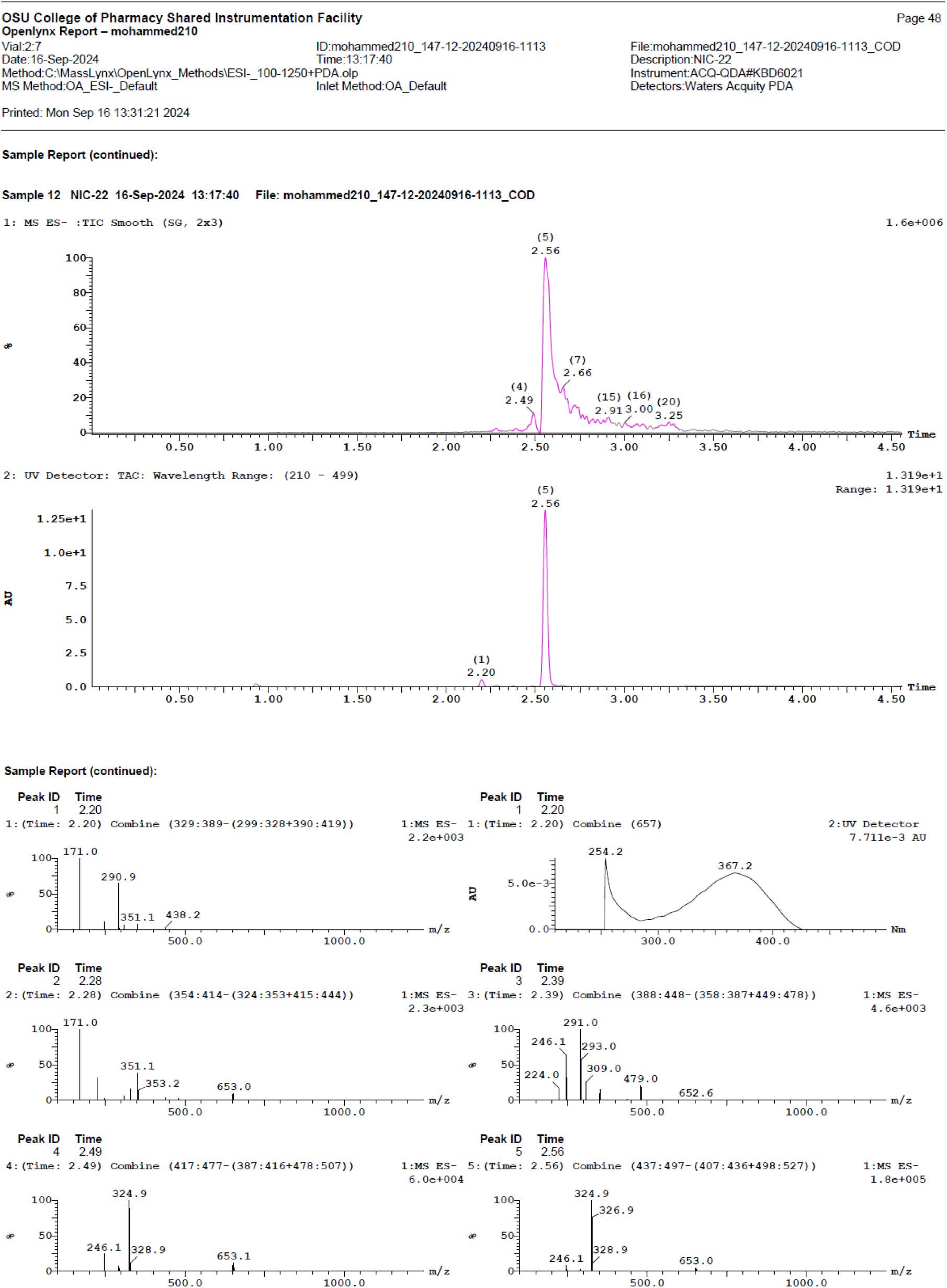

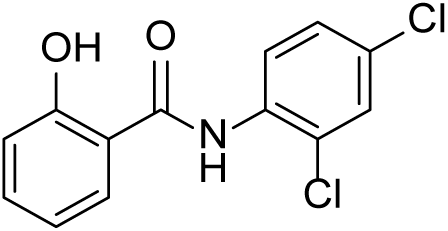

### N-(2,4-dichlorophenyl)-2-hydroxybenzamide (Nic-40)

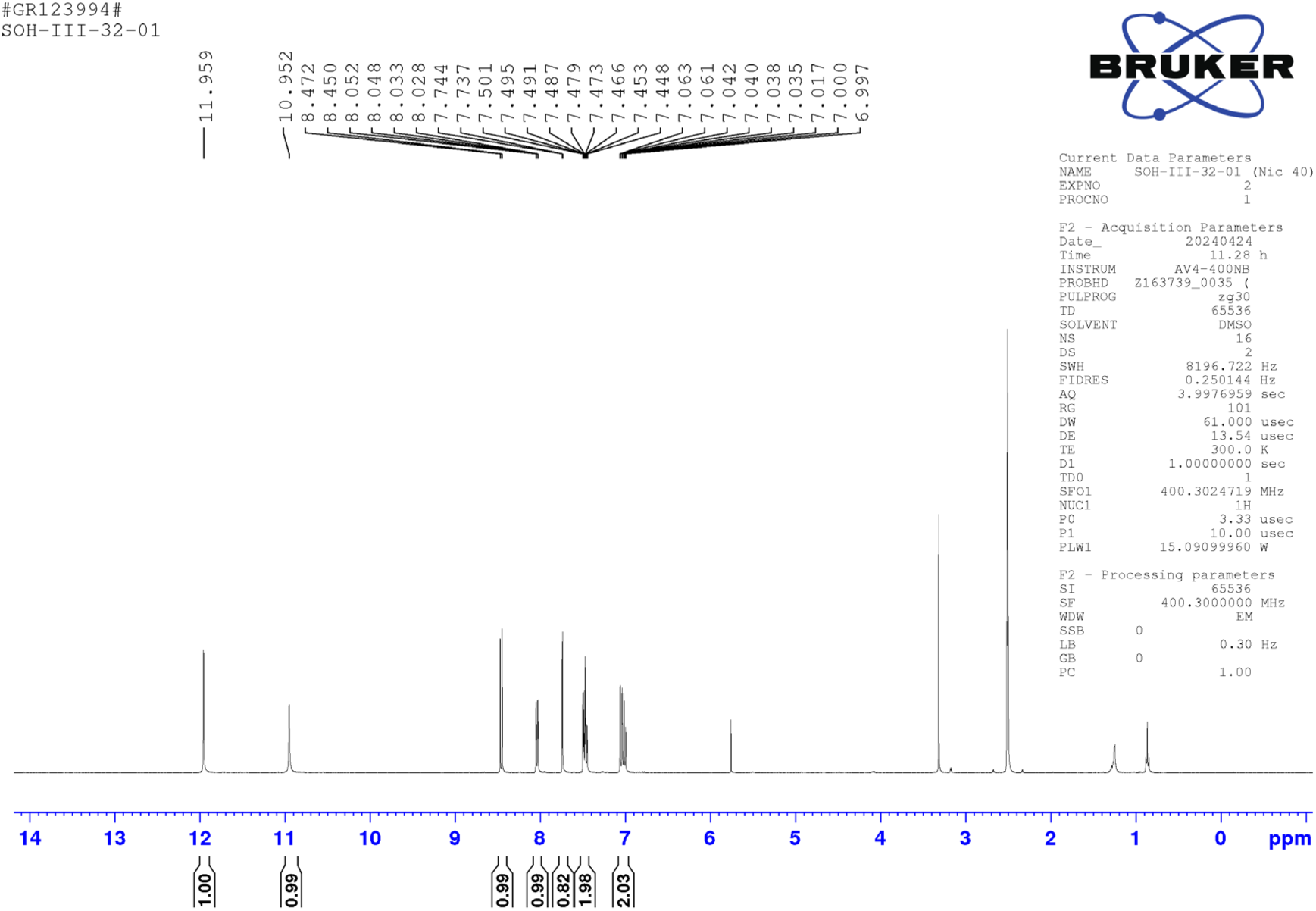

^1^HNMR (400 MHz, DMSO-d_6_) δ 11.95 (s, 1H), 10.95 (s, 1H), 8.45 (d, *J* = 8.8 Hz, 1H), 8.03 (dd, *J* = 7.9Hz, 1.7 Hz, 1H), 7.73 (d, *J* = 2.4 Hz, 1H), 7.50 - 7.44 (m, 2H), 7.06 - 6.99 (m, 2H); LC-MS (ESI); [M+]: 282.0, [M+2H]: 284.0, [M+4H]: 286.0

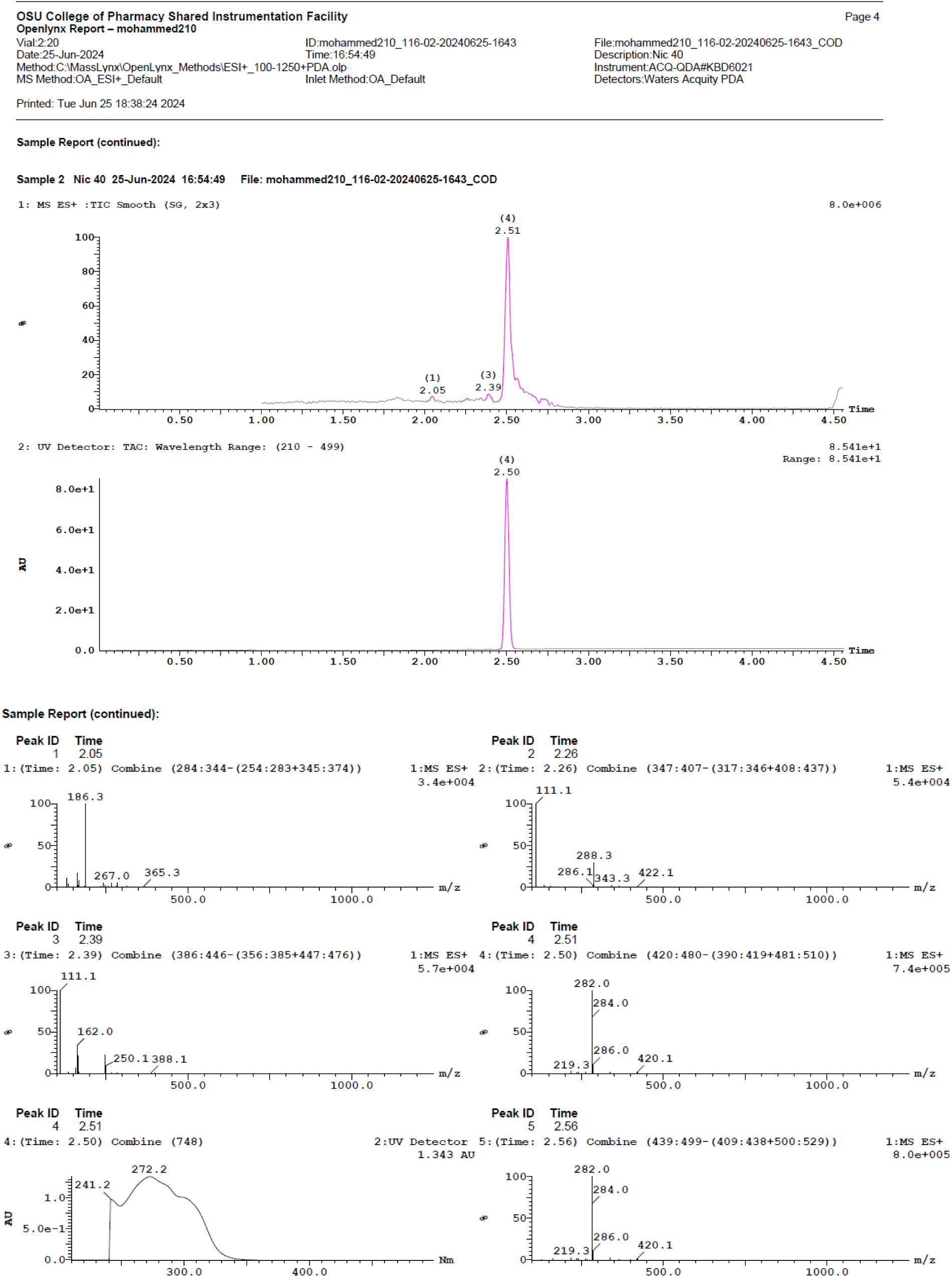

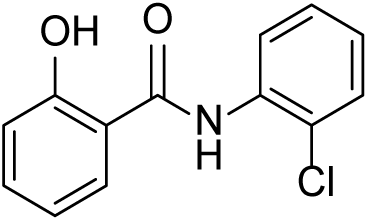

### N-(2-chlorophenyl)-2-hydroxybenzamide (Nic-41)

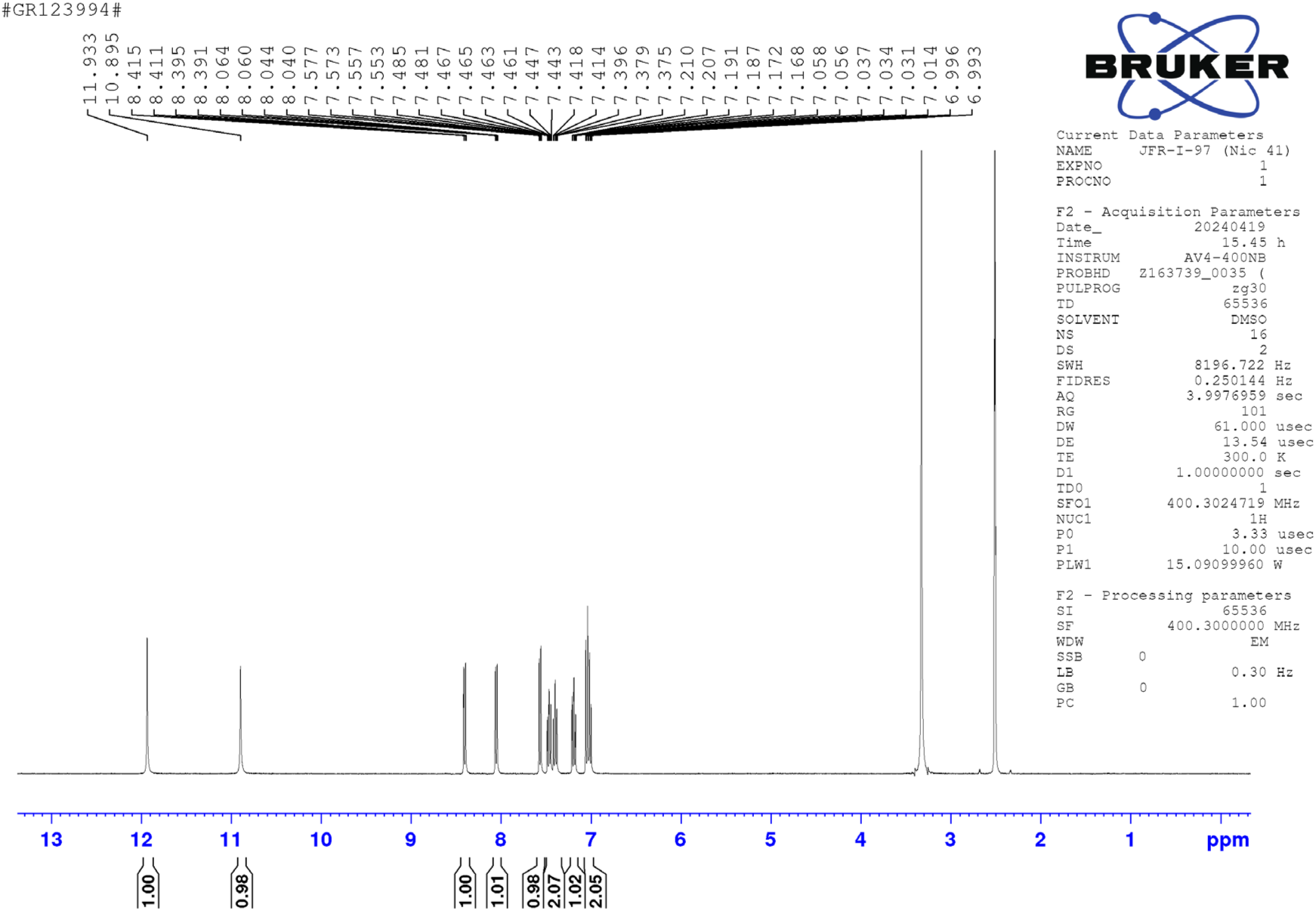

^1^HNMR (400 MHz, DMSO-d_6_) δ 11.93 (s, 1H), 10.89 (s, 1H), 8.40 (dd, *J* = 8.2 Hz, 1.5 Hz, 1H), 8.04 (dd, *J* = 7.9 Hz, 1.7 Hz, 1H), 7.56 (dd, *J* = 8.0 Hz, 1.4 Hz, 1H), 7.48 - 7.37 (m, 2H), 7.18 (td, *J* = 7.9 Hz, 1.6 Hz, 1H), 7.05 - 6.99 (m, 2H); LC-MS (ESI); [M+H]: 248.1

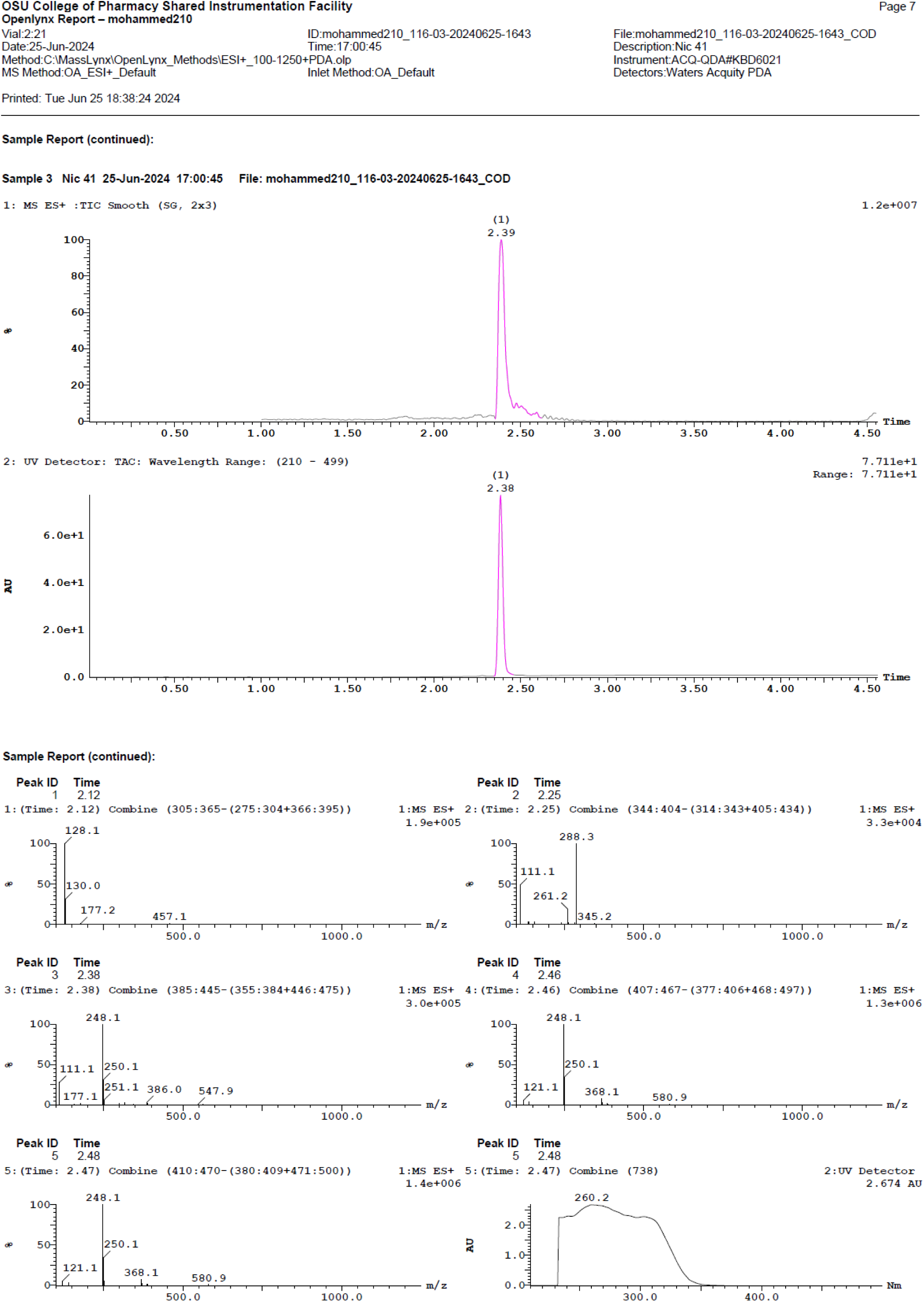

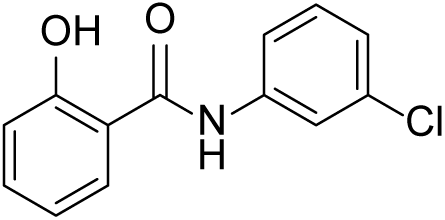

### N-(3-chlorophenyl)-2-hydroxybenzamide (Nic-42)

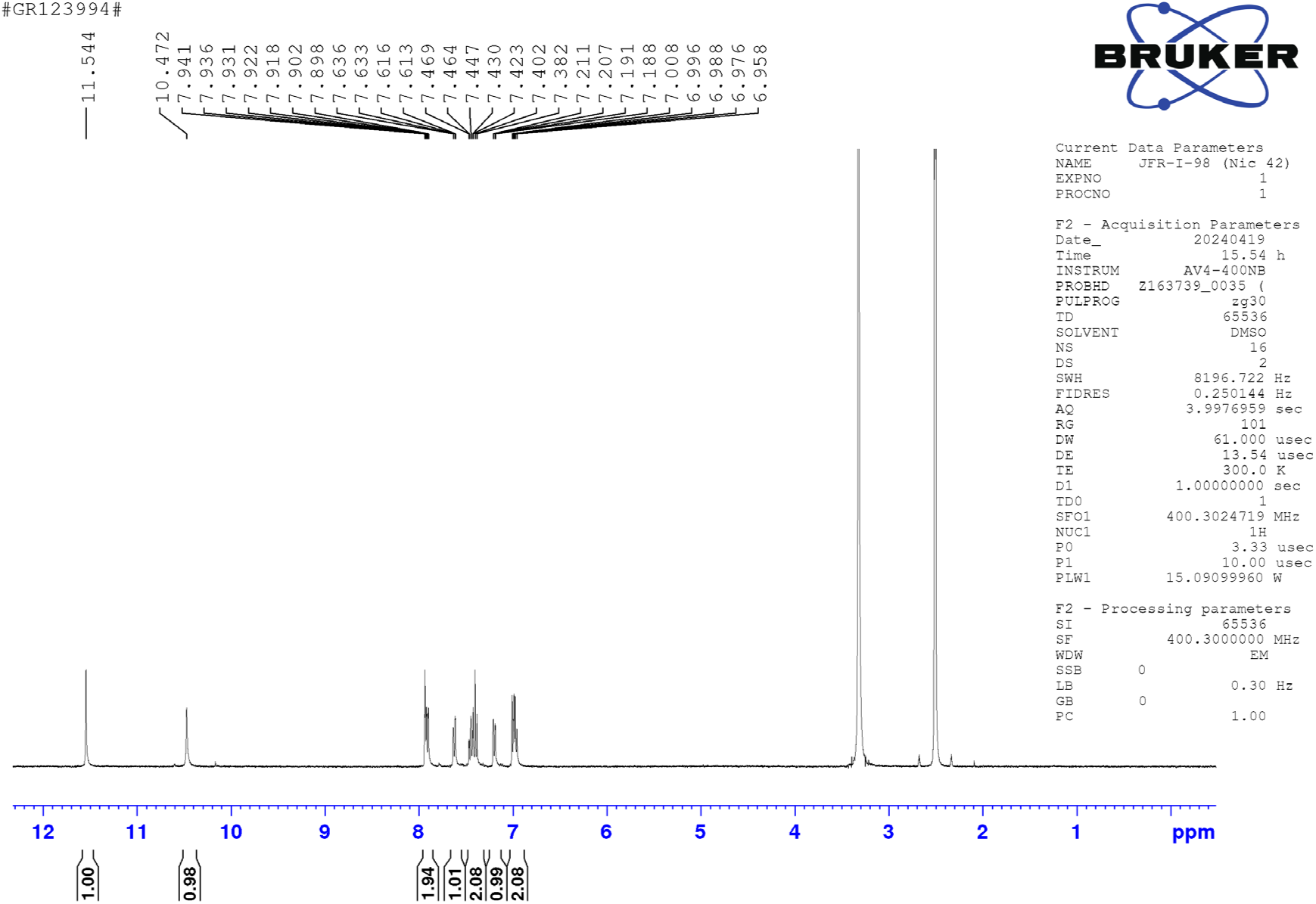

^1^HNMR (400 MHz, DMSO-d_6_) δ 11.54 (s, 1H), 10.47 (s, 1H), 7.94 - 7.89 (m, 2H), 7.62 (dd, *J* = 8.2 Hz, 1.0 Hz, 1H), 7.46 - 7.38 (m, 2H), 7.19 (dd, *J* = 7.9 Hz, 1.2 Hz, 1H), 7.00 - 6.95 (m, 2H); LC-MS (ESI); [M+H]: 248.1

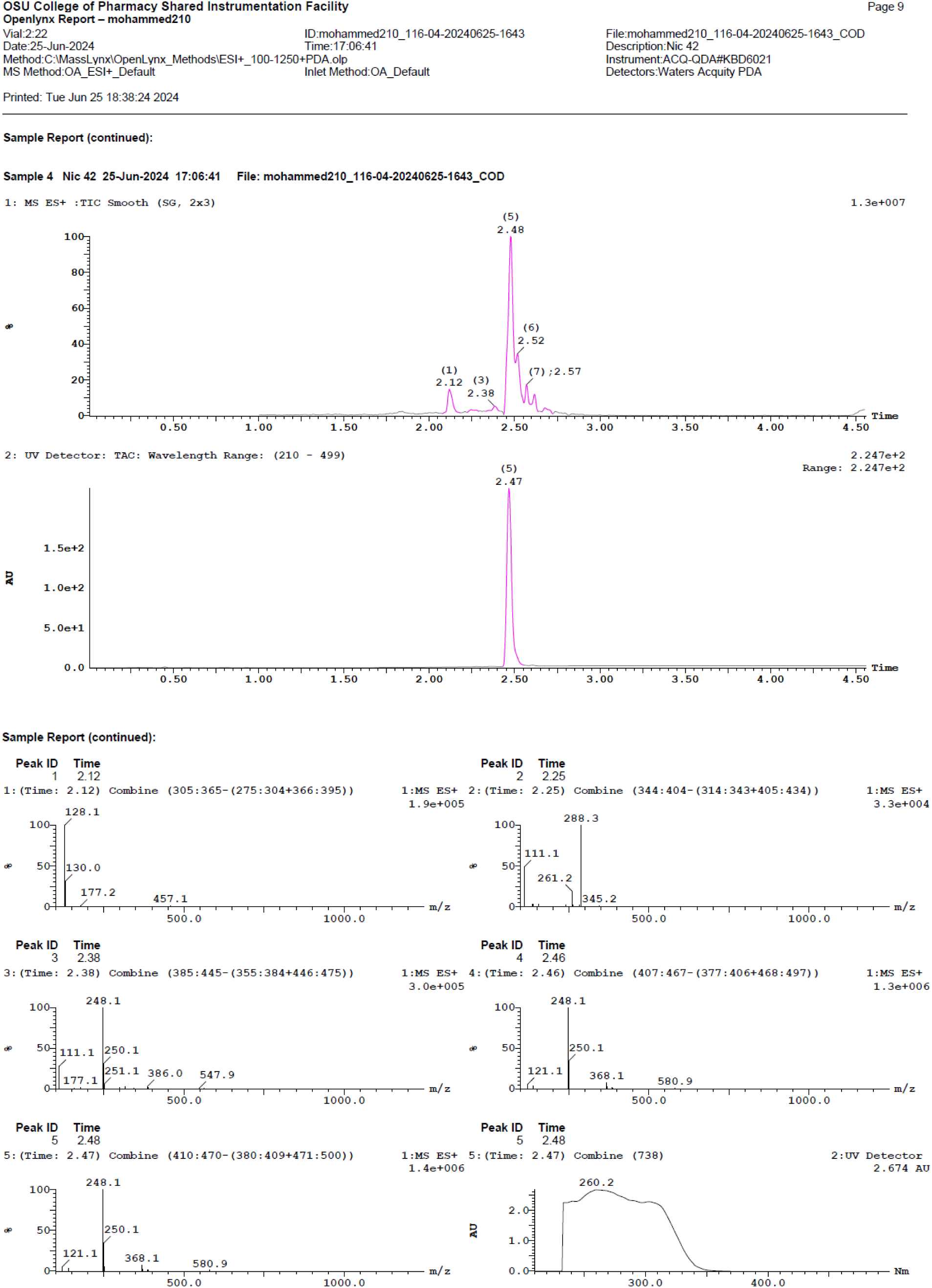

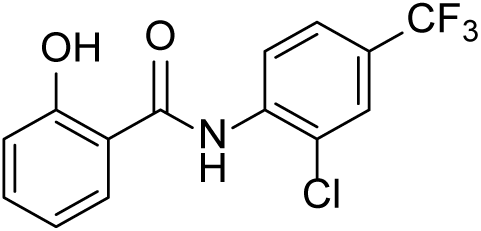

### N-(2-chloro-4-(trifluoromethyl)phenyl)-2-hydroxybenzamide (Nic-43)^7^

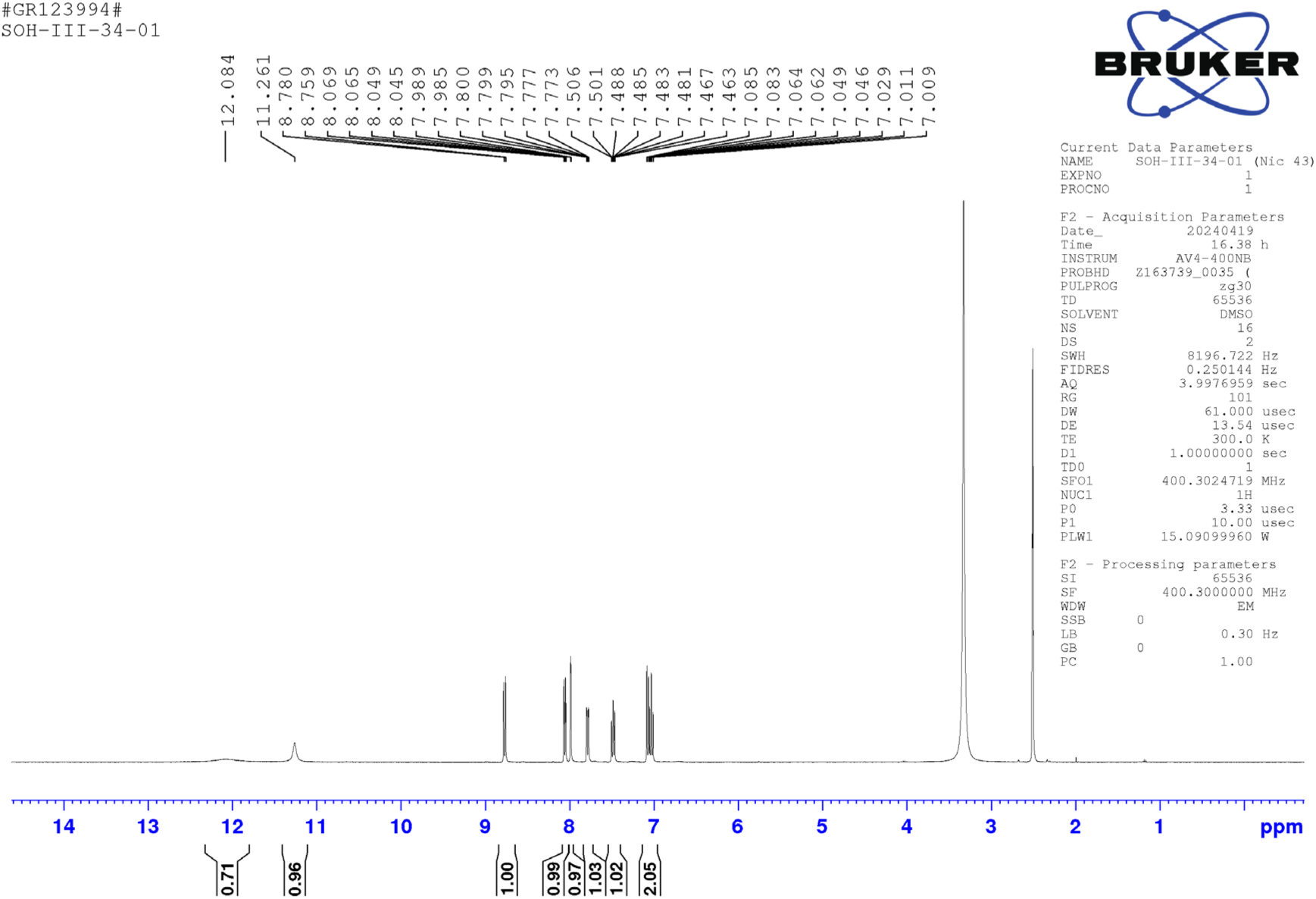

^1^HNMR (400 MHz, DMSO-d_6_) δ 12.08 (s, 1H), 11.26 (s, 1H), 8.76 (d, *J* = 8.2 Hz, 1.0 Hz, 1H), 8.05 (d, *J* = 7.9 Hz, 1.7 Hz, 1H), 7.98 (d, *J* = 1.6 Hz, 1H), 7.78 (dd, *J* = 9.3 Hz, 1.5 Hz, 1H), 7.50 - 7.46 (m, 1H), 7.08 - 7.00 (m, 2H); LC-MS (ESI); [M+H]: 316.1

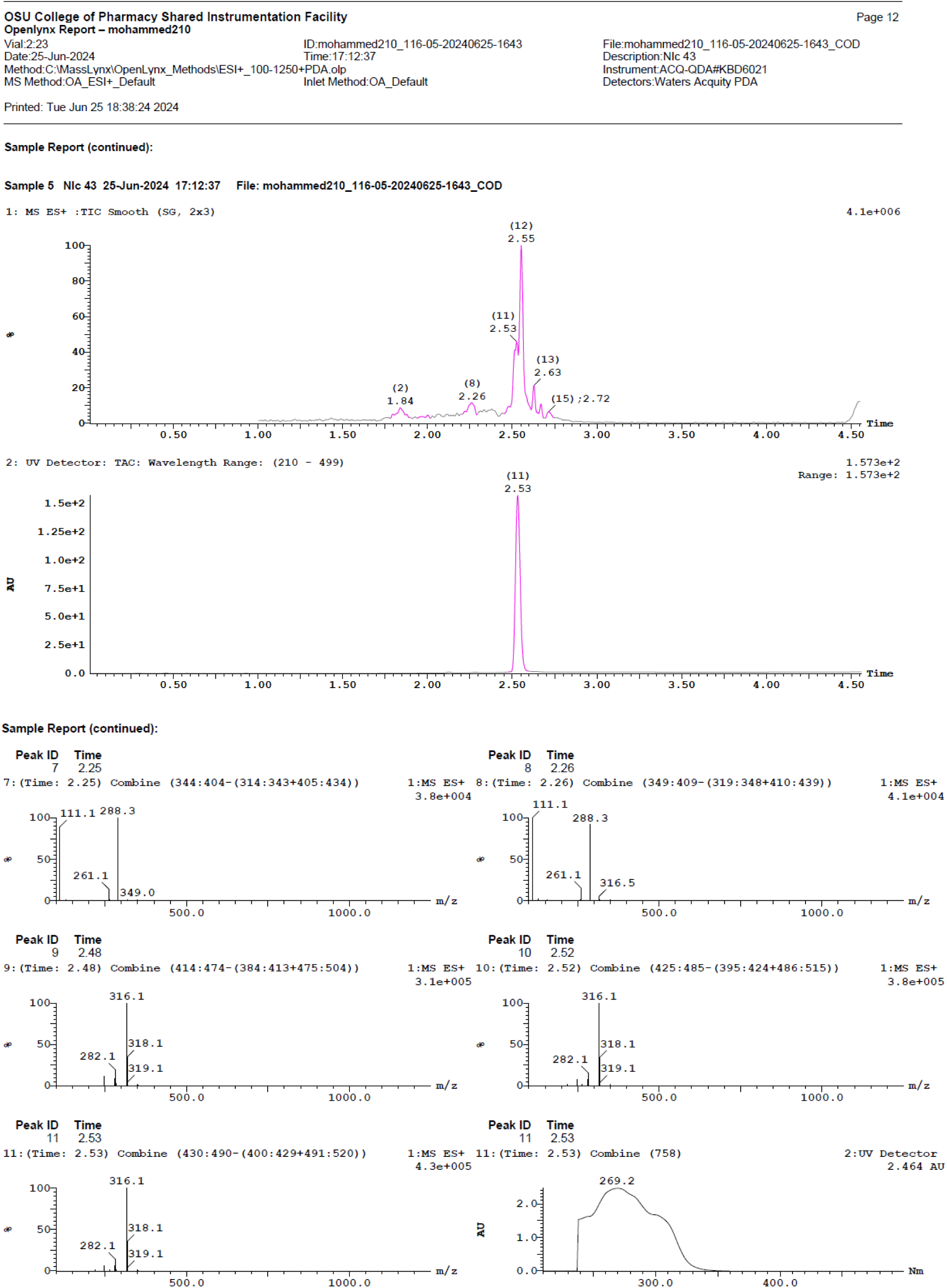

### 2-hydroxy-N-(4-(trifluoromethyl)phenyl)benzamide (Nic-44)

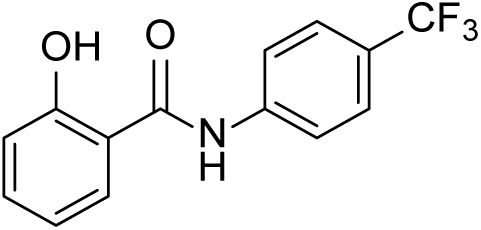

^1^HNMR (400 MHz, DMSO-d_6_) δ 11.5 (s, 1H), 10.63 (s, 1H), 7.96 - 7.90 (m, 3H), 7.73 (d, *J* = 8.5 Hz, 2H), 7.47 - 7.43 (m, 1H), 7.02 - 6.96 (m, 2H); LC-MS (ESI); [M+H]: 282.1

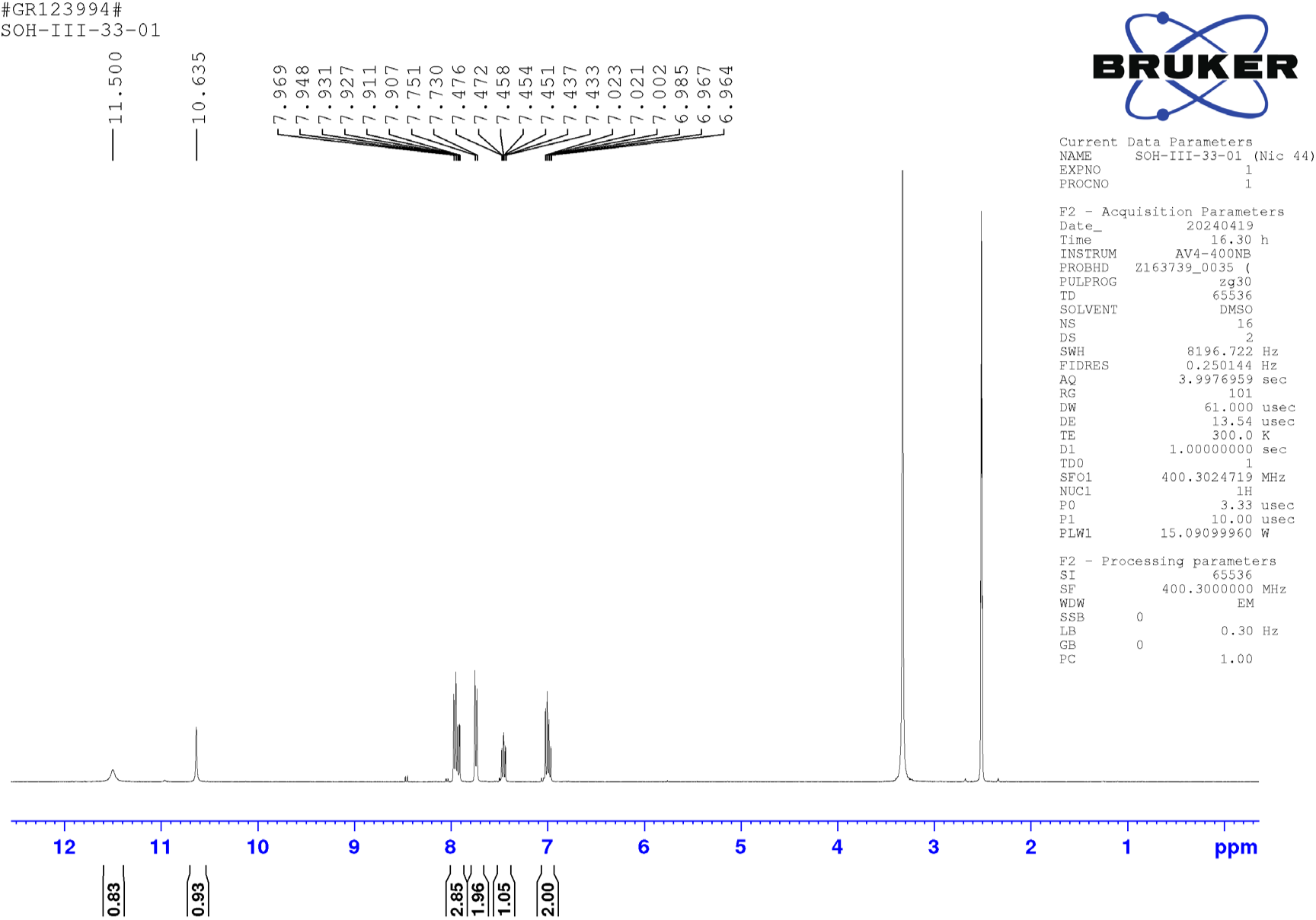

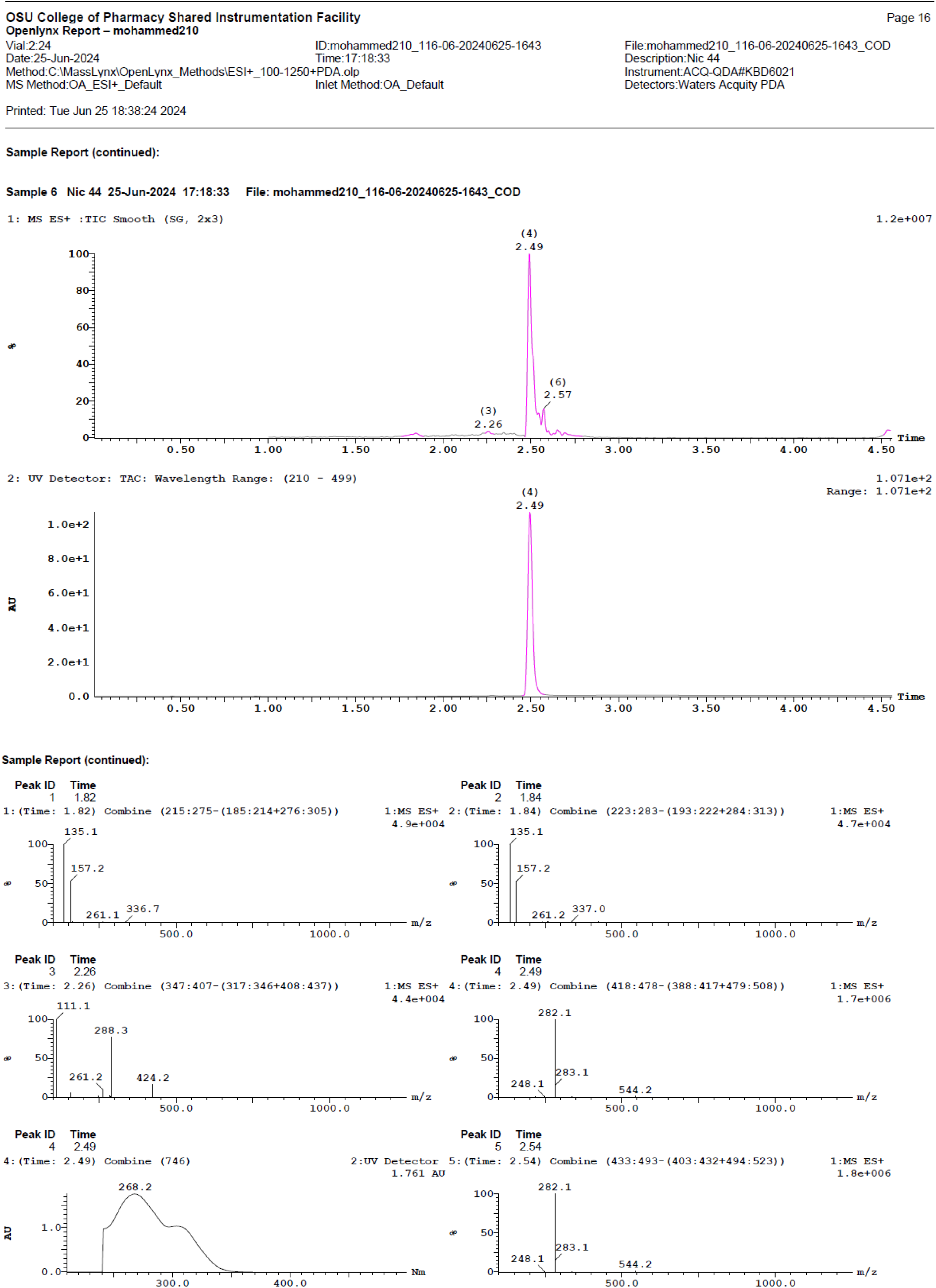

### N-(3,5-bis(trifluoromethyl)phenyl)-2-hydroxybenzamide (Nic-45)

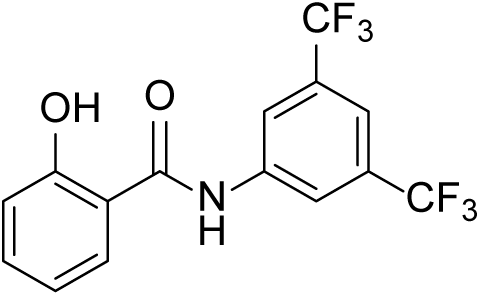

^1^HNMR (400 MHz, CDCl_3_) δ 11.4 (s, 1H), 8.17 - 8.13 (m, 3H), 7.69 (s, 1H), 7.56 - 7.48 (m, 2H), 7.06 (dd, *J* = 8.4 Hz, 0.9 Hz, 1H), 6.98- 6.94 (m, 1H); LC-MS (ESI); [M+H]: 350.1

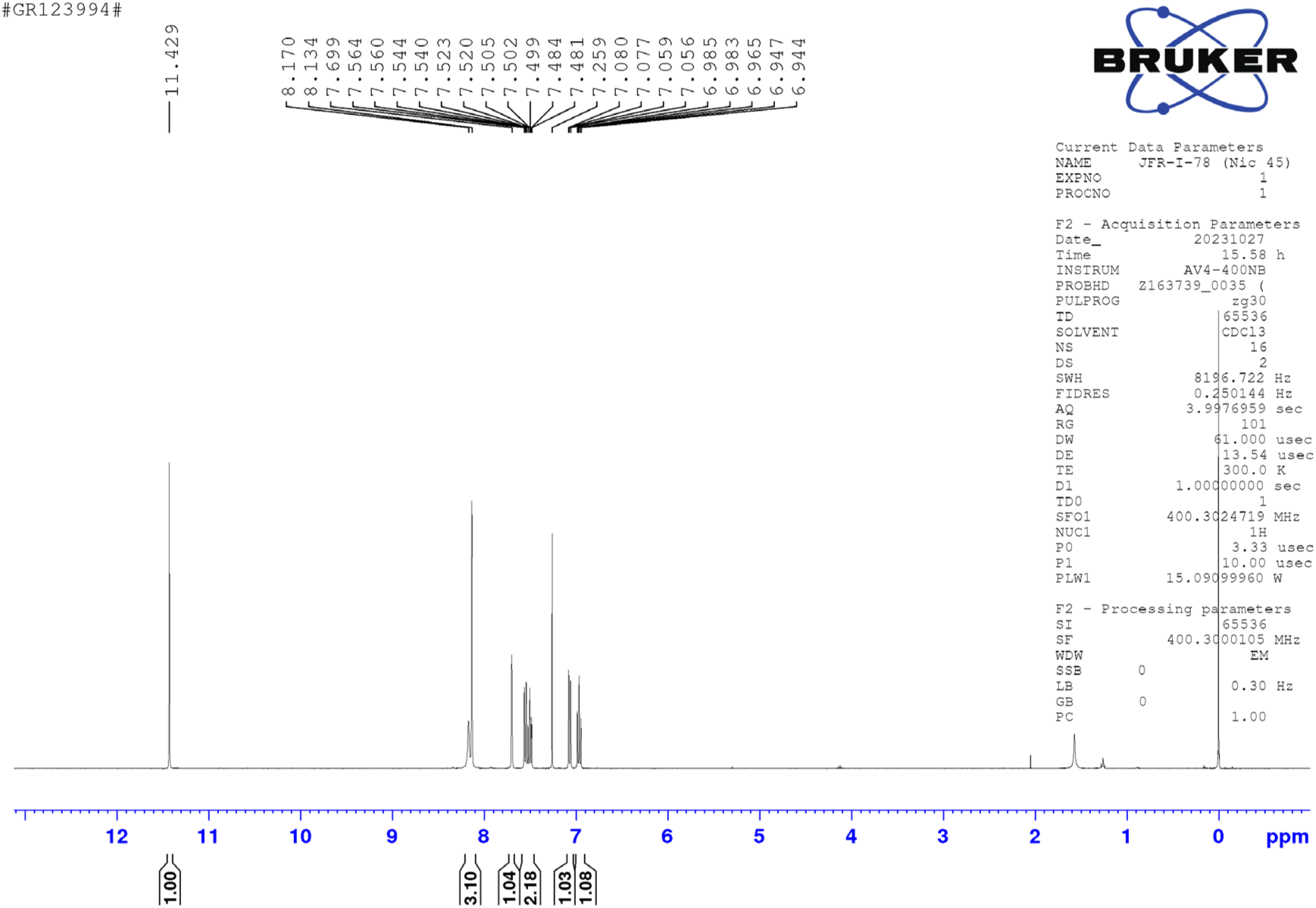

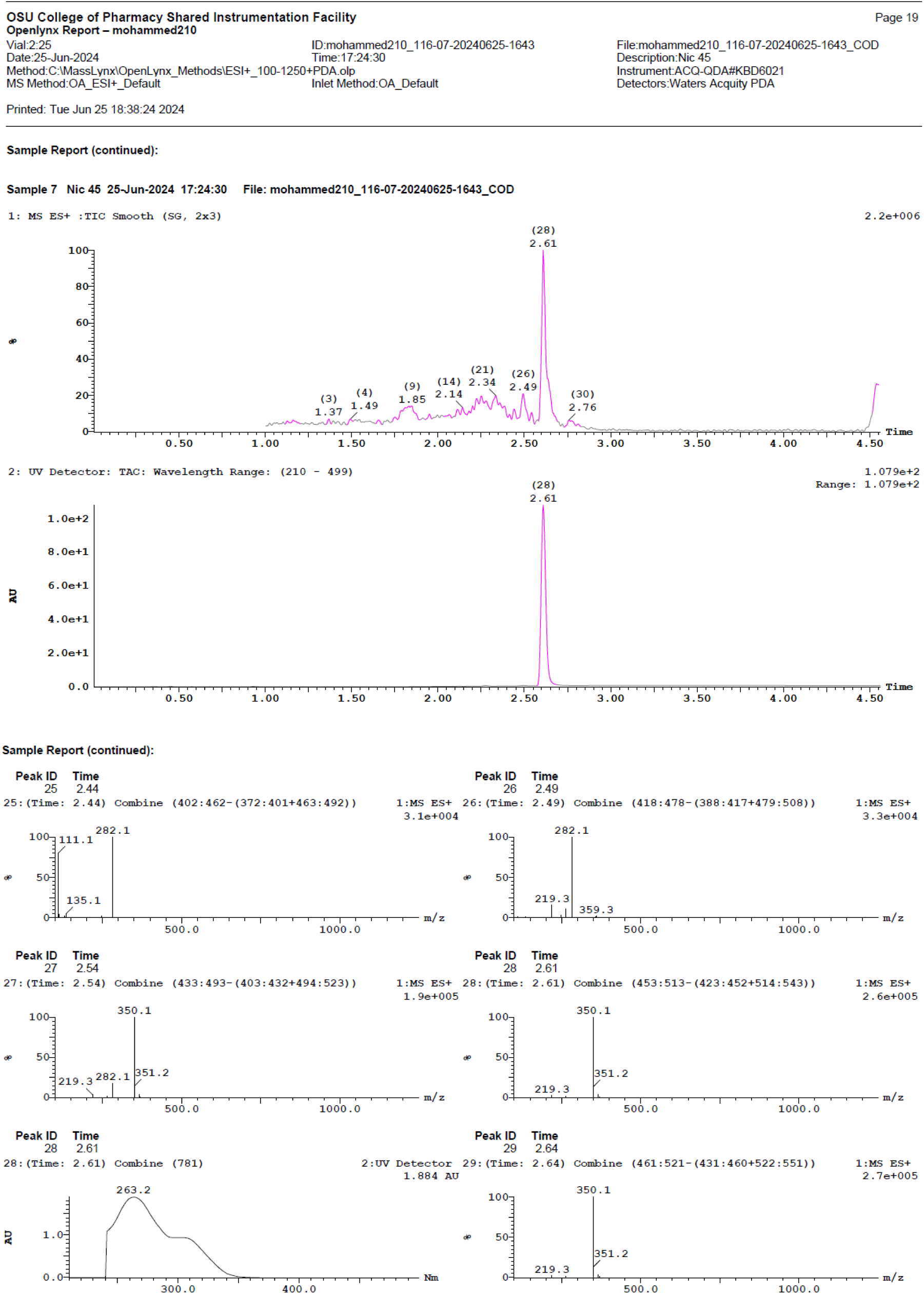

### 2-hydroxy-N-(3-(trifluoromethyl)phenyl)benzamide (Nic-50)

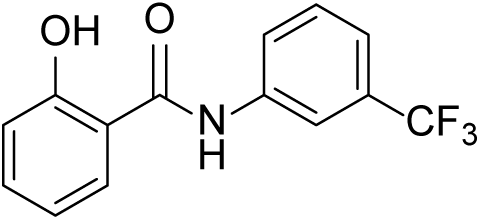

^1^HNMR (400 MHz, DMSO-d_6_) δ 11.51 (s, 1H), 10.60 (s, 1H), 8.22 (s, 1H), 7.96 - 7.91 (m, 2H), 7.62 (t, *J* = 8.0 Hz, 1H), 7.50 - 7.43 (m, 2H), 7.01-6.96 (m, 2H); LC-MS (ESI); [M+H]: 282.1

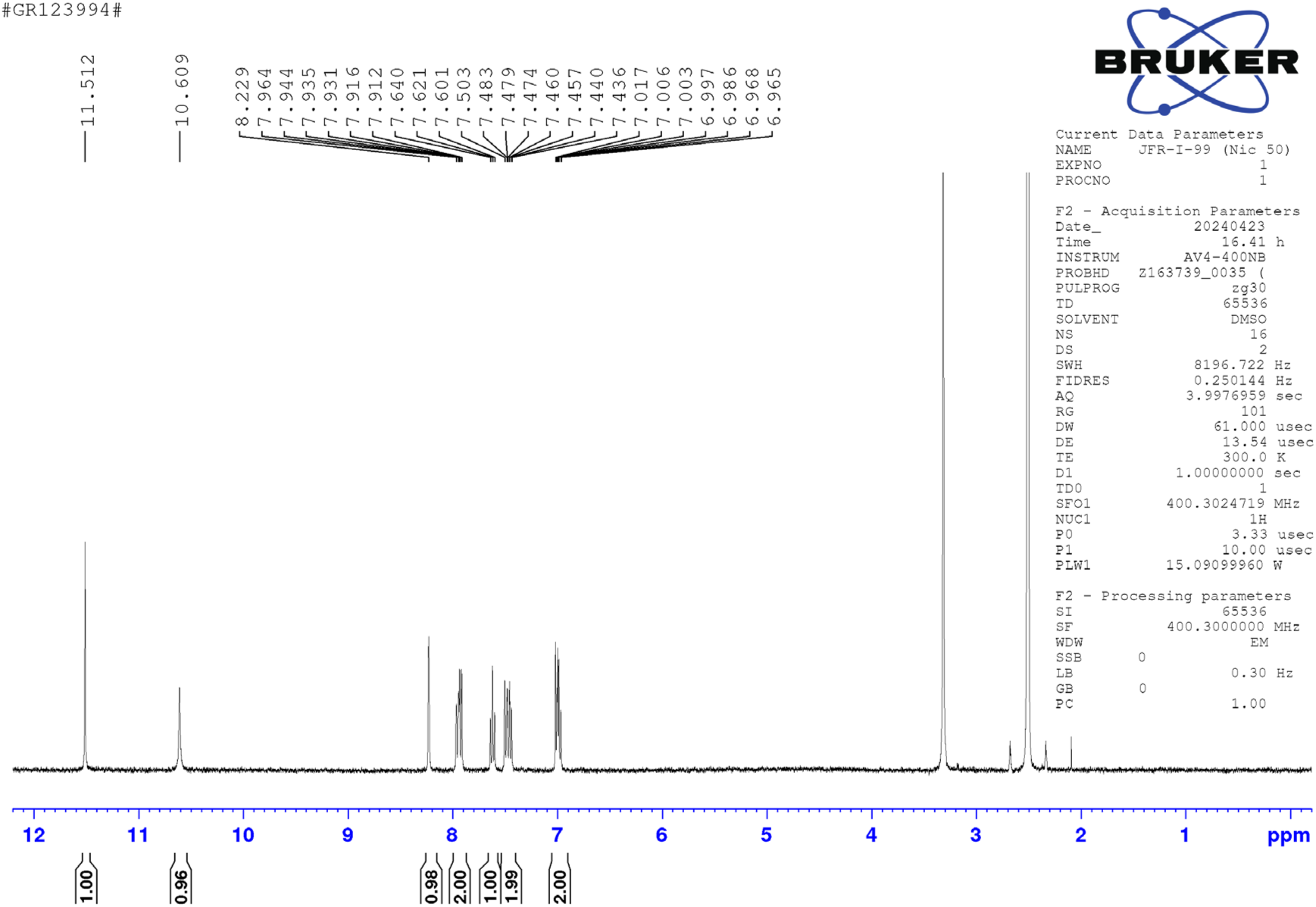

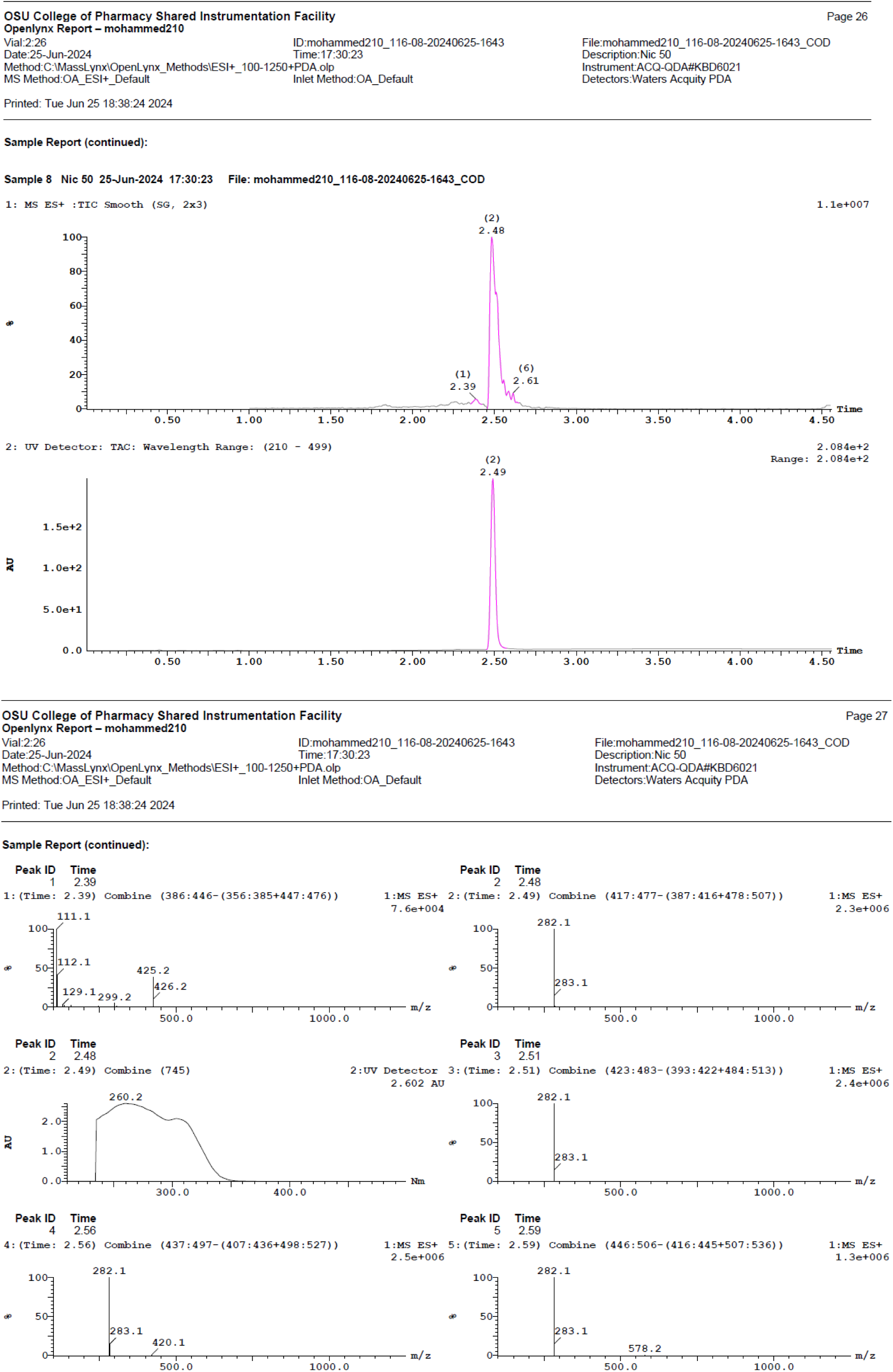

### 5-chloro-2-hydroxy-N-(2-(trifluoromethyl)phenyl)benzamide (Nic-51)

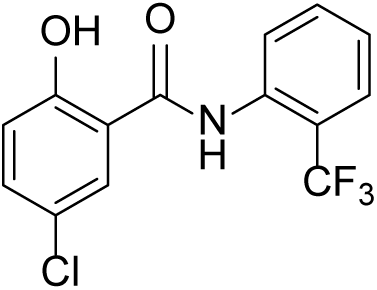

^1^HNMR (400 MHz, DMSO-d_6_) δ 11.57 (s, 1H), 10.65 (s, 1H), 7.94 (d, *J* = 8.4 Hz, 2H), 7.88 (d, *J* = 2.6 Hz, 1H), 7.74 (d, *J* = 8.6 Hz, 2H), 7.47 (dd, *J* = 8.8 Hz, 2.7 Hz, 1H), 7.03 (d, *J* = 8.7 Hz, 1H); LC-MS (ESI); [M+H]: 282.1

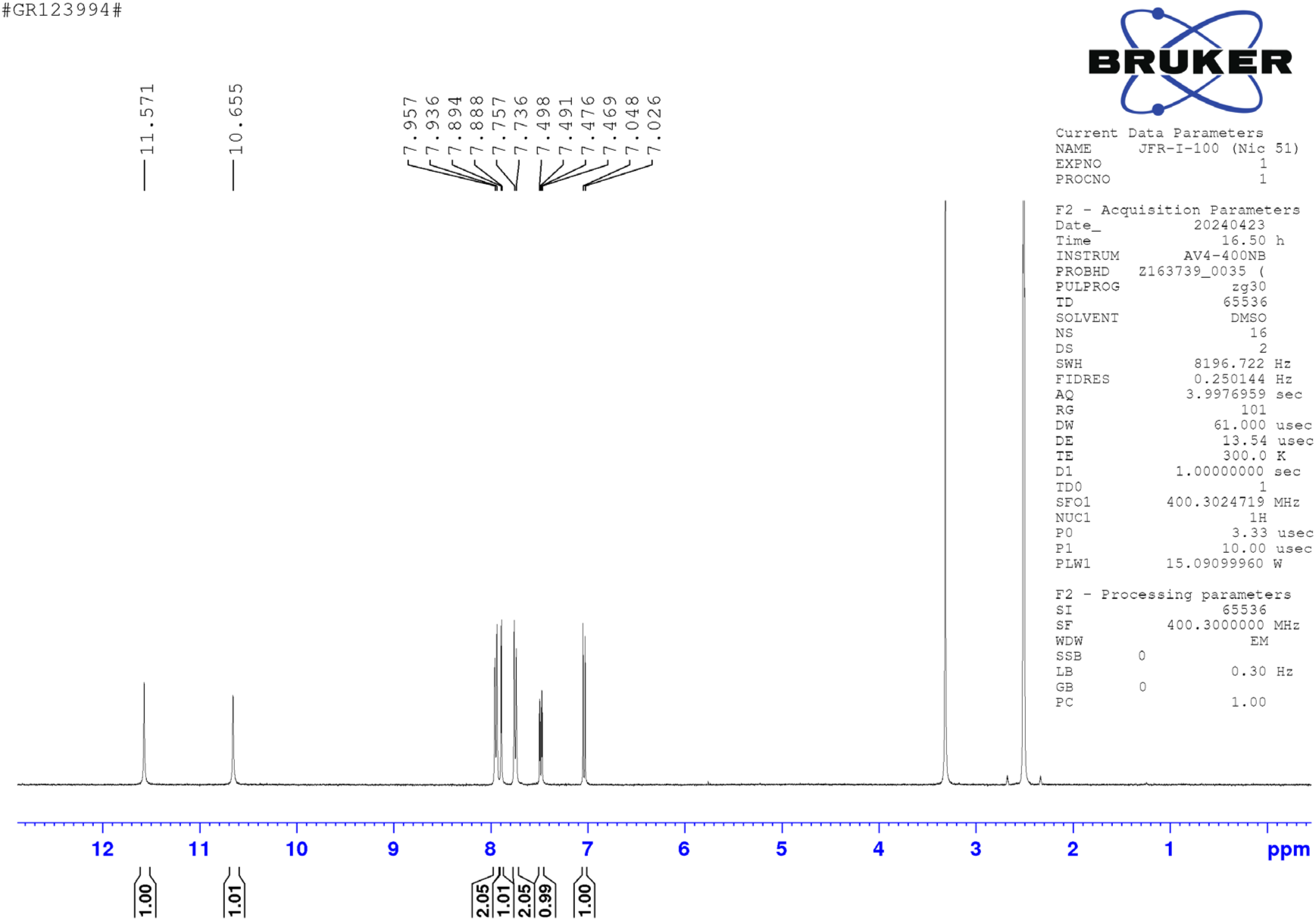

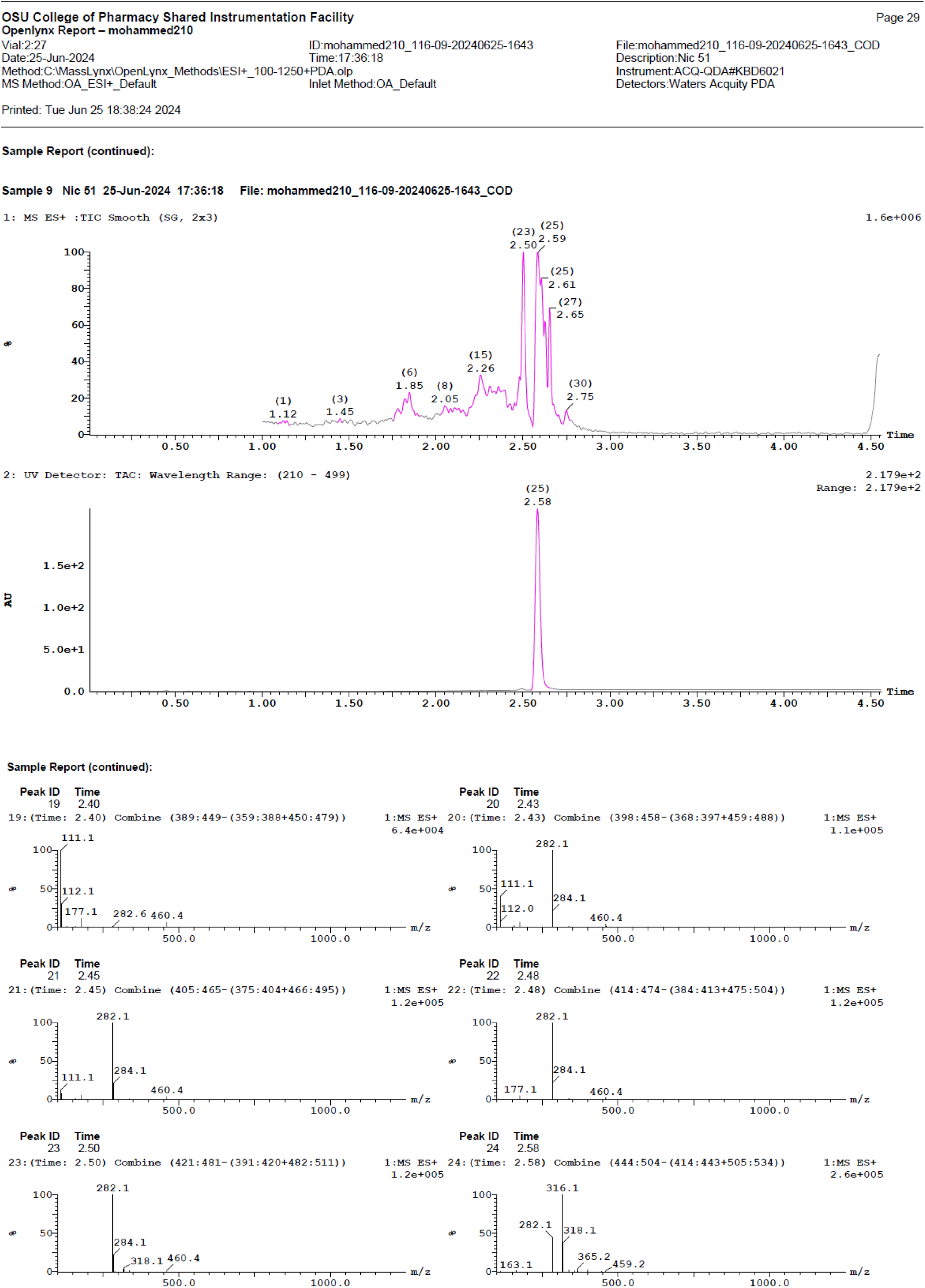

